# Soil microbial inoculants augment fertilizer performance across contrasting cropping systems in Rwanda

**DOI:** 10.64898/2026.08.27.747552

**Authors:** Paige M. Hansen, Anna Edlund, Benjamin Bukumbe, Abhinav Grama, Josue W. Mberwa, Thulani P. Makhalanyane, Janet K. Jansson, Thomas W. Crowther, Jack A. Gilbert

## Abstract

Smallholder farming systems in sub-Saharan Africa are constrained by declining soil fertility, erosion, and rising fertilizer costs, creating an urgent need for scalable inputs that sustain yields while maintaining soil health. While there is some evidence that microbial inoculants may offer a promising complement to conventional fertility management, field-scale evidence in tropical cereal and tuber systems remains limited. Here, we evaluated a multi-species inoculant composed of 20-22 *Bacillus* and *Streptomyces* species on potato and maize across four sites in Rwanda over two growing seasons (2025A and 2025B). Treatments included the inoculant applied at two rates (150 and 250 g ha⁻¹), both alone and in combination with standard fertilization (inorganic fertilizer plus manure), alongside untreated and fertilized controls. Co-application of the inoculant with standard fertilization increased yield and plant biomass beyond fertilization alone, with gains of 6-51% for maize and 3-58% for potato. However, while the inoculant applied alone outperformed untreated controls, it generally did not match standard fertilization. Responses were strongest and most consistent for large-grade potato tubers, and application rate interacted with crop type, whereby the lower dose maximized marketable tuber yield, while maize showed a positive dose-response for grain and biomass. Yield increases were not accompanied by reductions in crop nutrient density, which was instead governed by site-level differences. Altogether, these results indicate that multi-species microbial inoculants are an effective complement to existing fertility practices that may offer, pending further research, a potential pathway to partial fertilizer replacement while sustaining productivity and nutritional quality in smallholder tropical agriculture.

## Introduction

Food production in sub-Saharan Africa is limited by persistent environmental challenges that threaten long-term agricultural sustainability and food security (Luan et al., 2013; van Ittersum et al., 2016). These induce widespread soil nutrient depletion driven by continuous cultivation and accelerated erosion, necessitating the widespread use of inorganic fertilizers that deplete soil health. However, in recent years, rising costs and limited accessibility of these fertilizers have amplified the threats to smallholder farmers (Amankwah et al., 2025; Assefa et al., 2025), limiting crop productivity and resilience across many regions. Taken together, these social and ecological challenges underscore the urgent need for alternative, biologically-based solutions to maintain or improve yields, while restoring soil health rather than depleting it.

Over the past decade, microbial inoculants have emerged as a promising avenue to enhance crop productivity while also improving soil quality, nutrient availability, and biological activity within the soil (Schütz et al., 2018; Just et al., 2024; Li et al., 2024). A growing body of experimental studies and meta-analyses, suggests that microbial inoculants, particularly those composed of multiple, functionally diverse strains can increase crop yields and improve overall plant performance (e.g., Okereke et al., 2000; Adesemoye et al., 2009; Bashan et al., 2014; Jiang et al., 2023; Liu et al., 2023; Francioli et al., 2025; Bukombe et al., 2026). However, much of this evidence has traditionally been generated under controlled greenhouse conditions, focusing heavily on temperate agroecosystems, which limit its generalizability to tropical smallholder farms that face the most pressing sustainability challenges (Averill et al., 2022). In response, recent field studies have begun to demonstrate strong benefits of multi-species inoculants in enhancing the productivity of cereal crops and legumes in Rwanda (e.g., Masso et al., 2016; Koskey et al., 2017; Nyaguthii, 2017; Bukombe et al., 2026). However, it remains unclear whether these effects can partially or totally offset the need for standard inorganic fertilizers, or whether they have other long-term impacts on soil health and crop nutrient quality. Critically, whether inoculants can substitute for fertilizer, either fully or partially, or instead complement it implies different management strategies for smallholders. Yet, field evidence distinguishing these possibilities remains scarce.

A central consideration for any soil amendments is the potential trade-off between crop yield and nutrient quality. This concern is grounded in context-specific findings of nutrient dilution within harvested plant tissues, whereby increases in yield can be accompanied by decreases in nutrient concentrations within plant tissues, depending on crop type and local soil properties and nutrient concentrations (Buerkert et al., 1998; Feil et al., 2005; Cakmak, 2008; Davis, 2009; Zhao et al., 2020). This potential trade-off between yield and plant nutrient density represents a potential limitation for existing synthetic fertilizers, particularly in regions where micronutrient deficiencies are already prevalent, as in many parts of East Africa (Joy et al., 2014; Galani et al., 2022). This issue is further compounded by evidence that rising atmospheric CO_2_ concentrations may already be contributing to declines in nutrient density across a range of growing environments (Myers et al., 2014). However, it remains unclear whether these trade-offs are also apparent following microbial enrichment.

Here, we used a series of field experiments across smallholder farming systems in Rwanda to assess the potential of microbial inoculation to complement the need for standard fertilizer application, and to evaluate the potential trade-off between crop yield and quality. Specifically, we quantified the effects of an inoculant composed of 20-22 *Bacillus* and *Streptomyces* bacterial species (Table S1; Bukombe et al., 2026), on the yields of potato and maize grown in tropical agroecosystems across Rwanda. We hypothesized that microbial inoculation would increase yield and plant biomass relative to untreated controls as well as standard fertilization practices (i.e., inorganic fertilizer and manure co-application). We also tested whether any such yield gains were accompanied by nutrient dilution, a trade-off reported where yields are increased through fertilization or breeding (Davis, 2009; Guo et al., 2020). Because inorganic fertilizer supplies immediately available nutrients while microbial inoculants can enhance nutrient solubilization, acquisition, and uptake efficiency, we reasoned that the two inputs act on complementary limitations. With this, we predicted that the greatest increases in yield and biomass for both crops would occur under combined application of the inoculant and fertilizer, consistent with additive contribution from fertilizer-supplied and microbially mediated nutrient availability. Additionally, we tested whether treatment effects varied across crop type, soil fertility status, and season, expecting the largest relative effects in lower-fertility soils, where marginal benefits of any yield-enhancing input is typically greatest. .

## Materials and Methods

### Taxonomic classification and formulation of microbial inoculant

To capture broad functional diversity, we assembled our microbial inoculant from a diverse culture collection of beneficial soil microorganisms, including spore-forming *Streptomyces* and *Bacilli* bacteria, as described in Bukombe et al (2026). Standardized production and formulation workflows were applied, including separate bacterial propagation, quality control of viable cells and spores, and final blending into our multi-organism inoculant (Table S1).

Genomic sequences used for the identification of bacterial strains derived from Oath Inc.’s collection (Edlund et al., 2026) were obtained through whole-genome sequencing using either Illumina paired-end or Oxford Nanopore sequencing. Following Edlund et al., Illumina paired-end reads were preprocessed using fastp (v1.3.6; Chen et al., 2018), then assembled using SPAdes (v3.15.5; Bankevich et al., 2012) through VEBA (v2.5.3; Espinoza and Dupont, 2022; Espinoza et al., 2024). Oxford Nanopore reads were preprocessed using Fastplong (v0.7.0; Chen, 2023), assembled with Flye (v2.9.3; Wick et al., 2021), and polished using Medaka v2.2.2 implemented via VEBA. Each assembly met high-quality genome standards via CheckM2 (v1.1.0; Chklovski et al., 2023; Sorensen et al., 2026). All taxonomy was classified via GTDB-Tk (v2.6.1; release 226; Chaumeil et al., 2022).

All bacterial components were produced using agar growth surface fermentation as described in Bukombe et al., 2026 and Edlund et al., 2026, including liquid and agar media preparation, seed culture generation, plate-scale biomass production, harvesting biomass from the top of the plates, drying of biomass, mixing with dextrose sugar to provide an osmoprotectant and a mixing agent, and grinding dried biomass into a powder, as well as quality control that included visual inspection of growing bacteria on agar plates at multiple timepoints during incubation in 28°C. Any cultures that failed purity or growth criteria were excluded from downstream mixing. For each strain, 1 g replicate samples were shipped overnight to BioForm Solutions Inc. (San Diego, CA) for quantification of live cells and spore counts via flow cytometry, using the standard protocol ISO 19344 Part B. To formulate the final inoculant applied in the field, bacterial spores were combined and diluted with dextrose to a final viable count of ≥10⁷ CFU g⁻¹, and the wettable powder was packaged in non-transparent, sealable bags (Edlund et al., 2026).

Microbial inoculant formulations differed slightly between the two study seasons. Specifically, the inoculant applied in the 2025B season included two additional *Bacillus* strains relative to the formulation used in 2025A (Table S1), resulting in a total of 22 strains compared to 20 in the earlier formulation, published in Bukombe et al., 2026. Aside from these additions, both formulations were otherwise identical in composition and production procedures. Both products were approved for field application by the Rwanda Institute for Conservation Agriculture (RICA), with the 2025B formulation authorized as an amendment to the original 2025A product.

### Experimental design and set-up

Field trial establishment and execution was carried out by the Rwanda Agriculture and Animal Resources Development Board (RAB) and conducted without involvement of the parent organization for the inoculum (Oath, Inc.); sharing and publication of all data collected throughout this project is approved by RAB. Field trials were established in four districts (i.e., Nyamagabe, Musanze, Kayonza, and Ngoma) representing contrasting agroecological zones for each crop type, reflecting typical potato and maize cultivation regions in Rwanda (Figure 1, Table 1). For potato (*Solanum tuberosum*; early-mature Kinigi variety), sites were in the Congo-Nile Divide landscape region within Nyamagabe District (-2.5023, 29.4458) and in the Virunga zone within Musanze District (-1.4731, 29.6351). For maize (*Zea mays*; RHM1407 hybrid), sites were in the Eastern Plateau within Kayonza District (-1.8364, 30.4443) and in the Bugesera zone within Ngoma District (-2.0897, 30.4039). Musanze and Kayonza were selected to represent the more fertile soil types for potato and maize sites, respectively, as reflected in our analysis of soil properties across all four regions (Table 1). The four sites also differ in climate characteristics, with Kayonza and Musanze having a 20-year mean annual temperature (MAT) of 20.6°C and a 20-year mean annual precipitation (MAP) of 1285.5 mm, and Nyamagabe and Musanze having a MAT of 18.9°C and 17.3°C and a MAP of 1461 mm and 1726.4 mm, respectively (Sparks, 2018; NASA Langley Research Center (LaRC) POWER Project).

**Figure 1.**
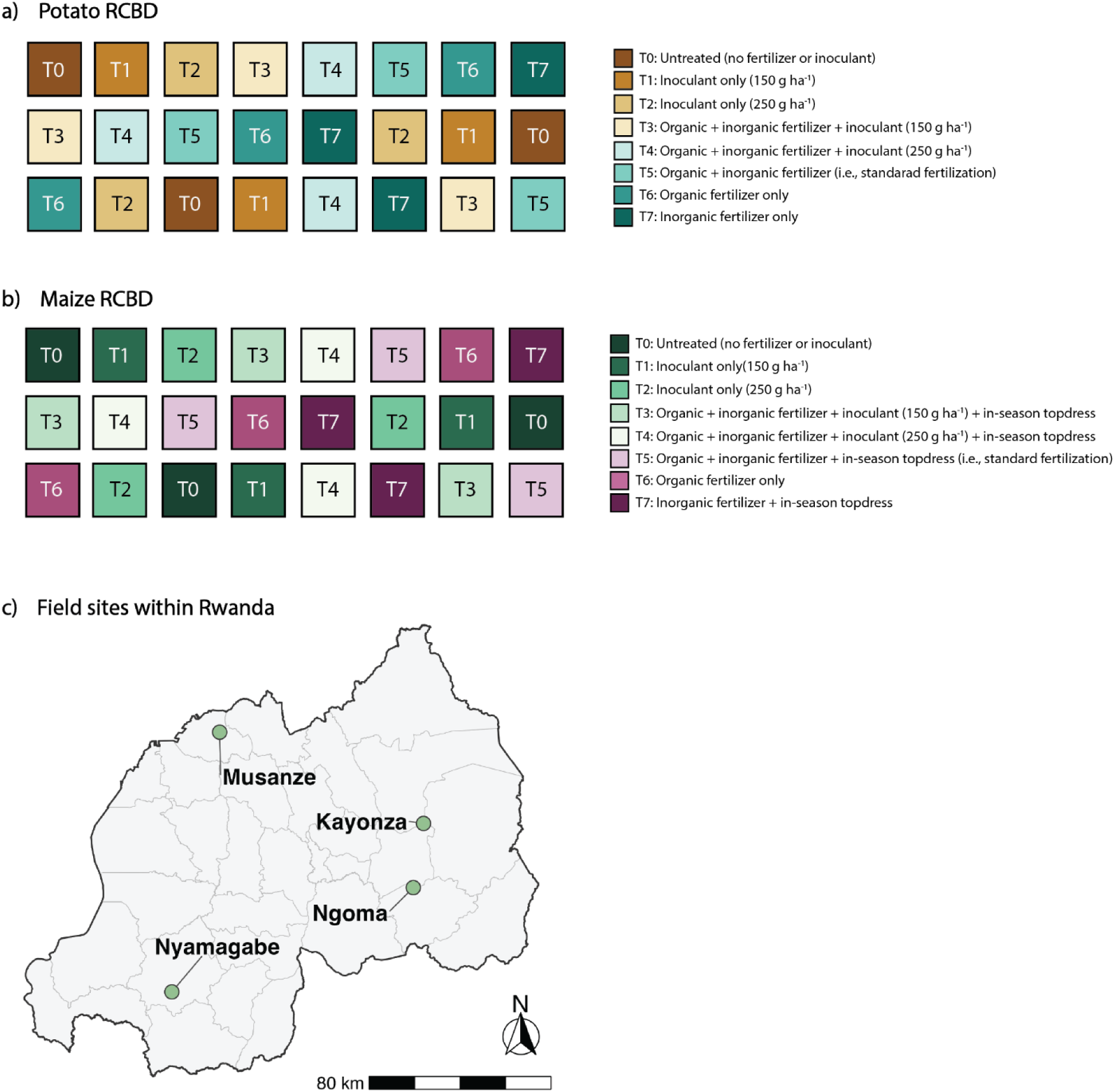
Randomized complete block design (RCBD) for a) both Musanze and Nyamagabe sites designated for potato experiments, and b) both Kayonza and Ngoma sites designated for maize. In both panels, colors indicate each of eight treatments applied to test the efficacy of the microbial inoculant. Untreated treatments (i.e., T0) as well as those involving application of fertilizers only (i.e., T5-7) serve as controls, with co-application of inorganic and organic fertilizers (T7) representing standard fertilization practice across both crops and all regions. Panel c) shows site locations across Rwanda.

**Table 1.** Site-averaged soil physicochemical properties (mean ± standard error). The “Significance” column indicates corrected p-values and significance levels of Welch’s t-test results.

|  | Potato trial sites |  |  | Maize trial sites |  |  |
| --- | --- | --- | --- | --- | --- | --- |
|  | Musanze | Nyamagabe | Significance | Ngoma | Kayonza | Significance |
| Soil pH | 5.2 ± 0.02 | 4.6 ± 0.04 | $p=3.3\text{e-}16$ *** | 5.0 ± 0.1 | 5.6 ± 0.1 | $p=5.2\text{e-}9$ *** |
| **CEC (cmol kg soil<sup>-1</sup>)** | 41.2 ± 0.4 | 11.8 ± 0.2 | $p=3.3\text{e-}42$ *** | 10.8 ± 0.5 | 13.5 ± 0.3 | $p=6.5\text{e-}5$ *** |
| **Sand (%)** | 52.2 ± 0.8 | 54.1 ± 1.6 | $p=0.29$ n.s. | 58.3 ± 0.7 | 61.1 ± 0.6 | $p=0.006$ ** |
| **Silt (%)** | 36.2 ± 0.8 | 20.8 ± 0.6 | $p=1.3\text{e-}21$ *** | 14.6 ± 0.5 | 11.9 ± 0.2 | $p=7.6\text{e-}6$ *** |
| **Clay (%)** | 11.5 ± 0.2 | 25.1 ± 1.1 | $p=2.6\text{e-}13$ *** | 27.1 ± 0.7 | 27.0 ± 0.7 | $p=0.95$ n.s. |
| **Total N (g kg soil<sup>-1</sup>)** | 7.9 ± 0.1 | 2.4 ± 0.1 | $p=8.3\text{e-}44$ *** | 1.6 ± 0.1 | 1.4 ± 0.04 | $p=0.06$ · |
| **SOC (g kg soil<sup>-1</sup>)** | 37.9 ± 0.6 | 24.7 ± 0.6 | $p=9.7\text{e-}23$ *** | 15.4 ± 0.3 | 19.6 ± 0.5 | $p=2.4\text{e-}10$ *** |
| **Total P (g kg soil<sup>-1</sup>)** | 3.6 ± 0.1 | 1.0 ± 0.04 | $p=1.4\text{e-}6$ *** | 0.7 ± 0.01 | 0.8 ± 0.2 | $p=0.002$ ** |
| **Available P (g kg soil<sup>-1</sup>)** | 9.1 ± 0.5 | 16.1 ± 1.1 | $p=1.5\text{e-}28$ *** | 9.0 ± 0.8 | 45.8 ± 5.9 | $p=1.1\text{e-}6$ *** |

CEC: Cation exchange capacity; N: Nitrogen; SOC: Soil organic carbon; P: Phosphorus

Experiments at each of the four sites (hereafter referred to by district name) were conducted using a randomized complete block design (RCBD) with three replicates and eight treatments during seasons 2025A (September 2024 to February 2025) and 2025B (February 2025 to July 2025; Figure 1). Outside of the inoculant and fertilization treatments described below, all sites were rain-fed and did not receive any irrigation water in either season. Season 2025A was wetter and slightly warmer than 2025B, with all four sites receiving 254 to 476 mm more rainfall and being an average of 0.1 to 0.5°C warmer in 2025A than in 2025B (Figure S1; Sparks, 2018; NASA Langley Research Center (LaRC) POWER Project).

As described above, the inoculant formulated for season 2025B included two additional strains compared to season 2025A (Table S1). All plots (25 m^2^) were separated by 1 m in potato and 2 m in maize experiments to minimize the risk of contamination. Each experiment in both seasons included eight treatments: one untreated control; two inoculant-only treatments that differed in application rate (i.e., 150 g ha^−1^and 250 g ha^−1^); one organic fertilizer-only control; one inorganic fertilizer-only control only; one combined organic and inorganic fertilizer control, representing fertilization practices across all four sites; and two treatments that included standard fertilization supplemented with either the low or high inoculant dose. In potato experiments, organic fertilizer consisted of 10 t ha^−1^ manure, and inorganic fertilizer consisted of 300 kg ha^−1^ NPK. In maize experiments, organic fertilizer also consisted of 10 t ha^−1^ manure, while inorganic fertilizer consisted of 100 kg ha⁻¹ diammonium phosphate (DAP). Additionally, all maize treatments receiving inorganic fertilizer also received 100 kg ha^−1^ urea fertilizer as topdress.

Potato experiments were established identically in both seasons. Organic manure and inorganic NPK fertilizer were applied in furrows along the planting lines and covered with a thin layer of soil before planting potato tubers at a rate of 2.5 t ha⁻¹. The microbial inoculant was diluted at 0.375 g in 5 L of water for the 150 g ha⁻¹ rate and at 0.625 g in 5 L for the 250 g ha⁻¹ rate, with each solution applied to 25 m² plots. At planting, the inoculant was applied twice: once directly to the seed tubers, and again after covering them with soil. Following planting, the inoculant was applied every 14 days until tuber hardening, for a total of five applications during the potato cropping cycle. While the same quantities of water were not applied to non-inoculated plots, the total of 25 L of water applied over the course of each growing season translates to only 1-1.2 additional mm of water, approximately five to ten times less than the standard deviation of daily precipitation estimates (Sparks, 2018; NASA Langley Research Center (LaRC) POWER Project).

Maize experiments were established similarly. Organic manure and inorganic DAP were applied in furrows along the planting lines and covered with a thin soil layer prior to sowing maize seed at a rate of 30 kg ha⁻¹. This variety is adapted to low- and medium-altitude regions in Rwanda (900–1700 m above sea level) and has a yield potential of 7.87 to 13.5 t ha⁻¹.

Inoculant application followed the same dilution and application procedure as in potato. At planting, inoculant was applied twice (once directly to the seeds and again after soil coverage). Subsequent applications were made every 14 days until the grain-filling stage, resulting in six applications over the maize cropping cycle.

### In-field data and sample collection

Data collection from potato experiments focused on small and large tuber yield, as well as plant biomass. Tubers and plants were collected at harvest from a net plot size of 14.92 m^2^, delineated by leaving out one row of plants approximately 30 cm from each plot edge to ensure that all data collected was not affected by treatment contamination from neighboring plots. To quantify fresh plant biomass for each net plot, full potato plants, including shoots and tubers, were uprooted, shaken to remove soil material, and weighed for each treatment and replicate. For marketable yield, tubers were separated into large and small size classes, weighed, then mixed to measure total yield.

Like the potato experiments, maize data collected focused on grain yield and plant biomass. Maize plants were harvested from a net plot size of 16 m^2^, delineated by excluding one border row of plants from each plot edge, when at least 97% of individual plants were dried to 15-20% moisture. Maize kernels were shelled from cobs by hand to prevent grain breakage and loss, separated from impurities by winnowing, and were then immediately weighed in the field to measure yield. Plants were cut at ground level, bundled together, and weighed to measure fresh aboveground biomass. As with potato harvest, a 1-kg subsample of grain yield was taken to measure maize nutrient density.

At the time of harvest, soil samples were also collected to assess across- and within-site contrasts in soil physicochemical properties that have the potential to impact yield and biomass. Soils were collected by taking five subsamples from each plot to a depth of 20 cm in a Y-shape, then combining and thoroughly homogenizing them to create one composite sample per plot.

Both plant tissues reserved for nutrient density analyses and soil samples were immediately transported to the Rwanda Agriculture and Animal Resources Board (RAB) laboratory (Rubona, Huye District, Rwanda) for further processing.

### Laboratory analysis of soil physicochemical properties

After arrival at the RAB laboratory, all soil samples were air-dried, homogenised, sieved to 2 mm, and subsampled for measurement of physical and chemical properties, including texture, pH, cation exchange capacity (CEC), soil organic carbon (SOC), total nitrogen (TN), total phosphorus (P), and available P.

Soil texture was analysed using the Bouyoucos hydrometer method (Bouyoucos, 1962), with modifications following Beretta et al., 2014. Briefly, 50 g of air-dried, 2 mm-sieved soil were dispersed with 10% sodium hexametaphosphate (NaPO_3_) and treated three times with 6% hydrogen peroxide (H_2_O_2_) at 60°C to remove organic C. After mixing the soil suspension and transferring it into a clean glass column, hydrometer readings on suspension density were taken after 40 s and 2 after hrs to distinguish between the silt and clay fractions, respectively.

Soil pH was measured potentiometrically using a glass electrode connected to a portable multiparameter meter (HI9828, Hanna Instruments US Inc., USA) following Black, 1965, where 20 g of air-dried, 2-mm sieved soil was suspended in 1M H_2_O at a 1:2.5 soil:solution ratio, stirred for 10 min, and allowed to equilibrate for 30 min before pH measurement.

Cation exchange capacity was quantified with atomic absorption spectrometry (AAS). Air-dried, sieved soils were extracted with an excess of 1 M ammonium acetate (NH_4_OAc), such that the maximum exchange occurs between the NH_4_ and the cations originally occupying exchange sites on the soil surface (Schollenberger and Simon, 1945). Total exchangeable cations (i.e., CEC) in the extract were then determined by AAS.

Soil organic carbon (SOC) was measured using a modified Walkley-Black procedure following Nelson and Sommers (1974), from a sulphuric acid (H_2_SO_4_) and aqueous potassium dichromate (K_2_Cr_2_O_7_) mixture after complete oxidation of the solution via external heating. The unused or residual K_2_Cr_2_O_7_ after oxidation was titrated against ferrous ammonium sulphate (Mohr’s salt). The amount of “used” K_2_Cr_2_O_7_, or the difference between added and residual K_2_Cr_2_O_7_, gives a measure of the SOC content of the soil (Nelson and Sommers, 1974).

Total nitrogen (N) and total phosphorus (P) concentrations were determined via wet digestion of soil samples using a mixture of H_2_SO_4_, H_2_O_2_, selenium (Se), and salicylic acid (C_7_H_6_O_3_; (Bremner and Mulvaney, 1982; Walinga et al., 1995). To prevent nitrate-N loss, C_7_H_6_O_3_ was employed in an acidic medium to facilitate the formation of nitro-salicylic compounds, which were subsequently reduced to amino forms by soil organic matter. Complete oxidation of organic materials was achieved through H_2_O_2_ treatment, with Se serving as a catalyst and H_2_SO_4_ completing the digestion under high temperature.

Given that soil pH was less than 7 for all sampled sites, available soil P was extracted using the Bray and Kurtz No. 1 method (Bray and Kurtz, 1945). Briefly, 3 g of air-dried, 2-mm sieved soil were shaken with 30 mL of Bray-1 extractant (0.03 M NH_4_F + 0.025 M HCl) for 5 min on a reciprocating shaker, then filtered through Whatman No. 42 paper. Available P concentration in the filtered extract was then determined colorimetrically by the molybdenum blue method at 884 nm using a UV-VIS spectrophotometer.

### Laboratory analysis of plant nutrient density

After receipt at the RAB laboratory, fresh plant material for nutrient density analyses was oven-dried at 65°C, ground to a fine powder using a mill, and then passed through a 0.5 mm sieve. Iron (Fe) concentration was measured on all potato plant tissues, while crude protein, calcium (Ca), potassium (K), phosphorus (P), and organic carbon (OC) content was measured on all maize plant tissues. Nutrient panels differed by crop to reflect crop-specific research priorities, with potato serving as a target for iron biofortification and maize more commonly evaluated for broader aspects of grain nutrient composition.

Iron concentration in potato plant tissue was determined following Kalra (1998). Approximately 0.5 g of ground, oven-dried plant material was digested in a block using a mixture of concentrated nitric acid (HNO_3_) and H_2_O_2_ until all organic matter was completely oxidized. The resulting digest was cooled, made up to a known volume with deionised water, and filtered through Whatman No. 42 paper. Flame AAS at 248.3 nm was then used to determine Fe concentrations from the filtered digest.

Crude protein in maize plant tissue was measured using the Kjeldahl method (Thiex et al., 2002). Briefly, 0.5 g of ground, oven-dried plant material was digested in concentrated H_2_SO_4_ with a copper catalyst and potassium sulfate (K_2_SO_4_) at 420°C until the digest ran clear. After cooling, the digest was diluted and made alkaline with concentrated sodium hydroxide (NaOH), and NH4 was released by steam distillation into a boric acid (H3BO3) receiving solution. The resulting NH4-B complex was titrated with standardized hydrochloric acid (HCl) to determine total N content. Crude protein was calculated by multiplying percent N concentrations by a conversion factor of 6.25.

Mineral concentrations (i.e., Ca, K, and P) in maize tissue were determined following the same Kalra (1998) used for measuring Fe concentrations in potato tissues. After plant material was completely digested in HNO_3_ and H_2_O_2_ and filtered, Ca concentrations were determined by flame AAS at 422.7 nm, K concentrations were determined by flame emission spectrophotometry at 766.5 nm, and P was quantified colorimetrically using the vanadomolybdate method at 400 nm with a UV-VIS spectrophotometer.

Organic C content of maize tissue was measured using the same Walkley-Black wet oxidation procedure applied to soil samples (Nelson and Sommers, 1974), adapted for plant material. Approximately 0.1 g of ground, oven-dried material was oxidised with H_2_SO_4_ and aqueous K_2_Cr_2_O_7_ under external heating, and the residual K_2_Cr_2_O_7_ was back-titrated against ferrous ammonium sulphate to quantify maize plant OC.

### Statistical analyses

We first evaluated whether differences in soil physicochemical properties between sites matched expectations (i.e., Musanze and Kayonza are generally more fertile than Nyamagabe and Ngoma, respectively; Table 1) using Welch’s t-tests, with *p*-value corrections for multiple comparisons.

Next, to assess the effects of inoculation on yield and biomass, we fitted ordinary least-squares linear models within defined strata of crop type, site, and season, depending on the contrast in question. For instance, models assessing the effects of treatment and site were stratified between crop type and season, while models assessing treatment and season were stratified between crop type and site. Though soil physicochemical properties are known to significantly regulate yield and plant biomass, the soil variables measured in this study were not included as covariates in our linear models because their variation largely aligned with site-level differences (Table 1; Figures S2-3; Table S2-3). All models used sum-to-zero contrasts with Type III ANOVA to test for main effects of 1) treatment and site; and 2) treatment and season, as well as their interactions. Given that the interactive effects of treatment and site or season were significant in nearly all models, we intentionally retained the interactive effect in place of incorporating site or season as random effects. When omnibus tests were significant (*α*=0.05), post-hoc pairwise comparisons of estimated marginal means were conducted with false discovery rate (FDR) adjustment. Although all pairwise contrasts were evaluated, emphasis was placed on comparisons between untreated controls, inoculant-only treatments, controls representing the standard fertilization practice for the region (i.e., both organic and inorganic fertilizers), and co-application of the inoculant and standard fertilizers to determine agronomic relevance. For visualization, site- and season-specific yield responses were expressed as percent change relative to the regional standard, with replicate means and 95% confidence intervals (*t* distribution, *n*=3).

To evaluate potential trade-offs between yield gains and nutrient density, identical linear modeling and post-hoc testing procedures were applied to nutrient metrics. Post-harvest soil data were also analyzed using the same framework, to compare within- and across-site soil heterogeneity.

All analyses were conducted in R (version 4.4.1; R Core Team, 2024) using a combination of car (Fox et al., 2026), emmeans (Lenth et al., 2026), and tidyverse (Wickham et al., 2019).

## Results

### Potato tuber yield and plant biomass

Across both growing seasons, all measured potato yield metrics varied significantly across treatments and sites (Figure 2; Tables S4-7). Yield and biomass were also shaped by seasonal variability across all districts, with interactions between season and treatment observed for total, large-grade, and small-grade tuber yield in Musanze (all *p*<0.001), as well as for plant biomass in both Musanze and Nyamagabe (both *p*<0.001; Figure 2; Tables S8-11). In Musanze, both the inoculant-alone treatments increased small-grade yield relative to untreated controls in 2025A (both *p*<0.001), with only the 250 g ha⁻¹ treatment remaining greater than the control in 2025B (*p*=0.012). In Nyamagabe, the inoculant alone had little effect during 2025A, but tended to increase small-grade yield in comparison to the control in 2025B. Combining the inoculant with standard fertilization generally outperformed the inoculant applied alone, although responses to inoculant and fertilizer co-application varied considerably. For example, low-dose inoculant increased small-grade yield compared to standard fertilization in Nyamagabe during 2025B (+3.72 t ha⁻¹, *p*<0.001), but reduced it in Musanze during 2025A (-2.37 t ha⁻¹, *p*=0.05). Comparisons between inoculant application rates showed similarly inconsistent responses, with the higher application rate generally producing greater small-grade yield than the lower rate in Musanze (both *p*<0.05), but not in Nyamagabe (Figure 2a-b; Table S4, S8).

**Figure 2.**
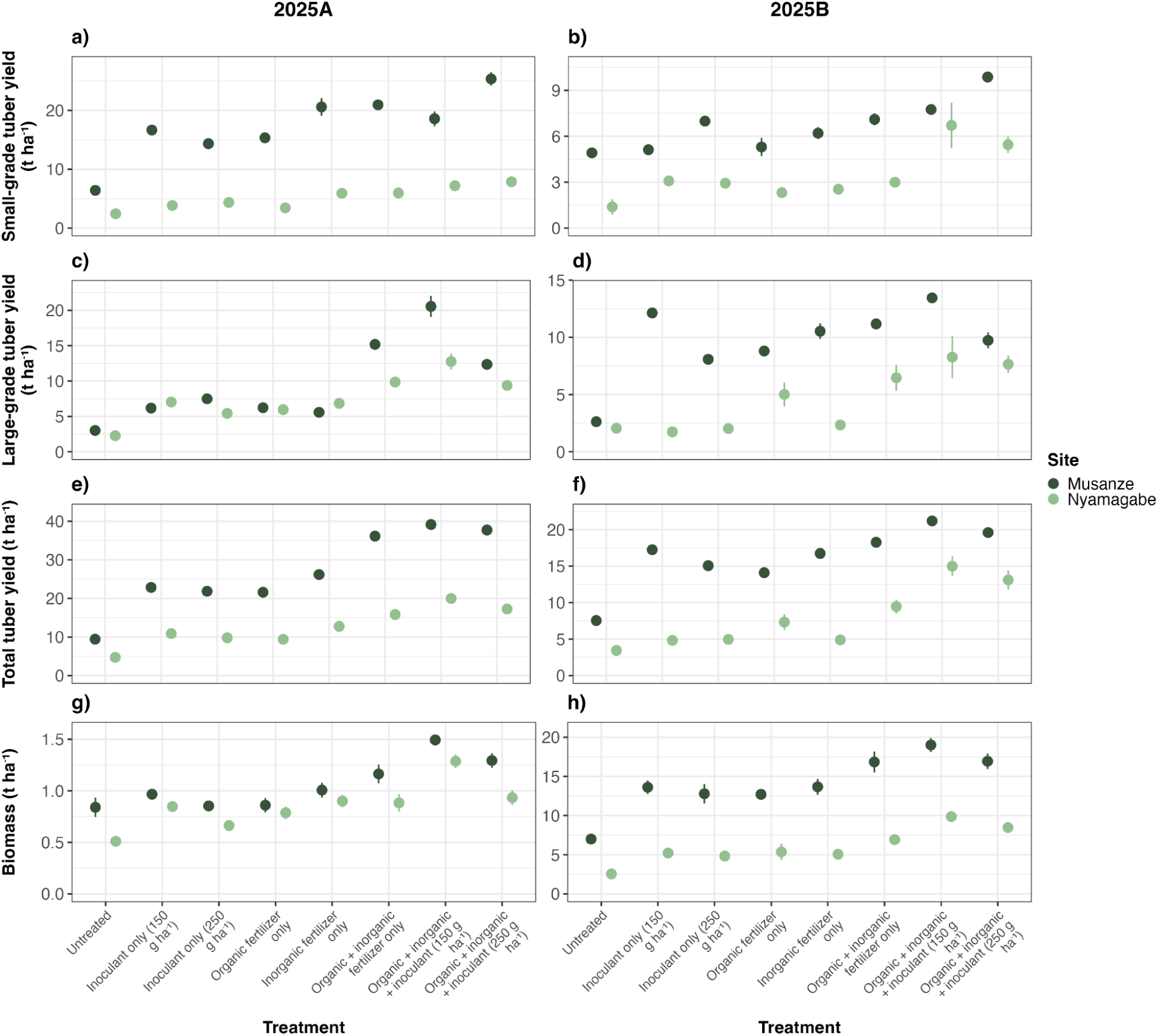
Differences in a-b) small-grade potato tuber yield; c-d) large-grade tuber yield; e-f) total tuber yield; and g-h) potato plant biomass across treatments in seasons 2025A (left column) and 2025B (right column). In each panel, points represent treatment x site means, error bars indicate standard error (n=3), and shades of green indicate site. Full statistical outputs are located in Table S4-11.

Patterns in large-grade potato tubers, which are primarily used as commodities for consumption, diverged from those of small-grade tubers in that they were relatively more consistent across sites and seasons (Figure 2c-d; Table S5). For large-grade tubers, inoculant-alone treatments generally increased yield relative to untreated controls, particularly in Musanze where both application rates increased yield in both seasons (all *p*<0.01). Additionally, standard fertilization typically produced greater large-grade yield than the inoculant alone (all *p*<0.001), except in Musanze during 2025B. Combining the inoculant with standard fertilization often increased large-grade yield relative to standard fertilization, although responses depended on application rate. In 2025A, low-dose of the inoculant combined with standard fertilization increased large-grade yield in both Musanze (+5.37 t ha⁻¹, *p*<0.001) and Nyamagabe (+2.91 t ha⁻¹, *p*=0.007), while the corresponding high-dose treatment reduced yield in Musanze (-2.82 t ha⁻¹, *p*=0.008) and had no effect in Nyamagabe. During 2025B, positive effects of low-dose inoculant persisted only in Musanze (+2.28 t ha⁻¹, *p*=0.049). Additionally, the lower application rate frequently outperformed the higher rate for large-grade yield, including in Musanze during both seasons (+8.19 t ha⁻¹ in 2025A, +3.71 t ha⁻¹ in 2025B; both *p*<0.01) and in Nyamagabe during 2025A (+3.39 t ha⁻¹, *p*=0.002; Figure 2c-d; Table S5, S9).

Total yield responses generally mirror those of large-grade tubers. When applied alone, the inoculant generally increased total yield relative to untreated controls (i.e., no fertilizer, no inoculant). In Musanze, both application rates (150 and 250 g ha⁻¹) increased total yield in both seasons, with gains of 13.41 and 12.41 t ha⁻¹ in 2025A, and 9.7 and 7.52 t ha⁻¹ in 2025B, respectively (all *p*<0.001). Similar increases occurred in Nyamagabe during 2025A only (+6.17 and +5.08 t ha⁻¹ for low and high dose rates, respectively; both *p*<0.001). Despite these increases, standard fertilization practices generally outperformed the inoculant applied alone, with inoculant-only treatments yielding less than standard practice in nearly all site x season combinations (all *p*<0.001 except Musanze 2025B low-dose, *p*=0.31). In contrast, combining the inoculant with standard fertilization frequently increased total yield relative to the standard practice, particularly at the 150 g ha⁻¹ rate. With the low dose, yield increases under combined application occurred in both sites and seasons (Nyamagabe +4.15 t ha⁻¹ in 2025A, +5.53 t ha⁻¹ in 2025B; Musanze: +3.00 t ha⁻¹ in 2025A, +2.92 t ha⁻¹ in 2025B; all *p*<0.05). At the 250 g ha⁻¹ rate, yield increases relative to standard practice occurred only in Nyamagabe during 2025B (+3.64 t ha⁻¹, *p*<0.001; Figure 2e-f; Table S6, S10).

These treatment effects on small-grade, large-grade, and total yield were reflected in percent-change analyses relative to standard fertilization practices, where only combined inoculant + fertilizer treatments generally produced positive yield responses relative to the agronomic standard (Figure 3). Additionally, when compared within individual seasons, biomass responses largely mirrored those of total and large-grade yield, with standard fertilization plus low-dose inoculant increasing biomass relative to standard practice alone in both 2025A (+0.37 t ha⁻¹, *p*<0.001) and 2025B (+2.56 t ha⁻¹, *p*=0.005), while the high-dose treatment had no effects (Figure 2g-h; Table S7). The effects of treatment on biomass also differed significantly with season across both sites (both p<0.001; Table S11), consistent with the season-specific magnitude of the low-dose inoculant effect noted above.

**Figure 3.**
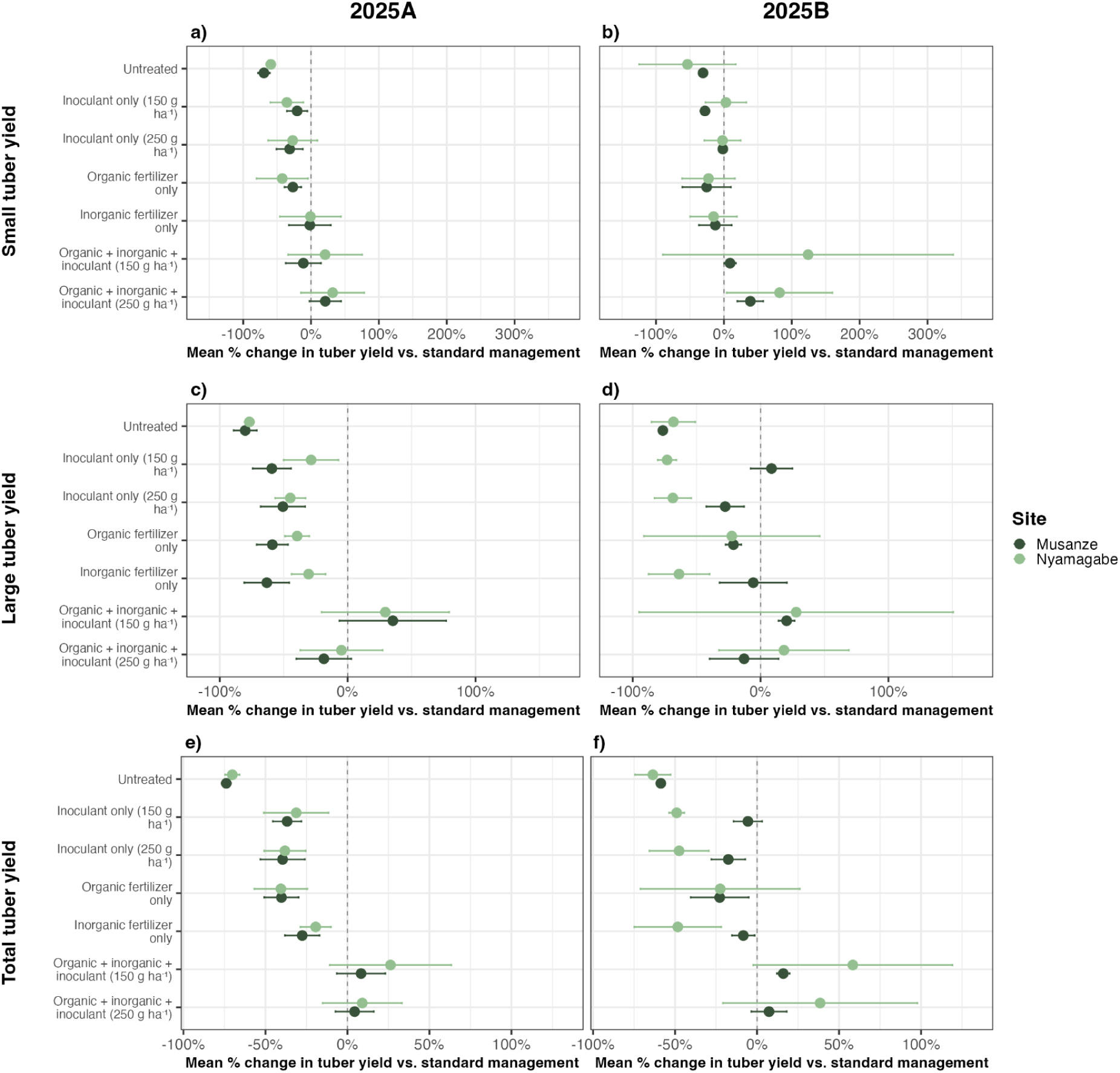
Mean percent change in a-b) small potato tuber yield; c-d) large potato tuber yield; and e-f) total yield relative to standard fertilization practices (i.e., co-application of organic and inorganic fertilizer) across seasons 2025A and 2025B. In all panels, points indicate mean percent change, error bars indicate 95% confidence intervals, and shades of green indicate site.

### Maize grain yield and plant biomass

Maize grain yield and plant biomass were significantly affected by treatment and site across both seasons (Figure 4; Tables S12–S13). Seasonal variability also moderated grain yield response to our treatments across both locations (Kayonza *p*<0.001; Ngoma *p*=0.021; Figure 4; Tables S14-S15). In 2025A, treatment responses differed between sites (*p*=0.004), but overall patterns were consistent across Kayonza and Ngoma. Inoculant applied alone increased grain yield relative to the untreated control at both application rates in both sites (all *p*<0.001), whereas standard fertilization consistently outperformed our inoculant when applied alone (all *p*<0.001). However, combining inoculation with standard fertilization increased grain yield relative to standard practice alone across both sites, with larger effects generally observed at the higher inoculation rate (all *p*<0.01). Direct comparisons between inoculation rates combined with standard fertilization confirmed greater yield gains at 250 g ha⁻¹ than at 150 g ha⁻¹ in both sites (*p*<0.001). In 2025B, treatment effects were consistent across sites (*p*=0.13). Applications of only our inoculant increased grain yield only at 250 g ha⁻¹ (+1.92 t ha⁻¹, *p*<0.001), while yields under standard fertilization again exceeded the inoculant when it was applied alone (all *p*≤0.014). As in 2025A, combining the inoculant with standard fertilization increased grain yield at both application rates, with stronger responses at 250 g ha⁻¹ (*p*<0.001), and the higher rate again outperforming the lower rate (p=0.014; Figure 4a-b; Table S12). These responses were generally mirrored in analyses of percent change in maize yield relative to standard fertilization practices, with only treatments that involved application of both the inoculant and standard fertilization leading to positive percent changes in yield compared to the standard (Figure 5; Tables S12, S14).

**Figure 4.**
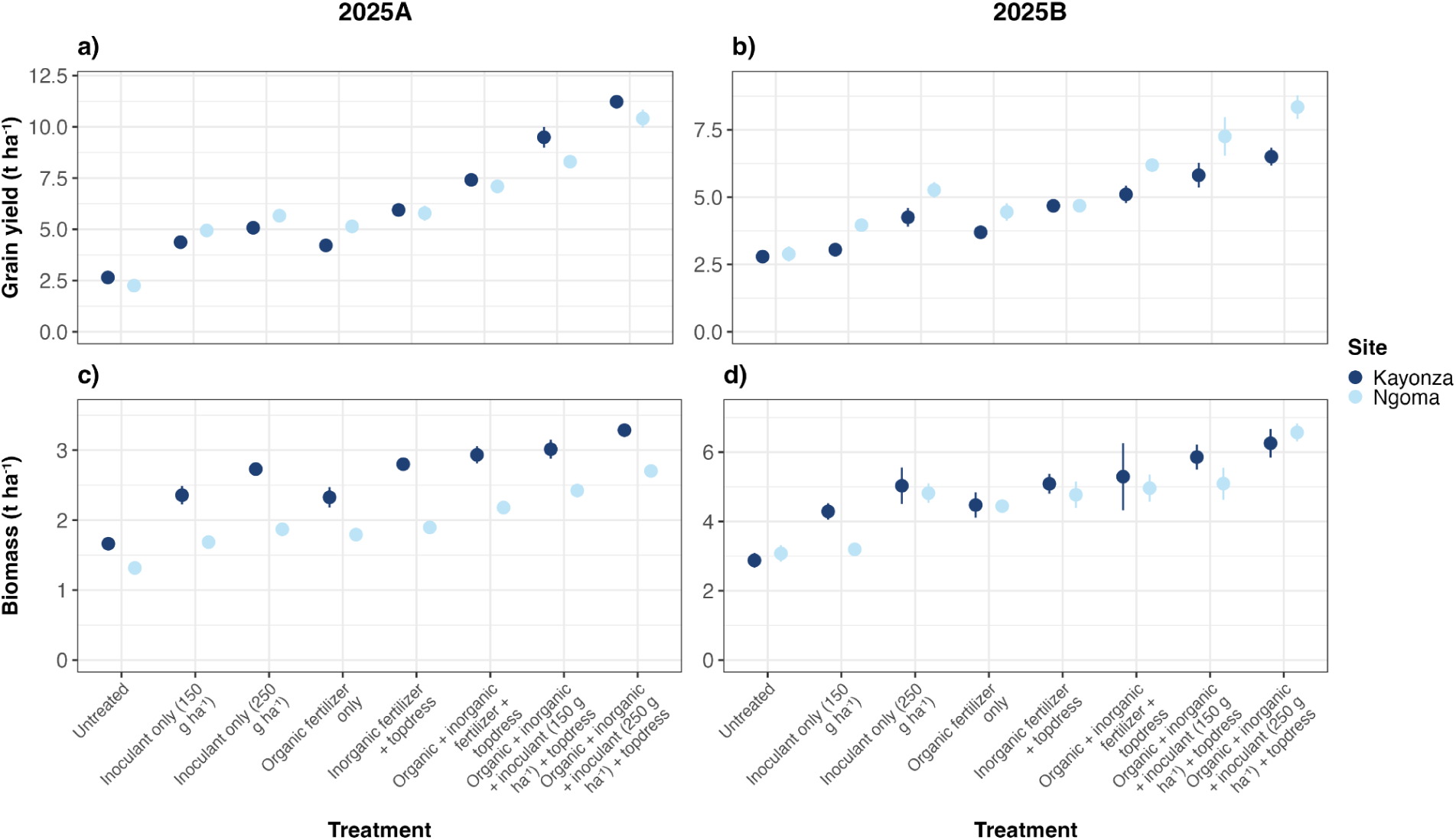
Differences in a-b) maize grain yield and c-d) maize plant biomass across treatments as well as seasons 2025A (left column) and 2025B (right column). In each panel, points represent treatment x site means, error bars indicate standard error (n=3), and shades of blue indicate site. Full statistical outputs are located in Tables S12-15.

**Figure 5.**
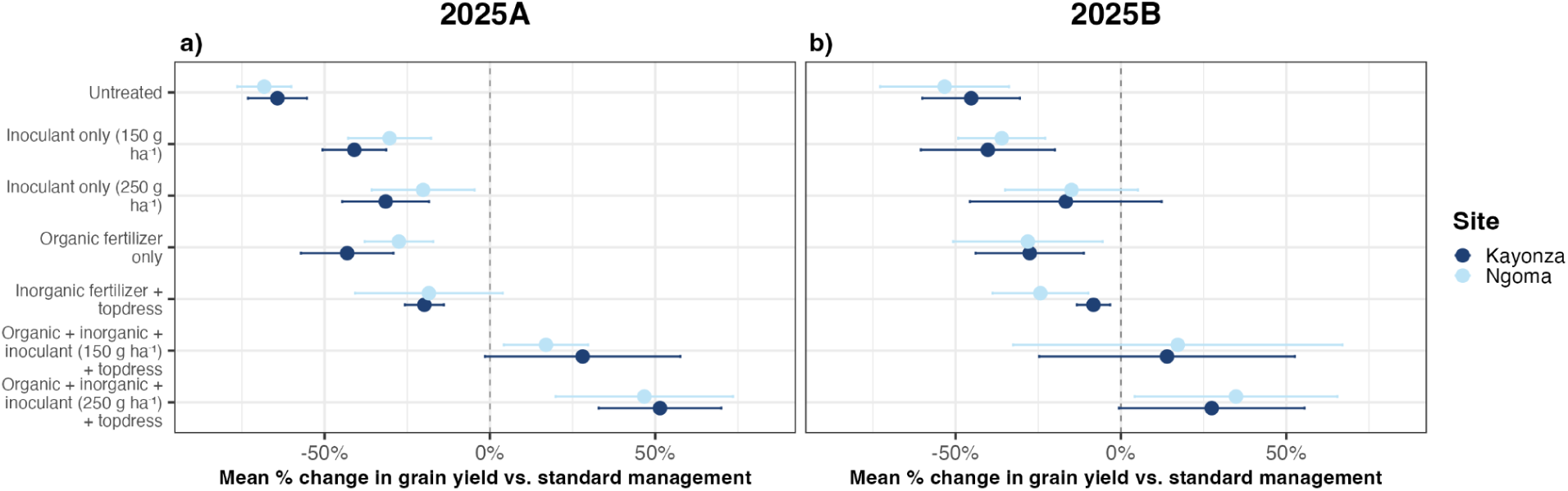
Mean percent change in maize grain yield relative to standard fertilization practices (i.e., co-application of organic and inorganic fertilizer) across seasons a) 2025A and b) 2025B. In both panels, points indicate mean percent change, error bars indicate 95% confidence intervals, and colors indicate field site.

Maize plant biomass responses generally mirrored those of grain yield (Figure 4c-d; Tables S13, S15). Treatment responses did not differ based on site in either season (2025A *p*=0.13; 2025B *p*=0.68), with inoculant-alone treatments increasing biomass relative to untreated controls (all p<0.001), but standard fertilization producing greater biomass than inoculant-alone treatments (all p<0.05). Unlike grain yield, biomass responses to treatment depended on season only in Ngoma (p<0.001; Figure 4c-d; Table S15).Combining inoculation with standard fertilization increased biomass relative to the standard at the 250 g ha⁻¹ rate (2025A: +0.44 t ha⁻¹, *p*<0.001; 2025B: (+1.29 t ha⁻¹, *p*=0.007), but not at 150 g ha⁻¹.

Additionally, direct comparisons between application rates further indicated greater biomass production under the 250 g ha⁻¹ than the 150 g ha⁻¹ treatment in both 2025A (+0.28 t ha⁻¹, *p*=0.009) and 2025B (+0.94 t ha⁻¹, *p*=0.042; Figure 4c-d; Tables S13, S15).

### Maize and potato nutrient density

Despite increases in yield associated with several treatments involving inoculant application, we observed little evidence of changes in nutrient density in harvested maize or potato tissues (Figures S4-5; Tables S16-17). While treatment significantly affected crude protein concentration in maize, (*p*=0.009), with protein concentrations being higher under standard fertilization than under the organic fertilizer-only treatment (*p*=0.032), nutrient densities were largely only affected by site. Specifically, maize phosphorus (*p*=0.009), calcium (*p*=0.001), and organic carbon concentrations (*p*=0.001) were all greater in Ngoma than in Kayonza (Figure S4; Table S17). Likewise, potato iron concentrations differed only by district (*p*<0.001), with substantially greater concentrations observed in Musanze than Nyamagabe (Figure S4; Table S16).

## Discussion

In this study, we evaluated whether a multi-species microbial inoculant comprised of spore-forming and Gram-positive *Bacillus* and *Streptomyces* bacterial species, could increase crop productivity across a range of agroecological zones in Rwanda. Across maize and potato systems, co-application of the inoculant with standard fertility inputs (including organic and inorganic fertilizers) increased yield and plant biomass beyond that achieved with fertilization alone. When applied in isolation, the inoculant-only treatments increased productivity relative to untreated controls, but generally did not match the productivity achieved under standard fertilization. Positive responses to the inoculant were observed across all crops, field sites, and growing seasons, but their magnitude varied substantially. For instance, potato systems generally exhibited stronger and more consistent responses, particularly with respect to large-grade tuber production, whereas maize responses varied more strongly across sites and seasons (Figures 2-5). Additionally, while we observed some potential dose-dependent trade-offs between small- versus large-grade potato tuber production, the overall increases in the yield of both size classes may be especially relevant in the smallholder farming context; given that large-grade tubers are typically used as commodities and small-grade tubers as seeds for future growing seasons or in local, low-value markets. Altogether, these results imply that microbial inoculants provide dual economic benefits to producers suggesting that microbial inoculants can serve as a particularly effective complement to existing fertility management practices (Vanlauwe et al, 2001), whereby co-application of inoculants and fertilizers has the ability to increase crop productivity beyond what can be achieved with fertilizers alone.

Despite variability in response magnitude within crops and across sites, the consistency of positive effects, particularly with respect to co-application of the microbial inoculant with standard fertilizers, supports a growing body of evidence demonstrating that multi-species inoculants are effective at improving crop performance under field conditions, especially in tropical agricultural systems (e.g., Okereke et al., 2000; Adesemoye et al., 2009; Bashan et al., 2014; Jiang et al., 2023; Liu et al., 2023; Francioli et al., 2025; Bukombe et al., 2026) as well as in comparison to single-strain inoculants (Liu et al., 2023). A recent global meta-analysis found that consortium inoculants enhanced plant productivity by ∼48% relative to untreated controls compared to ∼29% for single-species inoculants (Liu et al., 2023), an advantage attributed to functional complementarity among strains (Kaminsky et al., 2019) and greater resilience across variable environmental conditions (Liu et al., 2023). The yield gains observed in our study, ranging from 6-51% above fertilization alone for maize and 3-58% for potato, fall at or above the upper range of responses commonly reported for single-strain inoculants in tropical agroecosystems (e.g., Okereke et al., 2000; Thilakarathna and Raizada, 2017; Chibeba et al., 2018; Getachew Gebrehana and Abeble Dagnaw, 2020). Notably, the responses observed in our study were achieved under open field conditions, where inoculant efficacy is often more variable and lower than that of greenhouse environments (Liu et al., 2023). Given that most previous field studies from sub-Saharan Africa focus on rhizobial inoculation in legume crops, our findings provide additional support for the widespread utility of microbial consortia within the region (Bukombe et al., 2026). Importantly, our observed yield increases were not accompanied by reduced nutrient density in either crop. Instead, nutrient density was driven primarily by site-level differences, indicating that increased productivity as a result of inoculation did not come at a measurable cost of potato or maize nutritional quality, which may have been expected given site- and study-specific evidence of crop nutrient dilution with shifts in fertility management (Buerkert et al., 1998; Feil et al., 2005; Cakmak, 2008; Davis, 2009; Zhao et al., 2020). That being said, we note this null result should be interpreted cautiously: with three replicates per site and season, our power to detect modest reductions in nutrient concentration was limited, and the nutrient panel was narrow, particularly for potato, where only iron was quantified. Confirming the absence of a yield-quality trade-off will require broader nutrient profiling and greater replication. To the extent that yield gains are not offset by nutrient dilution, greater nutritional content can be produced per unit land area – an especially relevant benefit given evidence that rising atmospheric CO₂ concentrations are already contributing to global declines in crop nutrient density (Myers et al., 2014), with micronutrient deficiencies remaining a persistent public health concern across East Africa (Joy et al., 2014; Galani et al., 2022).

Our results also indicate that the effects of microbial enrichment are likely to be dose-dependent. Across both sites, co-application of standard fertilizers with the lower inoculant dose (i.e., 150 g ha⁻¹) consistently increased large-grade tuber yield across all site x season combinations, whereas the higher dose (i.e., 250 g ha⁻¹) was less consistent and in some cases, notably in Musanze during season 2025A, reduced large-grade yield relative to fertilization alone. Small-grade tuber yield showed more variable responses but were frequently positive across inoculant treatments, with high doses tending to increase small-grade yield relative to the low dose, suggesting a shift in plant resource toward increased tuber initiation at the expense of bulking. This pattern is consistent with well-established fertilizer effects on potato yield structure, where intermediate nutrient availability tends to maximize marketable yield, while higher nutrient stimulation can increase tuber set, but reduce average tuber size (Arsenault et al., 2001; Cambouris et al., 2007; Rosen and Bierman, 2008; Tsai et al., 2025). This aligns with broader observations from East Africa highland systems where management intensity can regulate marketable versus seed grade distribution (Gildemacher et al., 2009; Kaguongo et al., 2014; Schulte-Geldermann et al., 2022). In contrast, maize exhibited a clear positive dose-response relationship, with the high inoculant dose consistently outperforming the low for both grain yield and biomass. This is consistent with general crop yield-resource response theory in which productivity increases with resource availability up to an agronomic ceiling (Lobell et al., 2009; Dhakal and Lange, 2021), and with empirical evidence of strong fertilizer responsiveness in sub-Saharan African maize systems under smallholder management (Mukuralinda et al., 2010; Vanlauwe et al., 2015). While further research is needed to refine economically optimal dose ranges under different fertilizer regimes, these dose- and crop-specific resource allocation patterns indicate that optimal microbial inoculant application rates should be defined by both crop type and production objective (e.g., marketable or large-grade tubers versus seed or small-grade output in potato), rather than applied uniformly across agroecosystems.

Despite the directionally consistent effects of microbial inoculation, the magnitudes of treatment effects varied considerably. In particular, treatment effects were stronger and more consistent in both potato sites (i.e., Nyamagabe and Musanze) than in maize (Kayonza and Ngoma). Within crop types, treatment effects were also typically more variable within the lower-fertility site compared to the higher-fertility site, with seasonal effects of, for instance, contrasting weather patterns potentially contributing to shifts in inoculant efficacy and the observed trade-offs between yield and plant biomass across seasons (Figures 2-5; Figure S1). While this may be in part reflective of different interactions between the inoculant and crops, the contrasts in soil conditions across the four sites, including differences in pH, cation exchange capacity, texture, organic carbon, and nutrient availability, are consistent with local edaphic conditions playing a role in shaping inoculant performance (Azarbad and Junker, 2024; Figures S2-S3). One possibility is that highly acidic conditions at Nyamagabe, which exhibited the greatest variability in treatment responses, imposed constraints on Bacilli activity and spore formation under low pH (Iqbal, 2026); however, because pH co-varied with other soil properties across sites, this remains a candidate explanation rather than an isolated effect. Broadly, these patterns of inoculant efficacy are consistent with previous work showing that inoculant performance is context-dependent, and shaped by interactions among soil properties, resident microbiomes, plant traits, and climate (Jiang et al., 2023; Azarbad and Junker, 2024). However, because soil variables co-varied across our study sites, our experimental design cannot isolate the effects of individual soil properties; responses instead point to integrated differences in overall soil health as the more likely driver.

Although inoculants applied alone outperformed untreated controls, they were not able to achieve the same productivity levels observed in standard fertilizer treatments (Figures 2-5). The relatively weaker efficacy of inoculant-only treatments compared with fertilizer may reflect the timescale over which microbially mediated soil processes develop. Microbial-mediated build-up of the organic matter pools and shifts in aggregation that lead to improvements in nutrient cycling, soil structure, and water retention may emerge gradually over multiple growing seasons (e.g., Plaza-Bonilla et al., 2013; Jat et al., 2019; Ferreira et al., 2020), and are unlikely to deliver the soil fertility benefits required to compete with fertilizer within the six-month duration of this study. As such, longer-term, multi-season experiments are needed to determine whether repeated inoculant applications progressively improve soil function, particularly in ways that narrow productivity gaps between inoculant- and fertilizer-alone treatments. Nevertheless, microbial enrichment appeared to achieve approximately half of the effects of fertilizer treatment. While further research involving co-application of inoculants with reduced fertilization rates is needed to confirm, these findings imply a potential opportunity for inoculants to reduce fertilizer application to avoid soil depletion. In particular, inoculants based on locally adapted or native microbial strains could also improve performance, as natively-sourced strains may be better suited to local conditions where they co-evolved alongside resident microbial communities and within the context of local soil physicochemical conditions (Bashan et al., 2014; Jiang et al., 2023).

Though our results demonstrate strong potential of multi-species inoculants to augment fertility management, our study design does not allow us to identify the mechanisms responsible for these responses. However, the taxonomic composition of the inoculant is indicative of several plausible mechanisms. For instance, Bacilli and *Streptomyces* species are well-known to enhance multiple pathways of plant growth promotion (Sousa and Olivares, 2016; Olanrewaju and Babalola, 2019), including phosphorus solubilization into plant-available forms (Alori et al., 2017; Chouyia et al., 2022; Ahash et al., 2025; Sun et al., 2026), phytohormone production and root development (Al-Tammar and Khalifa, 2023), pathogen suppression (Iqbal, 2026), and improved plant tolerance to environmental stressors, especially drought (Chattaraj et al., 2025; Kesavardhini et al., 2025; Mishra and Choudhary, 2026). Introduced microbial inoculants also interact with existing soil microbiomes and plant communities in complex, context-dependent ways that alter nutrient cycling and plant performance, ultimately influencing agronomic outcomes (Li et al., 2024; Francioli et al., 2025; Huang et al., 2025). The relative importance of these mechanisms will require further environmental studies across sites and seasons.

Specifically, future work should prioritize direct characterization of several aspects of inoculant activity under field conditions, including persistence of strains included within the inoculant, shifts in resident microbiome composition, and temporal patterns of microbial gene expression throughout the growing season.

Overall, our findings show that multi-species microbial inoculants can enhance crop productivity, especially when integrated with existing soil fertility management practices, across contrasting tropical field conditions. Despite variation across crops, sites, and seasons, the consistently positive effects of co-application across all study locations suggest that microbial consortia are a promising tool for maximizing yield in smallholder agricultural systems. Additionally, while inoculant-only treatments did not achieve the same productivity as standard fertilization practices, the benefits of inoculant and fertilizer co-application imply that comparable yields may potentially be maintained with reduced fertilizer inputs (e.g., Adesemoye et al., 2009). Although further work is needed to directly test fertilizer reduction scenarios across crops, soil conditions, and climates, this may represent an important economic opportunity for smallholder farmers that also delivers co-benefits associated with fertilizer reduction, particularly given recent increases in fertilizer costs and continued vulnerability of supply chains. Together, these results support the use of microbial inoculants as part of integrated soil fertility management in tropical agroecosystems, with the potential to deliver continued economic and environmental benefits in the face of a rapidly changing climate as well as increasing needs for sustained food security.

## Acknowledgements

The authors would like to thank the Rwanda Agriculture and Animal Resources Development Board (RAB) for conducting the field trials, as well as providing the data and scientific support to study authors. Special thanks go to our laboratory staff for producing the microbial inoculant and their analysis of soil and plant metrics, as well as all field employees who facilitated the field experiments across Rwanda. We would also like to thank Dr. Martin Voss, Chief Innovation Officer at Oath Biome, for his work in the registration of the microbial inoculants used in this study for application to Rwandan soils and advice on project execution.

## References

Adesemoye, A.O., Torbert, H.A., Kloepper, J.W., 2009. Plant growth-promoting rhizobacteria allow reduced application rates of chemical fertilizers. Microbial Ecology 58, 921–929. 10.1007/s00248-009-9531-y.

Ahash, S., Manikandan, K., Sivasankari Devi, T., Elamathi, S., Maragatham, S., Subrahmaniyan, K., 2025. Phosphate-solubilizing microorganisms for sustainable phosphorus management in rice. Rhizosphere 34, 101096. 10.1016/j.rhisph.2025.101096.

Alori, E.T., Glick, B.R., Babalola, O.O., 2017. Microbial phosphorus solubilization and its potential for use in sustainable agriculture. Frontiers in Microbiology 8, 971. 10.3389/fmicb.2017.00971.

Al-Tammar, F.K., Khalifa, A.Y.Z., 2023. An update about plant growth promoting *Streptomyces* species. Journal of Applied Biology & Biotechnology. 10.7324/JABB.2023.130126.

Amankwah, A., Ambel, A., Gourlay, S., Kilic, T., Markhof, Y., Wollburg, P., 2025. Smallholder farming, fertilizer use, and the polycrisis period: Cross-country evidence from longitudinal surveys in Sub-Saharan Africa. Food Policy 133, 102885. 10.1016/j.foodpol.2025.102885.

Arsenault, W.J., LeBlanc, D.A., Tai, G.C.C., Boswall, P., 2001. Effects of nitrogen application and seedpiece spacing on yield and tuber size distribution in eight potato cultivars. American Journal of Potato Research 78, 301–309. 10.1007/BF02875695.

Assefa, T.W., Berhane, G., Abate, G.T., Abay, K.A., 2025. Fertilizer demand and profitability amid global fuel-food-fertilizer crisis: Evidence from Ethiopia. Food Policy 133, 102785. 10.1016/j.foodpol.2024.102785.

Averill, C., Anthony, M.A., Baldrian, P., Finkbeiner, F., van den Hoogen, J., Kiers, T., Kohout, P., Hirt, E., Smith, G.R., Crowther, T.W., 2022. Defending Earth’s terrestrial microbiome. Nature Microbiology 7, 1717–1725. 10.1038/s41564-022-01228-3.

Azarbad, H., Junker, R.R., 2024. Biological and experimental factors that define the effectiveness of microbial inoculation on plant traits: A meta-analysis. ISME Communications 4, ycae122. 10.1093/ismeco/ycae122.

Bankevich, A., Nurk, S., Antipov, D., Gurevich, A.A., Dvorkin, M., Kulikov, A.S., Lesin, V.M., Nikolenko, S.I., Pham, S., Prjibelski, A.D., Pyshkin, A.V., Sirotkin, A.V., Vyahhi, N., Tesler, G., Alekseyev, M.A., Pevzner, P.A., 2012. SPAdes: A new genome assembly algorithm and its applications to single-cell sequencing. Journal of Computational Biology 19, 455–477. 10.1089/cmb.2012.0021.

Bashan, Y., de-Bashan, L.E., Prabhu, S.R., Hernandez, J.-P., 2014. Advances in plant growth-promoting bacterial inoculant technology: formulations and practical perspectives (1998–2013). Plant and Soil 378, 1–33. 10.1007/s11104-013-1956-x.

Black, C.A. (Ed.), 1965. Methods of Soil Analysis: Part 1—Physical and Mineralogical Properties, Including Statistics of Measurement and Sampling, 1st ed. Agronomy Monographs. Wiley, Madison, WI. 10.2134/agronmonogr9.1.

Bouyoucos, G.J., 1962. Hydrometer method improved for making particle size analyses of soils. Agronomy Journal 54, 464–465. 10.2134/agronj1962.00021962005400050028x.

Bray, R.H., Kurtz, L.T., 1945. Determination of total, organic, and available forms of phosphorus in soils. Soil Science 59, 39–46. 10.1097/00010694-194501000-00006.

Bremner, J.M., Mulvaney, C.S., 1982. Nitrogen—Total, in: Page, A.L. (Ed.), Methods of Soil Analysis, Part 2: Chemical and Microbiological Properties, 2nd ed. Agronomy Monographs. Wiley, Madison, WI, pp. 595–624. 10.2134/agronmonogr9.2.2ed.c31.

Buerkert, A., Haake, C., Ruckwied, M., Marschner, H., 1998. Phosphorus application affects the nutritional quality of millet grain in the Sahel. Field Crops Research 57, 223–235. 10.1016/S0378-4290(97)00136-6.

Bukombe, B., Edlund, A., Saavedra, P.G., Mberwa, J.W., Gilbert, J.A., Naramabuye, F.X., Sirikare, S., Hansen, P.M., Jansson, J.K., Crowther, T.W., 2026. Microbial inoculation increases maize yield and root biomass across smallholder farming systems in Rwanda. EarthArXiv [Preprint]. 10.31223/X53R2P.

Cakmak, I., 2008. Enrichment of cereal grains with zinc: Agronomic or genetic biofortification? Plant and Soil 302, 1–17. 10.1007/s11104-007-9466-3.

Cambouris, A.N., Zebarth, B.J., Nolin, M.C., Laverdière, M.R., 2007. Response to added nitrogen of a continuous potato sequence as related to sand thickness over clay. Canadian Journal of Plant Science 87, 829–839. 10.4141/P06-126.

Chattaraj, S., Samantaray, A., Ganguly, A., Thatoi, H., 2025. Employing plant growth-promoting rhizobacteria for abiotic stress mitigation in plants: With a focus on drought stress. Discover Applied Sciences 7, 68. 10.1007/s42452-025-06468-6.

Chaumeil, P.-A., Mussig, A.J., Hugenholtz, P., Parks, D.H., 2022. GTDB-Tk v2: Memory friendly classification with the genome taxonomy database. Bioinformatics 38, 5315–5316. 10.1093/bioinformatics/btac672.

Chen, S., 2023. Ultrafast one-pass FASTQ data preprocessing, quality control, and deduplication using fastp. iMeta 2, e107. 10.1002/imt2.107.

Chen, S., Zhou, Y., Chen, Y., Gu, J., 2018. fastp: An ultra-fast all-in-one FASTQ preprocessor. Bioinformatics 34, i884–i890. 10.1093/bioinformatics/bty560.

Chibeba, A.M., Kyei-Boahen, S., Guimarães, M. de F., Nogueira, M.A., Hungria, M., 2018. Feasibility of transference of inoculation-related technologies: A case study of evaluation of soybean rhizobial strains under the agro-climatic conditions of Brazil and Mozambique. Agriculture, Ecosystems & Environment 261, 230–240. 10.1016/j.agee.2017.06.037.

Chklovski, A., Parks, D.H., Woodcroft, B.J., Tyson, G.W., 2023. CheckM2: A rapid, scalable and accurate tool for assessing microbial genome quality using machine learning. Nature Methods 20, 1203–1212. 10.1038/s41592-023-01940-w.

Chouyia, F.E., Ventorino, V., Pepe, O., 2022. Diversity, mechanisms and beneficial features of phosphate-solubilizing *Streptomyces* in sustainable agriculture: A review. Frontiers in Plant Science 13, 1035358. 10.3389/fpls.2022.1035358.

Davis, D.R., 2009. Declining fruit and vegetable nutrient composition: What is the evidence? HortScience 44, 15–19. 10.21273/HORTSCI.44.1.15.

Dhakal, C., Lange, K., 2021. Crop yield response functions in nutrient application: A review. Agronomy Journal 113, 5222–5234. 10.1002/agj2.20863.

Edlund, A., Espinoza, J.L., Basu, S.S., Grama, A., McCorrison, J., Boreux, V., Gilbert, J.A., Crowther, T.W., 2026. Complex microbial consortia improve yield and physiological performance of leafy greens under deficit irrigation. bioRxiv [Preprint]. 10.64898/2026.04.05.716566.

Espinoza, J.L., Dupont, C.L., 2022. VEBA: A modular end-to-end suite for in silico recovery, clustering, and analysis of prokaryotic, microeukaryotic, and viral genomes from metagenomes. BMC Bioinformatics 23, 419. 10.1186/s12859-022-04973-8.

Espinoza, J.L., Phillips, A., Prentice, M.B., Tan, G.S., Kamath, P.L., Lloyd, K.G., Dupont, C.L., 2024. Unveiling the microbial realm with VEBA 2.0: A modular bioinformatics suite for end-to-end genome-resolved prokaryotic, (micro)eukaryotic and viral multi-omics from either short- or long-read sequencing. Nucleic Acids Research 52, e63. 10.1093/nar/gkae528.

Feil, B., Moser, S.B., Jampatong, S., Stamp, P., 2005. Mineral composition of the grains of tropical maize varieties as affected by pre-anthesis drought and rate of nitrogen fertilization. Crop Science 45, 516–523. 10.2135/cropsci2005.0516.

Ferreira, C. dos R., Silva Neto, E.C. da, Pereira, M.G., Guedes, J. do N., Rosset, J.S., Anjos, L.H.C. dos, 2020. Dynamics of soil aggregation and organic carbon fractions over 23 years of no-till management. Soil and Tillage Research 198, 104533. 10.1016/j.still.2019.104533.

Fox, J., Weisberg, S., Price, B., Adler, D., Bates, D., Baud-Bovy, G., Bolker, B., Ellison, S., Firth, D., Friendly, M., Gorjanc, G., Graves, S., Heiberger, R., Krivitsky, P., Laboissiere, R., Maechler, G., Monette, G., Murdoch, D., Nilsson, H., Ogle, D., Proctor, I., Ripley, B., Short, T., Venables, W., Walker, S., Winsemius, D., Zeileis, A., 2026. car: Companion to Applied Regression. R package.

Francioli, D., Kampouris, I.D., Kuhl-Nagel, T., Babin, D., Sommermann, L., Behr, J.H., Chowdhury, S.P., Zrenner, R., Moradtalab, N., Schloter, M., Geistlinger, J., Ludewig, U., Neumann, G., Smalla, K., Grosch, R., 2025. Microbial inoculants modulate the rhizosphere microbiome, alleviate plant stress responses, and enhance maize growth at field scale. Genome Biology 26, 148. 10.1186/s13059-025-03621-7.

Galani, Y.J.H., Orfila, C., Gong, Y.Y., 2022. A review of micronutrient deficiencies and analysis of maize contribution to nutrient requirements of women and children in Eastern and Southern Africa. Critical Reviews in Food Science and Nutrition 62, 1568–1591. 10.1080/10408398.2020.1844636.

Getachew Gebrehana, Z., Abeble Dagnaw, L., 2020. Response of soybean to rhizobial inoculation and starter N fertilizer on Nitisols of Assosa and Begi areas, Western Ethiopia. Environmental Systems Research 9, 14. 10.1186/s40068-020-00174-5.

Gildemacher, P.R., Kaguongo, W., Ortiz, O., Tesfaye, A., Woldegiorgis, G., Wagoire, W.W., Kakuhenzire, R., Kinyae, P.M., Nyongesa, M., Struik, P.C., Leeuwis, C., 2009. Improving potato production in Kenya, Uganda and Ethiopia: A system diagnosis. Potato Research 52, 173–205. 10.1007/s11540-009-9127-4.

Guo, S., Chen, Y., Chen, X., Chen, Y., Yang, L., Wang, L., Qin, Y., Li, M., Chen, F., Mi, G., Gu, R., Yuan, L. 2020. Grain mineral accumulation changes in Chinese maize cultivars released in different decades and the responses to nitrogen fertilizer. Frontiers in Plant Science. 10:1662. 10.3389/fpls.2019.01662.

Huang, Y., Tang, S., Liu, R., Yu, P., Liu, J., Xiao, T., Zhang, Y., Fan, M., Zhang, F., Ni, B., 2025. Soil organic carbon mediates plant immunity-rhizosphere microbiome interactions and controls colonization resistance to microbial inoculants. Cell Host & Microbe 33, 1929–1944.e7. 10.1016/j.chom.2025.10.002.

Iqbal, M., 2026. Microbial biocontrol agents and the rhizosphere microbiome: Integrating ecological function and climate resilience in sustainable agriculture. Frontiers in Microbiology 17, 1771649. 10.3389/fmicb.2026.1771649.

Jat, H.S., Datta, A., Choudhary, M., Yadav, A.K., Choudhary, V., Sharma, P.C., Gathala, M.K., Jat, M.L., McDonald, A., 2019. Effects of tillage, crop establishment and diversification on soil organic carbon, aggregation, aggregate associated carbon and productivity in cereal systems of semi-arid Northwest India. Soil and Tillage Research 190, 128–138. 10.1016/j.still.2019.03.005.

Jiang, M., Delgado-Baquerizo, M., Yuan, M.M., Ding, J., Yergeau, E., Zhou, J., Crowther, T.W., Liang, Y., 2023. Home-based microbial solution to boost crop growth in low-fertility soil. New Phytologist 239, 752–765. 10.1111/nph.18943.

Joy, E.J.M., Ander, E.L., Young, S.D., Black, C.R., Watts, M.J., Chilimba, A.D.C., Chilima, B., Siyame, E.W.P., Kalimbira, A.A., Hurst, R., Fairweather-Tait, S.J., Stein, A.J., Gibson, R.S., White, P.J., Broadley, M.R., 2014. Dietary mineral supplies in Africa. Physiologia Plantarum 151, 208–229. 10.1111/ppl.12144.

Just, B.S., Marks, E.A.N., Roquer-Beni, L., Llenas, L., Ponsà, S., Vilaplana, R., 2024. Biofertilization increases soil organic carbon concentrations: Results of a meta-analysis. International Journal of Agricultural Sustainability 22, 2361578. 10.1080/14735903.2024.2361578.

Kaguongo, W., Maingi, G., Barker, I., Nganga, N., Guenthner, J., 2014. The value of seed potatoes from four systems in Kenya. American Journal of Potato Research 91, 109–118. 10.1007/s12230-013-9342-z.

Kalra, Y.P., 1998. Handbook of Reference Methods for Plant Analysis. CRC Press, Boca Raton, FL.

Kaminsky, L.M., Trexler, R.V., Malik, R.J., Hockett, K.L., Bell, T.H., 2019. The inherent conflicts in developing soil microbial inoculants. Trends in Biotechnology 37, 140–151. 10.1016/j.tibtech.2018.11.011.

Kesavardhini, K., Essa, I.M., Nayak, A.K., Gharban, H.A.J., Gayathri, K., Saranraj, P., 2025. Harnessing plant growth promoting rhizobacteria to bolster drought tolerance in plants. Discover Applied Sciences 8, 89. 10.1007/s42452-025-08095-7.

Koskey, G., Mburu, S.W., Njeru, E.M., Kimiti, J.M., Ombori, O., Maingi, J.M., 2017. Potential of native rhizobia in enhancing nitrogen fixation and yields of climbing beans (*Phaseolus vulgaris* L.) in contrasting environments of eastern Kenya. Frontiers in Plant Science 8, 443. 10.3389/fpls.2017.00443.

Lenth, R.V., Piaskowski, J., Banfai, B., Bolker, B., Buerkner, P., Ginè-Vázquez, I., Hervé, M., Jung, M., Love, J., Miguez, F., Riebl, H., Singmann, H., 2026. emmeans: Estimated Marginal Means, aka Least-Squares Means. R package.

Li, C., Chen, X., Jia, Z., Zhai, L., Zhang, B., Grüters, U., Ma, S., Qian, J., Liu, X., Zhang, J., Müller, C., 2024. Meta-analysis reveals the effects of microbial inoculants on the biomass and diversity of soil microbial communities. Nature Ecology & Evolution 8, 1270–1284. 10.1038/s41559-024-02437-1.

Liu, X., Mei, S., Salles, J.F., 2023. Inoculated microbial consortia perform better than single strains in living soil: A meta-analysis. Applied Soil Ecology 190, 105011. 10.1016/j.apsoil.2023.105011.

Lobell, D.B., Cassman, K.G., Field, C.B., 2009. Crop yield gaps: Their importance, magnitudes, and causes. Annual Review of Environment and Resources 34, 179–204. 10.1146/annurev.environ.041008.093740.

Luan, Y., Cui, X., Ferrat, M., 2013. Historical trends of food self-sufficiency in Africa. Food Security 5, 393–405. 10.1007/s12571-013-0260-1.

Masso, C., Mukhongo, R.W., Thuita, M., Abaidoo, R., Ulzen, J., Kariuki, G., Kalumuna, M., 2016. Biological inoculants for sustainable intensification of agriculture in sub-Saharan Africa smallholder farming systems, in: Lal, R., Kraybill, D., Hansen, D.O., Singh, B.R., Mosogoya, T., Eik, L.O. (Eds.), Climate Change and Multi-Dimensional Sustainability in African Agriculture. Springer International Publishing, Cham, pp. 639–658. 10.1007/978-3-319-41238-2_33.

Mishra, S., Choudhary, K.K., 2026. Elucidating the functional role of PGPR in modulating plant physiology for improved drought resilience and yield enhancement: A review. Journal of Soil Science and Plant Nutrition. 10.1007/s42729-026-03315-4.

Mukuralinda, A., Tenywa, J.S., Verchot, L., Obua, J., Nabahungu, N.L., Chianu, J.N., 2010. Phosphorus uptake and maize response to organic and inorganic fertilizer inputs in Rubona, Southern Province of Rwanda. Agroforestry Systems 80, 211–221. 10.1007/s10457-010-9324-9.

Myers, S.S., Zanobetti, A., Kloog, I., Huybers, P., Leakey, A.D.B., Bloom, A.J., Carlisle, E., Dietterich, L.H., Fitzgerald, G., Hasegawa, T., Holbrook, N.M., Nelson, R.L., Ottman, M.J., Raboy, V., Sakai, H., Sartor, K.A., Schwartz, J., Seneweera, S., Tausz, M., Usui, Y., 2014. Increasing CO₂ threatens human nutrition. Nature 510, 139–142. 10.1038/nature13179.

NASA Langley Research Center (LaRC) POWER Project, n.d. NASA/POWER CERES/MERRA-2 Native Resolution Daily Data. NASA Langley Research Center.

Nelson, D.W., Sommers, L.E., 1974. A rapid and accurate procedure for estimation of organic carbon in soils. Proceedings of the Indiana Academy of Science 84, 456–462.

Nie, M., Bell, C., Wallenstein, M.D., Pendall, E., 2015. Increased plant productivity and decreased microbial respiratory C loss by plant growth-promoting rhizobacteria under elevated CO₂. Scientific Reports 5, 9212. 10.1038/srep09212.

Nyaguthii, M.C., 2017. Soybean (Glycine max) Response to Rhizobia Inoculation as Influenced by Soil Nitrogen Levels. Kenyatta University.

Okereke, G.U., Onochie, C.C., Onukwo, A.U., Onyeagba, E., Ekejindu, G.O., 2000. Response of introduced *Bradyrhizobium* strains infecting a promiscuous soybean cultivar. World Journal of Microbiology and Biotechnology 16, 43–48. 10.1023/A:1008927327678.

Olanrewaju, O.S., Babalola, O.O., 2019. *Streptomyces*: Implications and interactions in plant growth promotion. Applied Microbiology and Biotechnology 103, 1179–1188. 10.1007/s00253-018-09577-y.

Plaza-Bonilla, D., Cantero-Martínez, C., Viñas, P., Álvaro-Fuentes, J., 2013. Soil aggregation and organic carbon protection in a no-tillage chronosequence under Mediterranean conditions. Geoderma 193–194, 76–82. 10.1016/j.geoderma.2012.10.022.

Rosen, C.J., Bierman, P.M., 2008. Potato yield and tuber set as affected by phosphorus fertilization. American Journal of Potato Research 85, 110–120. 10.1007/s12230-008-9001-y.

Schollenberger, C.J., Simon, R.H., 1945. Determination of exchange capacity and exchangeable bases in soil—ammonium acetate method. Soil Science 59, 13–24.

Schulte-Geldermann, E., Kakuhenzire, R., Sharma, K., Parker, M., 2022. Revolutionizing early generation seed potato in East Africa, in: Thiele, G., Friedmann, M., Campos, H., Polar, V., Bentley, J.W. (Eds.), Root, Tuber and Banana Food System Innovations: Value Creation for Inclusive Outcomes. Springer International Publishing, Cham, pp. 389–419. 10.1007/978-3-030-92022-7_13.

Schütz, L., Gattinger, A., Meier, M., Müller, A., Boller, T., Mäder, P., Mathimaran, N., 2018. Improving crop yield and nutrient use efficiency via biofertilization—A global meta-analysis. Frontiers in Plant Science 8, 2204. 10.3389/fpls.2017.02204.

Sorensen, P.O., Karaoz, U., Beller, H.R., Bill, M., Bouskill, N.J., Banfied, J.F., Chu, R.K., Hoyt, D.W., Eder, E., Eloe-Fadrosh, E., Sharrar, A., Tfaily, M.M., Toyoda, J., Tolic, N., Wang, S., Wong, A.R., Williams, K.H., Zhong, Y., Brodie, E.L., 2026. Multi-omics reveals nitrogen dynamics associated with soil microbial blooms during snowmelt. Nature Microbiology 11, 359–374. 10.1038/s41564-025-02213-2.

Sousa, J.A. de J., Olivares, F.L., 2016. Plant growth promotion by streptomycetes: Ecophysiology, mechanisms and applications. Chemical and Biological Technologies in Agriculture 3, 24. 10.1186/s40538-016-0073-5.

Sparks, A., 2018. nasapower: A NASA POWER global meteorology, surface solar energy and climatology data client for R. Journal of Open Source Software 3, 1035. 10.21105/joss.01035.

Sun, X., Tian, X., Jia, M., Hu, X., Zhang, C., Zhao, L., 2026. Phosphate-solubilizing bacteria: A review of diversity, mechanisms, and applications in sustainable agriculture. Frontiers in Microbiology 17, 1778470. 10.3389/fmicb.2026.1778470.

Thiex, N.J., Manson, H., Anderson, S., Persson, J.-Å., et al., 2002. Determination of crude protein in animal feed, forage, grain, and oilseeds by using block digestion with a copper catalyst and steam distillation into boric acid: Collaborative study. Journal of AOAC International 85, 309–317. 10.1093/jaoac/85.2.309.

Thilakarathna, M.S., Raizada, M.N., 2017. A meta-analysis of the effectiveness of diverse rhizobia inoculants on soybean traits under field conditions. Soil Biology and Biochemistry 105, 177–196. 10.1016/j.soilbio.2016.11.022.

Tsai, O., Stammer, A., Ruark, M.D., Wang, Y., 2025. Potato (*Solanum tuberosum* L.) responses to nitrogen fertilisation in different groundwater nitrate environments. Potato Research 68, 3949–3971. 10.1007/s11540-025-09898-2.

van Ittersum, M.K., van Bussel, L.G.J., Wolf, J., Grassini, P., van Wart, J., Guilpart, N., Claessens, L., de Groot, H., Wiebe, K., Mason-D’Croz, D., Yang, H., Boogaard, H., van Oort, P.A.J., van Loon, M.P., Saito, K., Adimo, O., Adjei-Nsiah, S., Agali, A., Bala, A., Chikowo, R., Kaizzi, K., Kouressy, K., Makoi, J.H.J.R., Ouattara, K., Tesfaye, K., Cassman, K.G., 2016. Can sub-Saharan Africa feed itself? Proceedings of the National Academy of Sciences 113, 14964–14969. 10.1073/pnas.1610359113.

Vanlauwe, B., Aihou, K., Aman, S., Iwuafor, E.N.O., Tossah, B.K., Diels, J., Sanginga, N., Lyasse, O., Merckx, R., Deckers, J., 2001. Maize yield as affected by organic inputs and urea in the West African Moist Savanna. Agronomy Journal 93: 1191–1199. 10.2134/agronj2001.1191.

Vanlauwe, B., Descheemaeker, K., Giller, K.E., Huising, J., Merckx, R., Nziguheba, G., Wendt, J., Zingore, S., 2015. Integrated soil fertility management in sub-Saharan Africa: Unravelling local adaptation. SOIL 1, 491–508. 10.5194/soil-1-491-2015.

Walinga, I., van der Lee, J.J., Houba, V.J.G., van Vark, W., Novozamsky, I., 1995. Digestion in tubes with H₂SO₄–salicylic acid–H₂O₂ and selenium and determination of Ca, K, Mg, N, Na, P, Zn, in: Walinga, I., van der Lee, J.J., Houba, V.J.G., van Vark, W., Novozamsky, I. (Eds.), Plant Analysis Manual. Springer Netherlands, Dordrecht, pp. 7–45. 10.1007/978-94-011-0203-2_2.

Wick, R.R., Judd, L.M., Cerdeira, L.T., Hawkey, J., Méric, G., Vezina, B., Wyres, K.L., Holt, K.E., 2021. Trycycler: Consensus long-read assemblies for bacterial genomes. Genome Biology 22, 266. 10.1186/s13059-021-02483-z.

Wickham, H., Averick, M., Bryan, J., Chang, W., McGowan, L.D., François, R., Grolemund, G., Hayes, A., Henry, L., Hester, J., Kuhn, M., Pedersen, T.L., Miller, E., Bache, S.M., Müller, K., Ooms, J., Robinson, D., Seidel, D.P., Spinu, V., Takahashi, K., Vaughan, D., Wilke, K., Woo, K., Yutani, H., 2019. Welcome to the tidyverse. Journal of Open Source Software 4, 1686. 10.21105/joss.01686.

Zhao, Q.-Y., Xu, S.-J., Zhang, W.-S., Zhang, Z., Yao, Z., Chen, X.-P., Zou, C.-Q., 2020. Identifying key drivers for geospatial variation of grain micronutrient concentrations in major maize production regions of China. Environmental Pollution 266, 115114. 10.1016/j.envpol.2020.115114.

