## Supplemental information for "Soil microbial inoculants augment fertilizer performance across contrasting cropping systems in Rwanda"

**Microbial inoculants increase potato and maize yield across Rwandan agricultural soils**

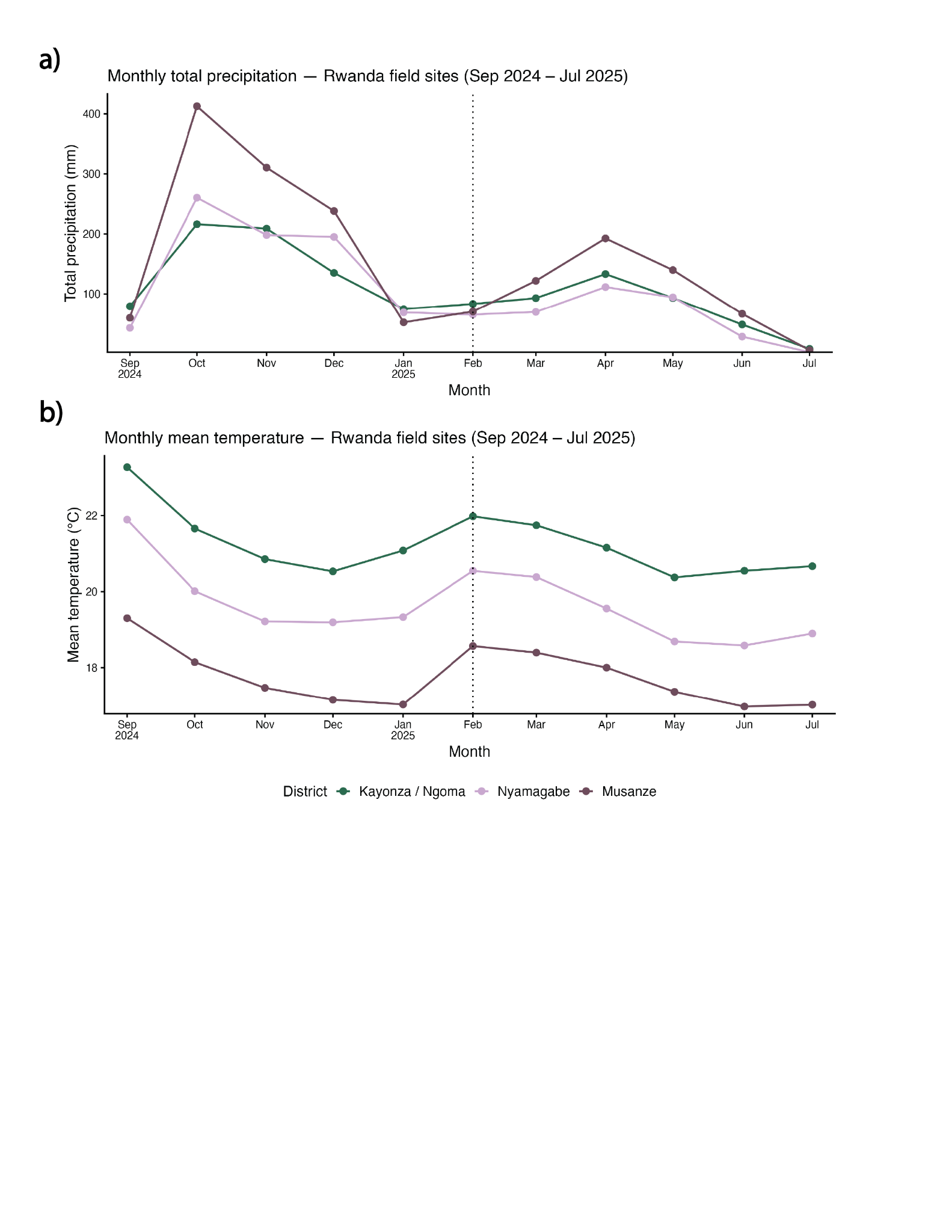

***Supplementary Figure 1.*** *Differences in a) total monthly precipitation and b) monthly mean temperature in seasons 2025A and 2025B across all four sites. In both panels, the dotted line along the x-axis at “February” indicates the transition between seasons 2025A and 2025B, and colors indicate sites. Given that Kayonza and Ngoma are only 27 km apart from each other, they are included in the same grid from the NASA POWER dataset used to extract daily temperature and precipitation values (Sparks, 2018; NASA Langley Research Center (LaRC) POWER Project). As such, they are represented by a single point and color.*

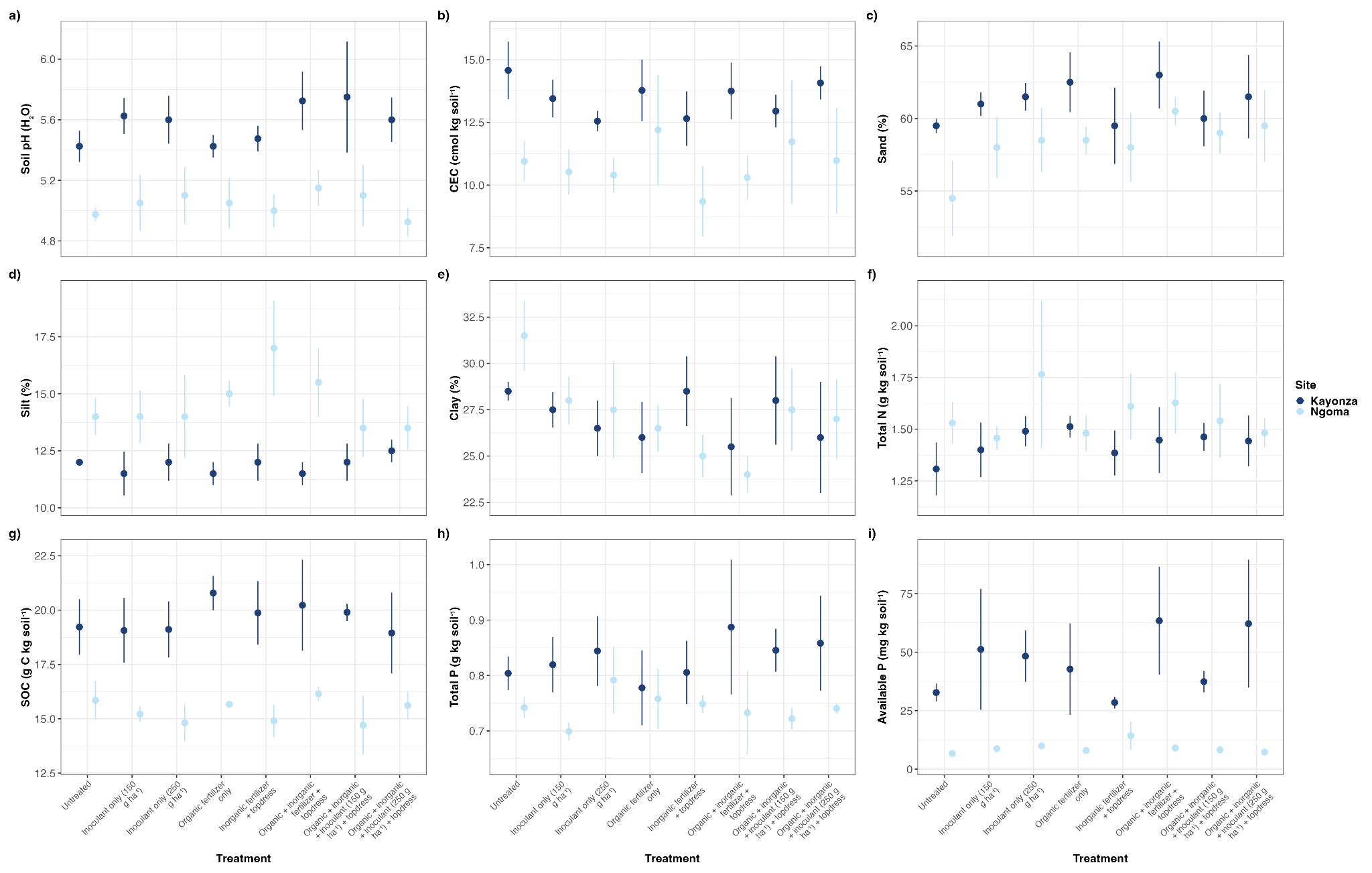

***Supplementary Figure 2.*** *Differences in soil properties among all maize sites. Across all panels, individual points and error bars represent group means and standard error, and different colors indicate site. Full statistical outputs are in Table S1.*

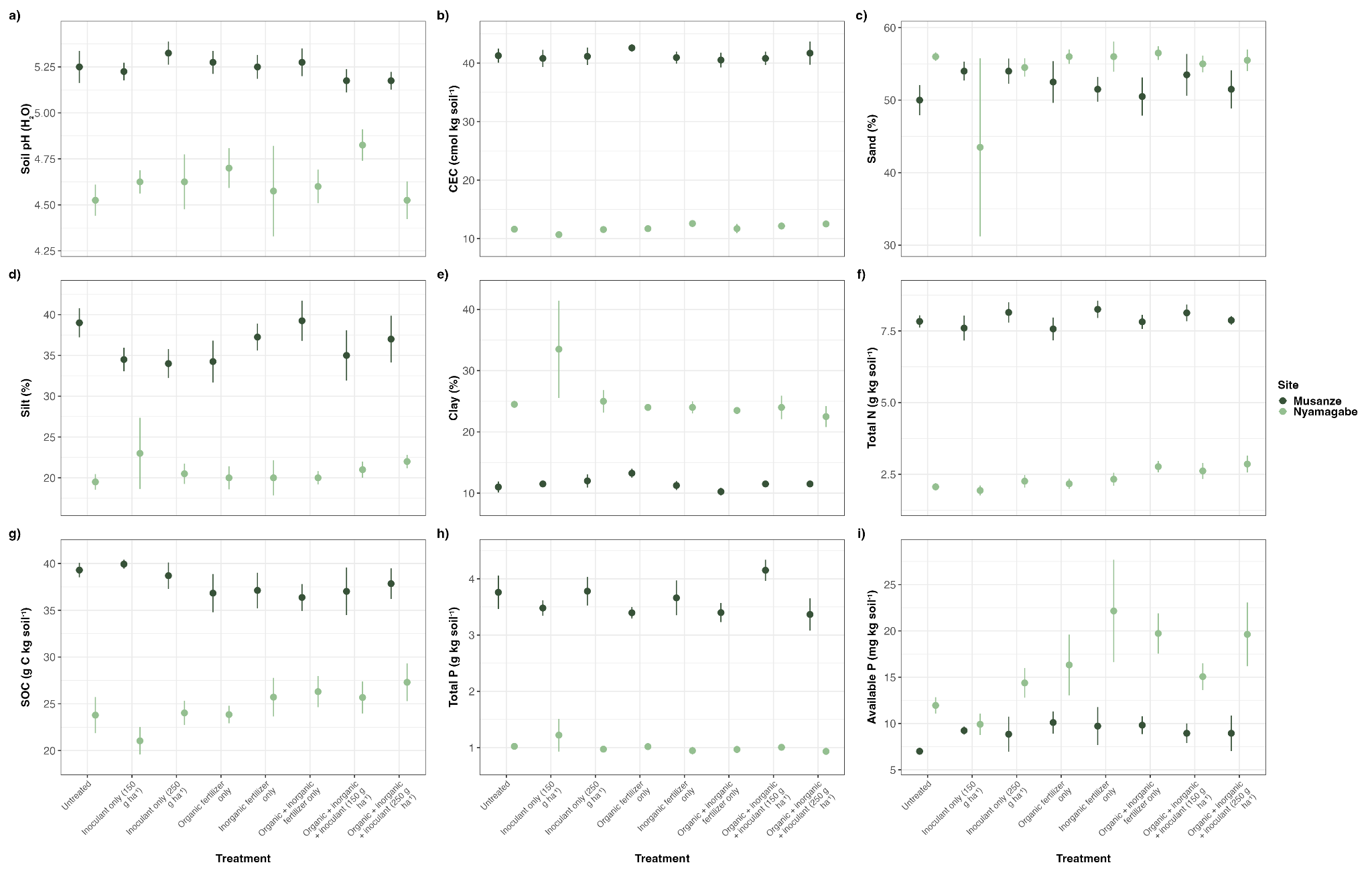

***Supplementary Figure 3.*** *Differences in soil properties among all potato sites. Across all panels, individual points and error bars represent group means and standard error, and different colors indicate site. Full statistical outputs are in Table S2.*

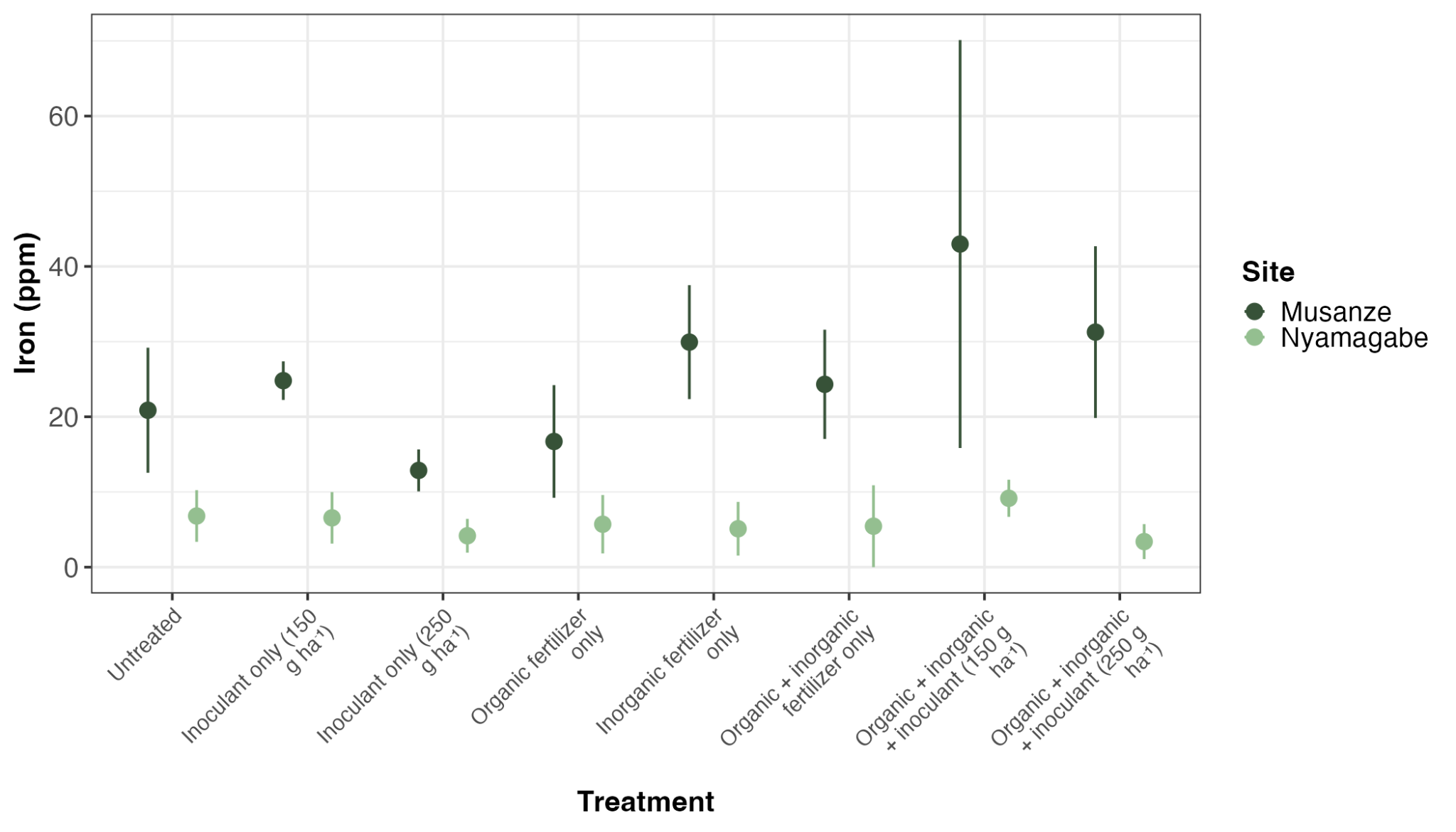

***Supplementary Figure 4.*** *Differences in potato iron concentration across treatments and sites, indicated by shades of green. Points represent treatment x site means and error bars indicate standard error (n=3). Full statistical outputs are in Table S16.*

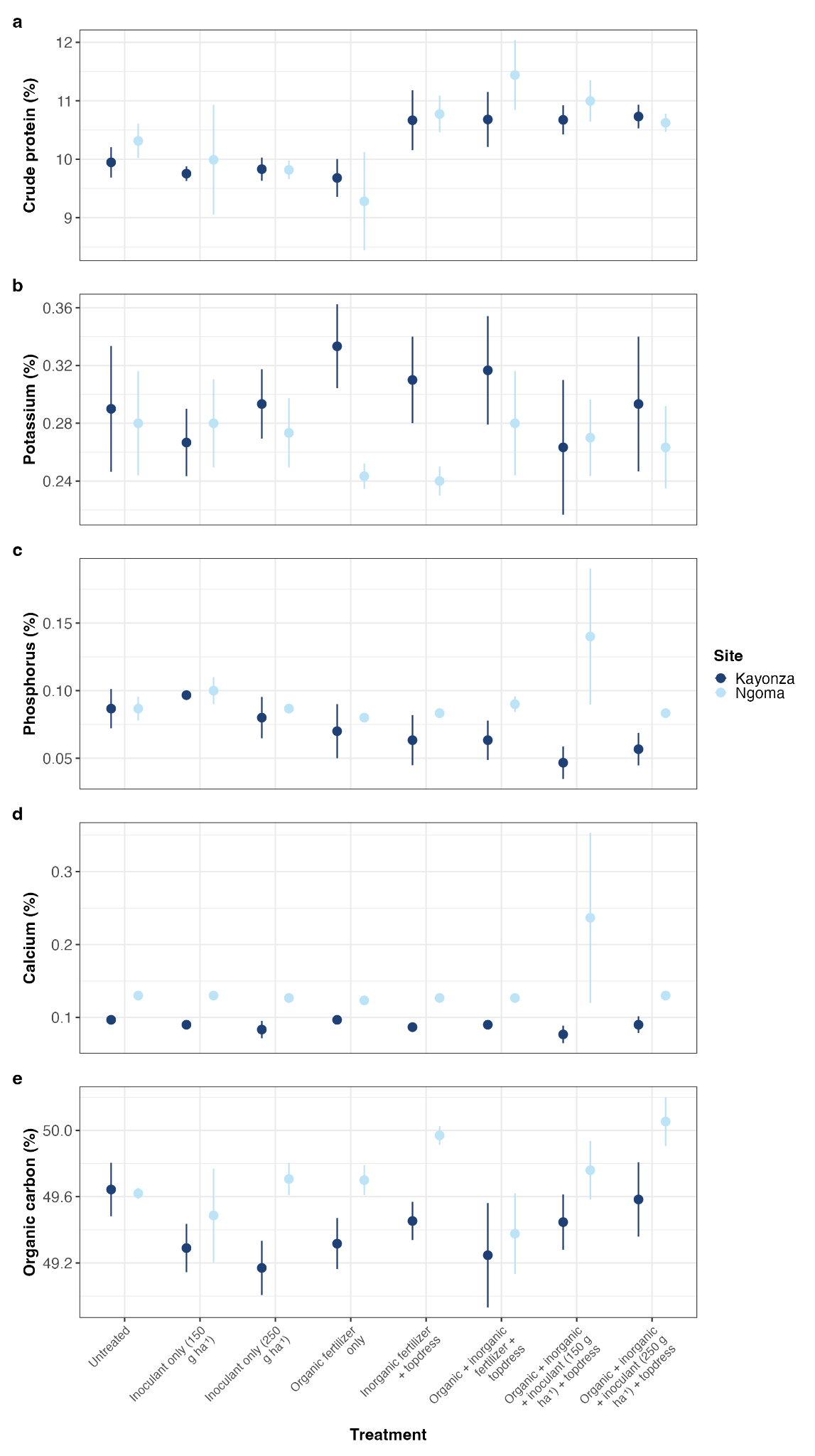

***Supplementary Figure 5.*** *Differences in concentration of five maize nutrients across treatments and sites, indicated by shades of blue. Points within each panel represent treatment x site means and error bars indicate standard error (n=3). Full statistical outputs are in Table S17.*

***Supplementary Table 1.*** *Bacterial taxa included in microbial inoculants applied across season 2025A and 2025B.*

| **ID** | **Species-level taxonomy** | **ID** | **Species-level taxonomy** |
| --- | --- | --- | --- |
| 1 | Bacillus licheniformis | 12 | Streptomyces globosus |
| 2 | Lysinibacillus sphaericus | 13 | Streptomyces griseoaurantiacus |
| 3 | Microbacterium oxydans | 14 | Streptomyces misionensis |
| 4 | Microbacterium paraoxydans | 15 | Streptomyces murinus |
| 5 | Paenibacillus cellulositrophicus | 16 | Streptomyces musisoli |
| 6 | Rhodococcus erythropolis | 17 | Streptomyces olivaceus |
| 7 | Streptomyces achromogenes | 18 | Streptomyces rochei |
| 8 | Streptomyces antibioticus | 19 | Streptomyces toxytricini |
| 9 | Streptomyces bacillari | 20 | Streptomyces sennicomposti |
| 10 | Streptomyces bikiniensis | 21* | Aneurinibacillus aneurinilyticus |
| 11 | Streptomyces echinatus | 22* | Bacillus velezensi |

**only included in formulations applied in season 2025B*

***Supplementary Table 2.*** *Analysis of Variance (ANOVA) results from linear models (~ Treatment * Site) run on soil variables measured in maize sites. *p<0.05; **p<0.01; ***p<0.001; ·p<0.1; n.s. non-significant.*

| **pH (measured in H_2_O)** | | | |
| --- | --- | --- | --- |
| **Fixed effect** | ***F* statistic** | **Degrees of freedom** | ***P* value** |
| Treatment | 0.60 | 7,32 | *p=*0.75 n.s. |
| Site | 42.4 | 1,32 | *p=*4.13e-8 *** |
| Treatment*Site | 0.2 | 7,32 | *p=*0.99 n.s. |
| **Cation exchange capacity (CEC)** | | | |
| **Fixed effect** | ***F* statistic** | **Degrees of freedom** | ***P* value** |
| Treatment | 0.52 | 7,32 | *p=*0.81 n.s. |
| Site | 17.04 | 1,32 | *p=*1.45e-4 *** |
| Treatment*Site | 0.245 | 7,32 | *p=*0.97 n.s. |
| **% sand** | | | |
| **Fixed effect** | ***F* statistic** | **Degrees of freedom** | ***P* value** |
| Treatment | 1.02 | 7,32 | *p=*0.43 n.s. |
| Site | 7.79 | 1,32 | *p=*0.008 ** |
| Treatment*Site | 0.22 | 7,32 | *p=*0.98 n.s. |
| **% silt** | | | |
| **Fixed effect** | ***F* statistic** | **Degrees of freedom** | ***P* value** |
| Treatment | 0.57 | 7,32 | *p=*0.77 n.s. |
| Site | 25.1 | 1,32 | *p=*7.79e-6 *** |
| Treatment*Site | 0.80 | 7,32 | *p=*0.59 n.s. |
| **% clay** | | | |
| **Fixed effect** | ***F* statistic** | **Degrees of freedom** | ***P* value** |
| Treatment | 1.26 | 7,32 | *p=*0.29 n.s. |
| Site | 0.004 | 1,32 | *p=*0.95 n.s. |
| Treatment*Site | 0.52 | 7,32 | *p=*0.82 n.s. |
| **Total nitrogen (N)** | | | |
| **Fixed effect** | ***F* statistic** | **Degrees of freedom** | ***P* value** |
| Treatment | 0.42 | 7,32 | *p=*0.88 n.s. |
| Site | 3.29 | 1,32 | *p=*0.076 · |
| Treatment*Site | 0.29 | 7,32 | *p=*0.96 n.s. |
| **Soil organic carbon (SOC)** | | | |
| **Fixed effect** | ***F* statistic** | **Degrees of freedom** | ***P* value** |
| Treatment | 0.33 | 7,32 | *p=*0.93 n.s. |
| Site | 56.18 | 1,32 | *p=*1.28e-9 *** |
| Treatment*Site | 0.218 | 7,32 | *p=*0.98 n.s. |
| **Total phosphorus (P)** | | | |
| **Fixed effect** | ***F* statistic** | **Degrees of freedom** | ***P* value** |
| Treatment | 0.27 | 7,32 | *p=*0.96 n.s. |
| Site | 9.57 | 1,32 | *p=*0.003 ** |
| Treatment*Site | 0.33 | 7,32 | *p=*0.94 n.s. |
| **Available P** | | | |
| **Fixed effect** | ***F* statistic** | **Degrees of freedom** | ***P* value** |
| Treatment | 0.47 | 7,32 | *p=*0.85 n.s. |
| Site | 34.2 | 1,32 | *p=*4.28e-7 *** |
| Treatment*Site | 0.61 | 7,32 | *p=*0.74 n.s. |

***Supplementary Table 3.*** *Analysis of Variance (ANOVA) results from linear models (~ Treatment * Site), and post-hoc pairwise comparisons run on soil variables measured in all potato sites. P-values from post-hoc analyses are corrected for multiple comparisons; only marginally and statistically significant contrasts are shown. *p<0.05; **p<0.01; ***p<0.001; ·p<0.1; n.s. non-significant.*

| **pH (measured in H_2_O)** | | | |
| --- | --- | --- | --- |
| **Fixed effect** | ***F* statistic** | **Degrees of freedom** | ***P* value** |
| Treatment | 0.51 | 7,32 | *p=*0.82 n.s. |
| Site | 147.75 | 1,32 | *p=*2.92e-16 *** |
| Treatment*Site | 0.69 | 7,32 | *p=*0.68 n.s. |
| **Cation exchange capacity (CEC)** | | | |
| **Fixed effect** | ***F* statistic** | **Degrees of freedom** | ***P* value** |
| Treatment | 0.46 | 7,32 | *p=*0.86 n.s. |
| Site | 3470.71 | 1,32 | *p=*1.99e-46 *** |
| Treatment*Site | 0.35 | 7,32 | *p=*0.93 n.s. |
| **% sand** | | | |
| **Fixed effect** | ***F* statistic** | **Degrees of freedom** | ***P* value** |
| Treatment | 0.53 | 7,32 | *p=*0.81 n.s. |
| Site | 1.17 | 1,32 | *p=*0.28 n.s. |
| Treatment*Site | 1.13 | 7,32 | *p=*0.36 n.s. |
| **% silt** | | | |
| **Fixed effect** | ***F* statistic** | **Degrees of freedom** | ***P* value** |
| Treatment | 0.42 | 7,32 | *p=*0.89 n.s. |
| Site | 214.60 | 1,32 | *p=*2.44-19 *** |
| Treatment*Site | 0.91 | 7,32 | *p=*0.51 n.s. |
| **% clay** | | | |
| **Fixed effect** | ***F* statistic** | **Degrees of freedom** | ***P* value** |
| Treatment | 1.31 | 7,32 | *p=*0.26 n.s. |
| Site | 150.78 | 1,32 | *p=*2.02e-16 *** |
| Treatment*Site | 1.28 | 7,32 | *p=*0.28 n.s. |
| **Total nitrogen (N)** | | | |
| **Fixed effect** | ***F* statistic** | **Degrees of freedom** | ***P* value** |
| Treatment | 1.59 | 7,32 | *p=*0.16 n.s. |
| Site | 1687.49 | 1,32 | *p=*4.66e-39 *** |
| Treatment*Site | 0.86 | 7,32 | *p=*0.54 n.s. |
| **Soil organic carbon (SOC)** | | | |
| **Fixed effect** | ***F* statistic** | **Degrees of freedom** | ***P* value** |
| Treatment | 0.39 | 7,32 | *p=*0.93 n.s. |
| Site | 251.034 | 1,32 | *p=*1.07e-20 *** |
| Treatment*Site | 1.64 | 7,32 | *p=*0.15 n.s. |
| **Total phosphorus (P)** | | | |
| **Fixed effect** | ***F* statistic** | **Degrees of freedom** | ***P* value** |
| Treatment | 1.19 | 7,32 | *p=*0.32 n.s. |
| Site | 834.29 | 1,32 | *p=*5.32e-32 *** |
| Treatment*Site | 1.3 | 7,32 | *p=*0.27 n.s. |
| **Available P ANOVA** | | | |
| **Fixed effect** | ***F* statistic** | **Degrees of freedom** | ***P* value** |
| Treatment | 2.25 | 7,32 | *p=*0.046 * |
| Site | 40.00 | 1,32 | *p=*7.98e-8 *** |
| Treatment*Site | 1.42 | 7,32 | *p=*0.22 n.s. |
| **Available P post-hoc pairwise comparisons** | | | |
| **ANOVA fixed effect** | **Contrast** | **Estimate** | **Corrected *P* value** |
| Treatment | Untreated - Inorganic | -6.46 | *p=*0.091 · |
|  | Inoc (low) - Inorganic | -6.36 | *p=*0.091 · |

***Supplementary Table 4.*** *Results of analyses of variance (ANOVAs) from linear models (~ Treatment * Site), and post-hoc pairwise comparisons run on small-grade potato tuber yield. Data were subset by Season prior to running models. P-values from post-hoc analyses are corrected for multiple comparisons; only marginally and statistically significant contrasts are shown. *p<0.05; **p<0.01; ***p<0.001; ·p<0.1; n.s. non-significant.*

| **Season 2025A ANOVA results** | | | |
| --- | --- | --- | --- |
| **Fixed effect** | **F statistic** | **Degrees of freedom** | ***P* value** |
| Treatment | 44.51 | 7,32 | *p=*1.02e-14 *** |
| Site | 969.73 | 1,32 | *p=*1.67e-25 *** |
| Treatment * Site | 13.53 | 7,32 | *p=*5.55e-8 *** |
| **Season 2025A post-hoc pairwise comparisons** | | | |
| **ANOVA fixed effect** | **Contrast** | **Estimate** | ***P* value** |
| Treatment * Site:  Contrasts within Musanze | Untreated - Inoc (low) | -10.25 | *p=*6.35e-10 *** |
|  | Untreated - Inoc (high) | -7.94 | *p=*1.03e-7 *** |
|  | Untreated - Organic + Inorganic + Inoc (low) | -12.17 | *p=*1.40e-11 *** |
|  | Untreated - Organic + Inorganic + Inoc (high) | -18.95 | *p=*3.0e-16 *** |
|  | Untreated - Organic + Inorganic | -14.54 | *p=*2.57e-13 *** |
|  | Untreated - Organic | -8.94 | *p=*1.07e-8 *** |
|  | Untreated - Inorganic | -14.18 | *p=*3.37e-13 *** |
|  | Inoc (low) - Organic + Inorganic + Inoc (high) | -8.7 | *p=*1.71e-8 *** |
|  | Inoc (low) - Organic + Inorganic | -4.29 | *p=*7.53e-4 *** |
|  | Inoc (low) - Inorganic | -3.93 | *p=*0.002 ** |
|  | Inoc (high) - Organic + Inorganic + Inoc (low) | -4.23 | *p=*8.22e-4 *** |
|  | Inoc (high) - Organic + Inorganic + Inoc (high) | -11.01 | *p=*1.37e-10 *** |
|  | Inoc (high) - Organic + Inorganic | -6.6 | *p=*2.69e-6 *** |
|  | Inoc (high) - Inorganic | -6.24 | *p=*6.39e-6 *** |
|  | Organic + Inorganic + Inoc (low) - Organic + Inorganic + Inoc (high) | -6.78 | *p=*1.86e-6 *** |
|  | Organic + Inorganic + Inoc (low) - Organic + Inorganic | -2.37 | *p=*0.05 * |
|  | Organic + Inorganic + Inoc (low) - Organic | 3.23 | *p=*0.008 ** |
|  | Organic + Inorganic + Inoc (high) - Organic + Inorganic | 4.41 | *p=*5.93e-4 *** |
|  | Organic + Inorganic + Inoc (high) - Organic | 10.01 | *p=*9.48e-10 *** |
|  | Organic + Inorganic + Inoc (high) - Inorganic | 4.77 | *p=*2.50e-4 *** |
|  | Organic + Inorganic - Organic | 5.60 | *p=*3.21e-5 *** |
|  | Organic - Inorganic | -5.24 | *p=*7.71e-5 *** |
| Treatment * Site:  Contrasts within Nyamagabe | Untreated - Organic + Inorganic + Inoc (low) | -4.77 | *p=*0.002 ** |
|  | Untreated - Organic + Inorganic + Inoc (high) | -5.43 | *p=*7.02e-4 *** |
|  | Untreated - Organic + Inorganic | -3.53 | *p=*0.012 * |
|  | Untreated - Inorganic | -3.48 | *p=*0.012 * |
|  | Inoc (low) - Organic + Inorganic + Inoc (low) | -3.36 | *p=*0.014 * |
|  | Inoc (low) - Organic + Inorganic + Inoc (high) | -4.02 | *p=*0.007 ** |
|  | Inoc (high) - Organic + Inorganic + Inoc (low) | -2.85 | *p=*0.041 * |
|  | Inoc (high) - Organic + Inorganic + Inoc (high) | -3.51 | *p=*0.012 * |
|  | Organic + Inorganic + Inoc (low) - Organic | 3.77 | *p=*0.01 ** |
|  | Organic + Inorganic + Inoc (high) - Organic | 4.43 | *p=*0.003 ** |
| Treatment * Site:  Contrasts between Musanze - Nyamagabe | Untreated | 3.98 | *p=*0.001 ** |
|  | Inoc (low) | 12.82 | *p=*5.19e-13 *** |
|  | Inoc (high) | 10.0 | *p=*2.45e-10 *** |
|  | Organic + Inorganic + Inoc (low) | 11.38 | *p=*1.08e-11 *** |
|  | Organic + Inorganic + Inoc (high) | 17.5 | *p=*1.06e-16 *** |
|  | Organic + Inorganic | 14.99 | *p=*7.98e-15 *** |
|  | Organic | 11.92 | *p=*3.41e-12 *** |
|  | Inorganic | 14.68 | *p=*1.40e-14 *** |
| **Season 2025B ANOVA results** | | | |
| **Fixed effect** | **F statistic** | **Degrees of freedom** | ***P* value** |
| Treatment | 21.65 | 7,32 | *p=*1.86e-10 *** |
| Site | 174.6 | 1,32 | *p=*1.66e-14 *** |
| Treatment*Site | 2.83 | 7,32 | *p=*0.021 * |
| **Season 2025B post-hoc pairwise comparisons** | | | |
| **ANOVA fixed effect** | **Contrast** | **Estimate** | ***P* value** |
| Treatment * Site:  Contrasts within Musanze | Untreated - Inoc (high) | -2.07 | *p=*0.012 * |
|  | Untreated - Organic + Inorganic + Inoc (low) | -2.83 | *p=*0.001 ** |
|  | Untreated - Organic + Inorganic + Inoc (high) | -4.96 | *p=*1.08e-6 *** |
|  | Untreated - Organic + Inorganic | -2.19 | *p=*0.009 ** |
|  | Inoc (low) - Inoc (high) | -1.87 | *p=*0.022 * |
|  | Inoc (low) - Organic + Inorganic + Inoc (low) | -2.63 | *p=*0.002 ** |
|  | Inoc (low) - Organic + Inorganic + Inoc (high) | -4.75 | *p=*1.23e-6 *** |
|  | Inoc (low) - Organic + Inorganic | -1.99 | *p=*0.015 * |
|  | Inoc (high) - Organic + Inorganic + Inoc (high) | -2.88 | *p=*0.001 ** |
|  | Inoc (high) - Organic | 1.69 | *p=*0.035 * |
|  | Organic + Inorganic + Inoc (low) - Organic + Inorganic + Inoc (high) | -2.12 | *p=*0.011 * |
|  | Organic + Inorganic + Inoc (low) - Organic | 2.45 | *p=*0.004 ** |
|  | Organic + Inorganic + Inoc (high) - Organic + Inorganic | 2.77 | *p=*0.001 ** |
|  | Organic + Inorganic + Inoc (high) - Organic | 4.57 | *p=*1.72e-6 *** |
|  | Organic + Inorganic + Inoc (high) - Inorganic | 3.67 | *p=*5.65e-5 *** |
|  | Organic + Inorganic - Organic | 1.81 | *p=*0.025 * |
| Treatment * Site:  Contrasts within Nyamagabe | Untreated - Inoc (low) | -1.7 | *p=*0.042 * |
|  | Untreated - Organic + Inorganic + Inoc (low) | -5.33 | *p=*2.45e-7 *** |
|  | Untreated - Organic + Inorganic + Inoc (high) | -4.07 | *p=*1.07e-5 *** |
|  | Untreated - Organic + Inorganic | -1.61 | *p=*0.053 · |
|  | Inoc (low) - Organic + Inorganic + Inoc (low) | -3.63 | *p=*3.82e-5 *** |
|  | Inoc (low) - Organic + Inorganic + Inoc (high) | -2.37 | *p=*0.004 ** |
|  | Inoc (high) - Organic + Inorganic + Inoc (low) | -3.78 | *p=*2.85e-5 *** |
|  | Inoc (high) - Organic + Inorganic + Inoc (high) | -2.52 | *p=*0.003 ** |
|  | Organic + Inorganic + Inoc (low) - Organic + Inorganic | 3.72 | *p=*3.10e-5 *** |
|  | Organic + Inorganic + Inoc (low) - Organic | 4.4 | *p=*5.37e-6 *** |
|  | Organic + Inorganic + Inoc (low) - Inorganic | 4.17 | *p=*9.09e-6 *** |
|  | Organic + Inorganic + Inoc (high) - Organic + Inorganic | 2.46 | *p=*0.003 ** |
|  | Organic + Inorganic + Inoc (high) - Organic | 3.14 | *p=*2.65e-4 *** |
|  | Organic + Inorganic + Inoc (high) - Inorganic | 2.91 | *p=*5.94e-4 *** |
| Treatment * Site:  Contrasts between Musanze - Nyamagabe | Untreated | 3.53 | *p=*1.46e-5 *** |
|  | Inoc (low) | 2.04 | *p=*0.006 ** |
|  | Inoc (high) | 4.06 | *p=*1.59e-6 *** |
|  | Organic + Inorganic + Inoc (high) | 4.42 | *p=*3.48e-7 *** |
|  | Organic + Inorganic | 4.11 | *p=*1.27e-6 *** |
|  | Organic | 2.98 | *p=*1.43e-4 *** |
|  | Inorganic | 3.66 | *p=*8.31e-6 *** |

***Supplementary Table 5.*** *Results of analyses of variance (ANOVAs) from linear models (~ Treatment * Site), and post-hoc pairwise comparisons run on large-grade potato tuber yield. Data were subset by Season prior to running models. P-values from post-hoc analyses are corrected for multiple comparisons; only marginally and statistically significant contrasts are shown. *p<0.05; **p<0.01; ***p<0.001; ·p<0.1; n.s. non-significant.*

| **Season 2025A ANOVA results** | | | |
| --- | --- | --- | --- |
| **Fixed effect** | **F statistic** | **Degrees of freedom** | ***P* value** |
| Treatment | 88.3 | 7,32 | *p=*4.21e-19 *** |
| Site | 39.57 | 1,32 | *p=*4.7e-7 *** |
| Treatment*Site | 10.73 | 7,32 | *p=*7.02e-7 *** |
| **Season 2025A post-hoc pairwise comparisons** | | | |
| **ANOVA fixed effect** | **Contrast** | **Estimate** | ***P* value** |
| Treatment * Site:  Contrasts within Musanze | Untreated - Inoc (low) | -3.16 | *p=*0.003 ** |
|  | Untreated - Inoc (high) | -4.47 | *p=*7.87e-5 *** |
|  | Untreated - Organic + Inorganic + Inoc (low) | -17.53 | *p=*4.36e-17 *** |
|  | Untreated - Organic + Inorganic + Inoc (high) | -9.34 | *p=*1.41e-10 *** |
|  | Untreated - Organic + Inorganic | -12.16 | *p=*2.17e-13 *** |
|  | Untreated - Organic | -3.22 | *p=*0.003 ** |
|  | Untreated - Inorganic | -2.56 | *p=*0.015 * |
|  | Inoc (low) - Organic + Inorganic + Inoc (low) | -14.37 | *p=*3.7e-15 *** |
|  | Inoc (low) - Organic + Inorganic + Inoc (high) | -6.18 | *p=*5.73e-7 *** |
|  | Inoc (low) - Organic + Inorganic | -9.0 | *p=*3.02e-10 *** |
|  | Inoc (high) - Organic + Inorganic + Inoc (low) | -13.06 | *p=*3.80e-14 *** |
|  | Inoc (high) - Organic + Inorganic + Inoc (high) | -4.87 | *p=*2.53e-5 *** |
|  | Inoc (high) - Organic + Inorganic | -7.69 | *p=*8.19e-9 *** |
|  | Inoc (high) - Inorganic | 1.913 | *p=*0.066 · |
|  | Organic + Inorganic + Inoc (low) - Organic + Inorganic + Inoc (high) | 8.19 | *p=*2.21e-9 *** |
|  | Organic + Inorganic + Inoc (low) - Organic + Inorganic | 5.37 | *p=*5.86e-6 *** |
|  | Organic + Inorganic + Inoc (low) - Organic | 14.31 | *p=*3.7e-15 *** |
|  | Organic + Inorganic + Inoc (low) - Inorganic | 14.97 | *p=*2.08e-15 *** |
|  | Organic + Inorganic + Inoc (high) - Organic + Inorganic | -2.82 | *p=*0.008 ** |
|  | Organic + Inorganic + Inoc (high) - Organic | 6.12 | *p=*6.34e-7 *** |
|  | Organic + Inorganic + Inoc (high) - Inorganic | 6.78 | *p=*1.04e-7 *** |
|  | Organic + Inorganic - Organic | 8.95 | *p=*3.16e-10 *** |
|  | Organic + Inorganic - Inorganic | 9.6 | *p=*8.18e-11 *** |
| Treatment * Site:  Contrasts within Nyamagabe | Untreated - Inoc (low) | -4.76 | *p=*7.35e-5 *** |
|  | Untreated - Inoc (high) | -3.16 | *p=*0.004 ** |
|  | Untreated - Organic + Inorganic + Inoc (low) | -10.49 | *p=*6.29e-11 *** |
|  | Untreated - Organic + Inorganic + Inoc (high) | -7.10 | *p=*1.32e-7 *** |
|  | Untreated - Organic + Inorganic | -7.58 | *p=*6.64e-8 *** |
|  | Untreated - Organic | -3.69 | *p=*0.001 ** |
|  | Untreated - Inorganic | -4.57 | *p=*1.19e-4 *** |
|  | Inoc (low) - Organic + Inorganic + Inoc (low) | -5.73 | *p=*4.49e-6 *** |
|  | Inoc (low) - Organic + Inorganic + Inoc (high) | -2.34 | *p=*0.027 * |
|  | Inoc (low) - Organic + Inorganic | -2.82 | *p=*0.009 ** |
|  | Inoc (high) - Organic + Inorganic + Inoc (low) | -7.33 | *p=*9.04e-8 *** |
|  | Inoc (high) - Organic + Inorganic + Inoc (high) | -3.95 | *p=*6.23e-4 *** |
|  | Inoc (high) - Organic + Inorganic | -4.43 | *p=*1.63e-4 *** |
|  | Organic + Inorganic + Inoc (low) - Organic + Inorganic + Inoc (high) | 3.39 | *p=*0.002 ** |
|  | Organic + Inorganic + Inoc (low) - Organic + Inorganic | 2.91 | *p=*0.007 ** |
|  | Organic + Inorganic + Inoc (low) - Organic | 6.80 | *p=*2.53e-7 *** |
|  | Organic + Inorganic + Inoc (low) - Inorganic | 5.92 | *p=*2.89e-6 *** |
|  | Organic + Inorganic + Inoc (high) - Organic | 3.42 | *p=*0.002 ** |
|  | Organic + Inorganic + Inoc (high) - Inorganic | 2.54 | *p=*0.017 * |
|  | Organic + Inorganic - Organic | 3.9 | *p=*6.62e-4 *** |
|  | Organic + Inorganic - Inorganic | 3.02 | *p=*0.006 ** |
| Treatment * Site:  Contrasts between Musanze - Nyamagabe | Inoc (high) | 2.06 | *p=*0.039 * |
|  | Organic + Inorganic + Inoc (low) | 7.78 | *p=*2.72e-9 *** |
|  | Organic + Inorganic + Inoc (high) | 2.98 | *p=*0.004 ** |
|  | Organic + Inorganic | 5.32 | *p=*3.86e-6 *** |
| **Season 2025B ANOVA results** | | | |
| **Fixed effect** | **F statistic** | **Degrees of freedom** | ***P* value** |
| Treatment | 26.10 | 7,32 | *p=*1.62e-11 *** |
| Site | 204.59 | 1,32 | *p=*1.88e-15 *** |
| Treatment*Site | 9.74 | 7,32 | *p=*1.91e-6*** |
| **Season 2025B post-hoc pairwise comparisons** | | | |
| **ANOVA fixed effect** | **Contrast** | **Estimate** | ***P* value** |
| Treatment * Site:  Contrasts within Musanze | Untreated - Inoc (low) | -9.5 | *p=*1.48e-9 *** |
|  | Untreated - Inoc (high) | -5.45 | *p=*2.63e-5 *** |
|  | Untreated - Organic + Inorganic + Inoc (low) | -10.81 | *p=*1.24e-10 *** |
|  | Untreated - Organic + Inorganic + Inoc (high) | -7.1 | *p=*3.36e-7 *** |
|  | Untreated - Organic + Inorganic | -8.54 | *p=*1.15e-8 *** |
|  | Untreated - Organic | -6.17 | *p=*3.85e-6 *** |
|  | Untreated - Inorganic | -7.9 | *p=*4.69e-8 *** |
|  | Inoc (low) - Inoc (high) | 4.05 | *p=*9.79e-4 *** |
|  | Inoc (low) - Organic + Inorganic + Inoc (high) | 2.4 | *p=*0.042 * |
|  | Inoc (low) - Organic | 3.33 | *p=*0.006 ** |
|  | Inoc (high) - Organic + Inorganic + Inoc (low) | -5.37 | *p=*2.90e-5 *** |
|  | Inoc (high) - Organic + Inorganic | -3.09 | *p=*0.01 ** |
|  | Inoc (high) - Inorganic | -2.45 | *p=*0.04 * |
|  | Organic + Inorganic + Inoc (low) - Organic + Inorganic + Inoc (high) | 3.71 | *p=*0.002 ** |
|  | Organic + Inorganic + Inoc (low) - Organic + Inorganic | 2.28 | *p=*0.049 * |
|  | Organic + Inorganic + Inoc (low) - Organic | 4.64 | *p=*2.05e-4 *** |
|  | Organic + Inorganic + Inoc (low) - Inorganic | 2.91 | *p=*0.014 * |
|  | Organic + Inorganic - Organic | 2.37 | *p=*0.043 * |
| Treatment * Site:  Contrasts within Nyamagabe | Untreated - Organic + Inorganic + Inoc (low) | -6.21 | *p=*6.88e-6 *** |
|  | Untreated - Organic + Inorganic + Inoc (high) | -5.59 | *p=*1.74e-5 *** |
|  | Untreated - Organic + Inorganic | -4.41 | *p=*3.28e-4 *** |
|  | Untreated - Organic | -2.95 | *p=*0.011 * |
|  | Inoc (low) - Organic + Inorganic + Inoc (low) | -6.53 | *p=*6.88e-6 *** |
|  | Inoc (low) - Organic + Inorganic + Inoc (high) | -5.91 | *p=*9.73e-6 *** |
|  | Inoc (low) - Organic + Inorganic | -4.73 | *p=*1.61e-4 *** |
|  | Inoc (low) - Organic | -3.27 | *p=*0.006 ** |
|  | Inoc (high) - Organic + Inorganic + Inoc (low) | -6.24 | *p=*6.88e-6 *** |
|  | Inoc (high) - Organic + Inorganic + Inoc (high) | -5.62 | *p=*1.74e-5 *** |
|  | Inoc (high) - Organic + Inorganic | -4.44 | *p=*3.28e-4 *** |
|  | Inoc (high) - Organic | -2.98 | *p=*0.011 * |
|  | Organic + Inorganic + Inoc (low) - Organic | 3.26 | *p=*0.006 ** |
|  | Organic + Inorganic + Inoc (low) - Inorganic | 5.93 | *p=*9.73e-6 *** |
|  | Organic + Inorganic + Inoc (high) - Organic | 2.64 | *p=*0.022 * |
|  | Organic + Inorganic + Inoc (high) - Inorganic | 5.31 | *p=*3.44e-5 *** |
|  | Organic + Inorganic - Inorganic | 4.12 | *p=*6.65e-4 *** |
|  | Organic - Inorganic | 2.67 | *p=*0.021 * |
| Treatment * Site:  Contrasts between Musanze - Nyamagabe | Inoc (low) | 10.39 | *p=*1.20e-11 *** |
|  | Inoc (high) | 6.05 | *p=*1.16e-6 *** |
|  | Organic + Inorganic + Inoc (low) | 5.18 | *p=*1.43e-5 *** |
|  | Organic + Inorganic + Inoc (high) | 2.08 | *p=*0.048 * |
|  | Organic + Inorganic | 4.70 | *p=*5.55e-5 *** |
|  | Organic | 3.79 | *p=*7.15e-4 *** |
|  | Inorganic | 8.19 | *p=*3.08e-9 *** |

***Supplementary Table 6.*** *Results of analyses of variance (ANOVAs) from linear models (~ Treatment * Site), and post-hoc pairwise comparisons run on total potato tuber yield. Data were subset by Season prior to running models. P-values from post-hoc analyses are corrected for multiple comparisons; only marginally and statistically significant contrasts are shown. *p<0.05; **p<0.01; ***p<0.001; ·p<0.1; n.s. non-significant.*

| **Season 2025A ANOVA results** | | | |
| --- | --- | --- | --- |
| **Fixed effect** | **F statistic** | **Degrees of freedom** | ***P* value** |
| Treatment | 146.23 | 7,32 | *p=*1.89e-22 *** |
| Site | 1051.56 | 1,32 | *p=*4.75e-26 *** |
| Treatment*Site | 18.89 | 7,32 | *p=*1.04e-9 *** |
| **Season 2025A post-hoc pairwise comparisons** | | | |
| **ANOVA fixed effect** | **Contrast** | **Estimate** | ***P* value** |
| Treatment * Site:  Contrasts within Musanze | Untreated - Inoc (low) | -13.41 | *p=*7.94e-12 *** |
|  | Untreated - Inoc (high) | -12.41 | *p=*4.40e-11 *** |
|  | Untreated - Organic + Inorganic + Inoc (low) | -29.7 | *p=*1.73e-20 *** |
|  | Untreated - Organic + Inorganic + Inoc (high) | -28.29 | *p=*3.78e-20 *** |
|  | Untreated - Organic + Inorganic | -26.7 | *p=*1.43e-19 *** |
|  | Untreated - Organic | -12.15 | *p=*6.88e-11 *** |
|  | Untreated - Inorganic | -16.74 | *p=*4.99e-14 *** |
|  | Inoc (low) - Organic + Inorganic + Inoc (low) | -16.29 | *p=*8.87e-14 *** |
|  | Inoc (low) - Organic + Inorganic + Inoc (high) | -14.88 | *p=*7.01e-13 *** |
|  | Inoc (low) - Organic + Inorganic | -13.29 | *p=*9.14e-12 *** |
|  | Inoc (low) - Inorganic | -3.33 | *p=*0.015 * |
|  | Inoc (high) - Organic + Inorganic + Inoc (low) | -17.29 | *p=*2.45e-14 *** |
|  | Inoc (high) - Organic + Inorganic + Inoc (high) | -15.88 | *p=*1.39e-13 *** |
|  | Inoc (high) - Organic + Inorganic | -14.29 | *p=*1.68e-12 *** |
|  | Inoc (high) - Inorganic | -4.33 | *p=*0.002 ** |
|  | Organic + Inorganic + Inoc (low) - Organic + Inorganic | 3 | *p=*0.027 * |
|  | Organic + Inorganic + Inoc (low) - Organic | 17.55 | *p=*2.04e-14 *** |
|  | Organic + Inorganic + Inoc (low) - Inorganic | 12.96 | *p=*1.60e-11 *** |
|  | Organic + Inorganic + Inoc (high) - Organic | 16.13 | *p=*1.01e-13 *** |
|  | Organic + Inorganic + Inoc (high) - Inorganic | 11.55 | *p=*2.19e-10 *** |
|  | Organic + Inorganic - Organic | 14.55 | *p=*1.15e-12 *** |
|  | Organic + Inorganic - Inorganic | 9.96 | *p=*5.87e-09 *** |
|  | Organic - Inorganic | -4.58 | *p=*0.001 ** |
| Treatment * Site:  Contrasts within Nyamagabe | Untreated - Inoc (low) | -6.17 | *p=*4.98e-5 *** |
|  | Untreated - Inoc (high) | -5.08 | *p=*5.32e-4 *** |
|  | Untreated - Organic + Inorganic + Inoc (low) | -15.26 | *p=*3.60e-12 *** |
|  | Untreated - Organic + Inorganic + Inoc (high) | -12.54 | *p=*2.75e-10 *** |
|  | Untreated - Organic + Inorganic | -11.11 | *p=*3.28e-9 *** |
|  | Untreated - Organic | -4.68 | *p=*0.001 ** |
|  | Untreated - Inorganic | -8.04 | *p=*1.17e-6 *** |
|  | Inoc (low) - Organic + Inorganic + Inoc (low) | -9.09 | *p=*1.26e-7 *** |
|  | Inoc (low) - Organic + Inorganic + Inoc (high) | -6.37 | *p=*3.35e-5 *** |
|  | Inoc (low) - Organic + Inorganic | -4.94 | *p=*6.78e-4 *** |
|  | Inoc (high) - Organic + Inorganic + Inoc (low) | -10.18 | *p=*1.39e-8 *** |
|  | Inoc (high) - Organic + Inorganic + Inoc (high) | -7.46 | *p=*3.52e-6 *** |
|  | Inoc (high) - Organic + Inorganic | -6.03 | *p=*6.35e-5 *** |
|  | Inoc (high) - Inorganic | -2.97 | *p=*0.03 * |
|  | Organic + Inorganic + Inoc (low) - Organic + Inorganic + Inoc (high) | 2.72 | *p=*0.044 * |
|  | Organic + Inorganic + Inoc (low) - Organic + Inorganic | 4.15 | *p=*0.003 ** |
|  | Organic + Inorganic + Inoc (low) - Organic | 10.58 | *p=*7.48e-9 *** |
|  | Organic + Inorganic + Inoc (low) - Inorganic | 7.22 | *p=*5.58e-6 *** |
|  | Organic + Inorganic + Inoc (high) - Organic | 7.85 | *p=*1.59e-6 *** |
|  | Organic + Inorganic + Inoc (high) - Inorganic | 4.49 | *p=*0.002 ** |
|  | Organic + Inorganic - Organic | 6.42 | *p=*3.23e-5 *** |
|  | Organic + Inorganic - Inorganic | 3.06 | *p=*0.026 * |
|  | Organic - Inorganic | -3.36 | *p=*0.016 * |
| Treatment * Site:  Contrasts between Musanze - Nyamagabe | Untreated | 4.72 | *p=*6.29e-4 *** |
|  | Inoc (low) | 11.96 | *p=*6.11e-11 *** |
|  | Inoc (high) | 12.06 | *p=*5.07e-11 *** |
|  | Organic + Inorganic + Inoc (low) | 19.16 | *p=*2.47e-16 *** |
|  | Organic + Inorganic + Inoc (high) | 20.47 | *p=*3.75e-17 *** |
|  | Organic + Inorganic | 20.32 | *p=*4.67e-17 *** |
|  | Organic | 12.19 | *p=*3.86e-11 *** |
|  | Inorganic | 13.42 | *p=*3.62e-12 *** |
| **Season 2025B ANOVA results** | | | |
| **Fixed effect** | **F statistic** | **Degrees of freedom** | ***P* value** |
| Treatment | 68.81 | 7,32 | *p=*1.77e-17 *** |
| Site | 611.41 | 1,32 | *p=*2.01e-22 *** |
| Treatment*Site | 9.52 | 7,32 | *p=*2.43e-6 *** |
| **Season 2025B post-hoc pairwise comparisons** | | | |
| **ANOVA fixed effect** | **Contrast** | **Estimate** | ***P* value** |
| Treatment * Site:  Contrasts within Musanze | Untreated - Inoc (low) | -9.70 | *p=*1.07e-10 *** |
|  | Untreated - Inoc (high) | -7.52 | *p=*2.56e-8 *** |
|  | Untreated - Organic + Inorganic + Inoc (low) | -13.65 | *p=*5.34e-14 *** |
|  | Untreated - Organic + Inorganic + Inoc (high) | -12.06 | *p=*7.86e-13 *** |
|  | Untreated - Organic + Inorganic | -10.73 | *p=*1.14e-11 *** |
|  | Untreated - Organic | -6.56 | *p=*3.14e-7 *** |
|  | Untreated - Inorganic | -9.19 | *p=*3.18e-10 *** |
|  | Inoc (low) - Inoc (high) | 2.18 | *p=*0.039 * |
|  | Inoc (low) - Organic + Inorganic + Inoc (low) | -3.95 | *p=*4.81e-4 *** |
|  | Inoc (low) - Organic + Inorganic + Inoc (high) | -2.36 | *p=*0.027 * |
|  | Inoc (low) - Organic | 3.14 | *p=*0.004 ** |
|  | Inoc (high) - Organic + Inorganic + Inoc (low) | -6.13 | *p=*1.01e-6 *** |
|  | Inoc (high) - Organic + Inorganic + Inoc (high) | -4.54 | *p=*1.03e-4 *** |
|  | Inoc (high) - Organic + Inorganic | -3.21 | *p=*0.004 ** |
|  | Organic + Inorganic + Inoc (low) - Organic + Inorganic | 2.92 | *p=*0.007 ** |
|  | Organic + Inorganic + Inoc (low) - Organic | 7.09 | *p=*7.57e-8 *** |
|  | Organic + Inorganic + Inoc (low) - Inorganic | 4.46 | *p=*1.22e-4 *** |
|  | Organic + Inorganic + Inoc (high) - Organic | 5.5 | *p=*6.09e-6 *** |
|  | Organic + Inorganic + Inoc (high) - Inorganic | 2.87 | *p=*0.008 ** |
|  | Organic + Inorganic - Organic | 4.17 | *p=*2.69e-4 *** |
|  | Organic - Inorganic | -2.63 | *p=*0.014 * |
| Treatment * Site:  Contrasts within Nyamagabe | Untreated - Organic + Inorganic + Inoc (low) | -11.54 | *p=*5.04e-12 *** |
|  | Untreated - Organic + Inorganic + Inoc (high) | -9.66 | *p=*9.62e-11 *** |
|  | Untreated - Organic + Inorganic | -6.02 | *p=*1.27e-6 *** |
|  | Untreated - Organic | -3.89 | *p=*0.0005 *** |
|  | Inoc (low) - Organic + Inorganic + Inoc (low) | -10.17 | *p=*4.73e-11 *** |
|  | Inoc (low) - Organic + Inorganic + Inoc (high) | -8.28 | *p=*3.06e-9 *** |
|  | Inoc (low) - Organic + Inorganic | -4.64 | *p=*6.43e-5 *** |
|  | Inoc (low) - Organic | -2.51 | *p=*0.02 * |
|  | Inoc (high) - Organic + Inorganic + Inoc (low) | -10.03 | *p=*4.73e-11 *** |
|  | Inoc (high) - Organic + Inorganic + Inoc (high) | -8.14 | *p=*3.38e-9 *** |
|  | Inoc (high) - Organic + Inorganic | -4.5 | *p=*8.53e-5 *** |
|  | Inoc (high) - Organic | -2.37 | *p=*0.026 * |
|  | Organic + Inorganic + Inoc (low) - Organic + Inorganic | 5.53 | *p=*4.68e-6 *** |
|  | Organic + Inorganic + Inoc (low) - Organic | 7.66 | *p=*1.16e-8 *** |
|  | Organic + Inorganic + Inoc (low) - Inorganic | 10.10 | *p=*4.73e-11 *** |
|  | Organic + Inorganic + Inoc (high) - Organic + Inorganic | 3.64 | *p=*9.74e-4 *** |
|  | Organic + Inorganic + Inoc (high) - Organic | 5.77 | *p=*2.44e-6 *** |
|  | Organic + Inorganic + Inoc (high) - Inorganic | 8.22 | *p=*3.12e-9 *** |
|  | Organic + Inorganic - Organic | 2.13 | *p=*0.044 * |
|  | Organic + Inorganic - Inorganic | 4.58 | *p=*7.24e-5 *** |
|  | Organic - Inorganic | 2.45 | *p=*0.023 * |
| Treatment * Site:  Contrasts between Musanze - Nyamagabe | Untreated | 4.1 | *p=*1.52e-4 *** |
|  | Inoc (low) | 12.43 | *p=*2.51e-14 *** |
|  | Inoc (high) | 10.1 | *p=*5.58e-12 *** |
|  | Organic + Inorganic + Inoc (low) | 6.21 | *p=*2.57e-7 *** |
|  | Organic + Inorganic + Inoc (high) | 6.50 | *p=*1.06e-7 *** |
|  | Organic + Inorganic | 8.81 | *p=*1.56e-10 *** |
|  | Organic | 6.77 | *p=*4.78e-8 *** |
|  | Inorganic | 11.85 | *p=*8.91e-14 *** |

***Supplementary Table 7.*** *Results of analyses of variance (ANOVAs) from linear models (~ Treatment * Site), and post-hoc pairwise comparisons run on potato plant biomass. Data were subset by Season prior to running models. P-values from post-hoc analyses are corrected for multiple comparisons; only marginally and statistically significant contrasts are shown. *p<0.05; **p<0.01; ***p<0.001; ·p<0.1; n.s. non-significant.*

| **Season 2025A ANOVA results** | | | |
| --- | --- | --- | --- |
| **Fixed effect** | **F statistic** | **Degrees of freedom** | ***P* value** |
| Treatment | 25.28 | 7,32 | *p=*2.48e-11 *** |
| Site | 43.18 | 1,32 | *p=*2.10e-7 *** |
| Treatment*Site | 1.41 | 7,32 | *p=*0.23 n.s. |
| **Post-hoc pairwise comparisons** | | | |
| **ANOVA fixed effect** | **Contrast** | **Estimate** | ***P* value** |
| Treatment | Untreated - Inoc (low) | -0.23 | *p=*0.002 *** |
|  | Untreated - Organic + Inorganic + Inoc (low) | -0.72 | *p=*3.11e-11 *** |
|  | Untreated - Organic + Inorganic + Inoc (high) | -0.44 | *p=*4.0e-7 *** |
|  | Untreated - Organic + Inorganic | -0.35 | *p=*1.47e-5 *** |
|  | Untreated - Organic | -0.15 | *p=*0.036 * |
|  | Untreated - Inorganic | -0.28 | *p=*2.92e-4 *** |
|  | Inoc (low) - Inoc (high) | 0.15 | *p=*0.036 * |
|  | Inoc (low) - Organic + Inorganic + Inoc (low) | -0.48 | *p=*7.70e-8 *** |
|  | Inoc (low) - Organic + Inorganic + Inoc (high) | -0.21 | *p=*0.005 ** |
|  | Inoc (high) - Organic + Inorganic + Inoc (low) | -0.63 | *p=*3.49e-10 *** |
|  | Inoc (high) - Organic + Inorganic + Inoc (high) | -0.36 | *p=*1.22e-5 *** |
|  | Inoc (high) - Organic + Inorganic | -0.27 | *p=*4.55e-4 *** |
|  | Inoc (high) - Inorganic | -0.2 | *p=*0.007 ** |
|  | Organic + Inorganic + Inoc (low) - Organic + Inorganic + Inoc (high) | 0.28 | *p=*2.92e-4 *** |
|  | Organic + Inorganic + Inoc (low) - Organic + Inorganic | 0.37 | *p=*8.16e-6 *** |
|  | Organic + Inorganic + Inoc (low) - Organic | 0.57 | *p=*3.07e-9 *** |
|  | Organic + Inorganic + Inoc (low) - Inorganic | 0.44 | *p=*4.0e-7 *** |
|  | Organic + Inorganic + Inoc (high) - Organic | 0.29 | *p=*1.91e-4 *** |
|  | Organic + Inorganic + Inoc (high) - Inorganic | 0.16 | *p=*0.026 * |
|  | Organic + Inorganic - Organic | 0.2 | *p=*0.006 ** |
| **Season 2025B ANOVA results** | | | |
| **Fixed effect** | **F statistic** | **Degrees of freedom** | ***P* value** |
| Treatment | 27.36 | 7,32 | *p=*8.63e-12 *** |
| Site | 405.98 | 1,32 | *p=*9.56e-20 *** |
| Treatment*Site | 2.10 | 7,32 | *p=*0.072 · |
| **Season 2025B post-hoc pairwise comparisons** | | | |
| **ANOVA fixed effect** | **Contrast** | **Estimate** | ***P* value** |
| Treatment | Untreated - Inoc (low) | -4.64 | *p=*6.46e-6 *** |
|  | Untreated - Inoc (high) | -4.03 | *p=*4.27e-5 *** |
|  | Untreated - Organic + Inorganic + Inoc (low) | -9.66 | *p=*4.76e-12 *** |
|  | Untreated - Organic + Inorganic + Inoc (high) | -7.92 | *p=*3.73e-10 *** |
|  | Untreated - Organic + Inorganic | -7.11 | *p=*3.27e-9 *** |
|  | Untreated - Organic | -4.25 | *p=*2.12e-5 *** |
|  | Untreated - Inorganic | -4.59 | *p=*6.93e-6 *** |
|  | Inoc (low) - Organic + Inorganic + Inoc (low) | -5.02 | *p=*1.83e-6 *** |
|  | Inoc (low) - Organic + Inorganic + Inoc (high) | -3.28 | *p=*4.69e-4 *** |
|  | Inoc (low) - Organic + Inorganic | -2.47 | *p=*0.006 ** |
|  | Inoc (high) - Organic + Inorganic + Inoc (low) | -5.63 | *p=*3.69e-7 *** |
|  | Inoc (high) - Organic + Inorganic + Inoc (high) | -3.89 | *p=*6.64e-5 *** |
|  | Inoc (high) - Organic + Inorganic | -3.07 | *p=*9.21e-4 *** |
|  | Organic + Inorganic + Inoc (low) - Organic + Inorganic + Inoc (high) | 1.74 | *p=*0.049 * |
|  | Organic + Inorganic + Inoc (low) - Organic + Inorganic | 2.56 | *p=*0.005 ** |
|  | Organic + Inorganic + Inoc (low) - Organic | 5.41 | *p=*6.40e-7 *** |
|  | Organic + Inorganic + Inoc (low) - Inorganic | 5.08 | *p=*1.77e-6 *** |
|  | Organic + Inorganic + Inoc (high) - Organic | 3.67 | *p=*1.35e-4 *** |
|  | Organic + Inorganic + Inoc (high) - Inorganic | 3.34 | *p=*4.18e-4 *** |
|  | Organic + Inorganic - Organic | 2.86 | *p=*0.002 ** |
|  | Organic + Inorganic - Inorganic | 2.52 | *p=*0.005 ** |

***Supplementary Table 8.*** *Results of analyses of variance (ANOVAs) from linear models (~ Treatment * Season), and post-hoc pairwise comparisons run on small-grade potato tuber yield. Data were subset by Site prior to running models. P-values from post-hoc analyses are corrected for multiple comparisons; only marginally and statistically significant contrasts are shown. *p<0.05; **p<0.01; ***p<0.001; ·p<0.1; n.s. non-significant.*

| **Musanze ANOVA results** | | | |
| --- | --- | --- | --- |
| **Fixed effect** | **F statistic** | **Degrees of freedom** | ***P* value** |
| Treatment | 47.50 | 7,32 | *p=*4.0e-15 *** |
| Season | 883.71 | 1,32 | *p=*7.04e-25 *** |
| Treatment * Season | 20.05 | 7,32 | *p=*4.91e-10 *** |
| **Musanze post-hoc pairwise comparisons** | | | |
| **ANOVA fixed effect** | **Contrast** | **Estimate** | ***P* value** |
| Treatment * Season:  Contrasts within 2025A | Untreated - Inoc (low) | -10.25 | *p=*7.49e-11 *** |
|  | Untreated - Inoc (high) | -7.94 | *p=*1.64e-8 *** |
|  | Untreated - Organic + Inorganic + Inoc (low) | -12.17 | *p=*1.40e-12 *** |
|  | Untreated - Organic + Inorganic + Inoc (high) | -18.95 | *p=*2.26e-17 *** |
|  | Untreated - Organic + Inorganic | -14.54 | *p=*2.24e-14 *** |
|  | Untreated - Organic | -8.94 | *p=*1.47e-9 *** |
|  | Untreated - Inorganic | -14.18 | *p=*3.0e-14 *** |
|  | Inoc (low) - Inoc (high) | 2.31 | *p=*0.035 * |
|  | Inoc (low) - Organic + Inorganic + Inoc (high) | -8.70 | *p=*2.43e-9 *** |
|  | Inoc (low) - Organic + Inorganic | -4.29 | *p=*2.73e-4 *** |
|  | Inoc (low) - Inorganic | -3.93 | *p=*6.70e-4 *** |
|  | Inoc (high) - Organic + Inorganic + Inoc (low) | -4.23 | *p=*3.03e-4 *** |
|  | Inoc (high) - Organic + Inorganic + Inoc (high) | -11.01 | *p=*1.50e-11 *** |
|  | Inoc (high) - Organic + Inorganic | -6.60 | *p=*5.46e-7 *** |
|  | Inoc (high) - Inorganic | -6.24 | *p=*1.40e-6 *** |
|  | Organic + Inorganic + Inoc (low) - Organic + Inorganic + Inoc (high) | -6.78 | *p=*3.65e-7 *** |
|  | Organic + Inorganic + Inoc (low) - Organic + Inorganic | -2.37 | *p=*0.032 * |
|  | Organic + Inorganic + Inoc (low) - Organic | 3.233 | *p=*0.004 ** |
|  | Organic + Inorganic + Inoc (high) - Organic + Inorganic | 4.42 | *p=*2.08e-4 *** |
|  | Organic + Inorganic + Inoc (high) - Organic | 10.01 | *p=*1.15e-10 *** |
|  | Organic + Inorganic + Inoc (high) - Inorganic | 4.77 | *p=*7.92e-5 *** |
|  | Organic + Inorganic - Organic | 5.60 | *p=*8.16e-6 *** |
|  | Organic - Inorganic | -5.24 | *p=*2.15e-5 *** |
| Treatment * Season:  Contrasts within 2025B | Untreated - Organic + Inorganic + Inoc (low) | -2.83 | *p=*0.04 * |
|  | Untreated - Organic + Inorganic + Inoc (high) | -4.96 | *p=*6.58e-4 *** |
|  | Inoc (low) - Organic + Inorganic + Inoc (low) | -2.63 | *p=*0.049 * |
|  | Inoc (low) - Organic + Inorganic + Inoc (high) | -4.75 | *p=*6.58-4 *** |
|  | Inoc (high) - Organic + Inorganic + Inoc (high) | -2.88 | *p=*0.04 * |
|  | Organic + Inorganic + Inoc (high) - Organic + Inorganic | 2.77 | *p=*0.04 * |
|  | Organic + Inorganic + Inoc (high) - Organic | 4.57 | *p=*7.33-4 *** |
|  | Organic + Inorganic + Inoc (high) - Inorganic | 3.67 | *p=*0.007 ** |
| **Nyamagabe ANOVA results** | | | |
| **Fixed effect** | **F statistic** | **Degrees of freedom** | ***P* value** |
| Treatment | 18.22 | 7,32 | *p=*1.63e-9 *** |
| Season | 33.94 | 1,32 | *p=*1.80e-6 *** |
| Treatment*Season | 1.71 | 7,32 | *p=*0.14 n.s. |
| **Nyamagabe post-hoc pairwise comparisons** | | | |
| **ANOVA fixed effect** | **Contrast** | **Estimate** | ***P* value** |
| Treatment | Untreated - Inoc (low) | -1.55 | *p=*0.019 * |
|  | Untreated - Inoc (high) | -1.74 | *p=*0.01 * |
|  | Untreated - Organic + Inorganic + Inoc (low) | -5.05 | *p=*1.77e-8 *** |
|  | Untreated - Organic + Inorganic + Inoc (high) | -4.75 | *p=*3.47e-8 *** |
|  | Untreated - Organic + Inorganic | -2.57 | *p=*2.94e-4 *** |
|  | Untreated - Inorganic | -2.31 | *p=*7.87e-4 *** |
|  | Inoc (low) - Organic + Inorganic + Inoc (low) | -3.50 | *p=*5.82e-6 *** |
|  | Inoc (low) - Organic + Inorganic + Inoc (high) | -3.20 | *p=*1.86e-5 *** |
|  | Inoc (high) - Organic + Inorganic + Inoc (low) | -3.32 | *p=*1.20e-5 *** |
|  | Inoc (high) - Organic + Inorganic + Inoc (high) | -3.02 | *p=*4.04e-5 *** |
|  | Organic + Inorganic + Inoc (low) - Organic + Inorganic | 2.48 | *p=*4.23e-4 *** |
|  | Organic + Inorganic + Inoc (low) - Organic | 4.08 | *p=*5.42e-7 *** |
|  | Organic + Inorganic + Inoc (low) - Inorganic | 2.73 | *p=*1.47e-4 *** |
|  | Organic + Inorganic + Inoc (high) - Organic + Inorganic | 2.18 | *p=*0.001 ** |
|  | Organic + Inorganic + Inoc (high) - Organic | 3.78 | *p=*1.76e-6 *** |
|  | Organic + Inorganic + Inoc (high) - Inorganic | 2.43 | *p=*4.84e-4 *** |
|  | Organic + Inorganic - Organic | 1.61 | *p=*0.017 * |
|  | Organic - Inorganic | -1.35 | *p=*0.042 * |

***Supplementary Table 9.*** *Results of analyses of variance (ANOVAs) from linear models (~ Treatment * Season) and post-hoc pairwise comparisons run on large-grade potato tuber yield. Data were subset by Site prior to running models. P-values from post-hoc analyses are corrected for multiple comparisons; only marginally and statistically significant contrasts are shown. *p<0.05; **p<0.01; ***p<0.001; ·p<0.1; n.s. non-significant.*

| **Musanze ANOVA results** | | | |
| --- | --- | --- | --- |
| **Fixed effect** | **F statistic** | **Degrees of freedom** | ***P* value** |
| Treatment | 92.55 | 7,32 | *p=*2.07e-19 *** |
| Season | 8.66e-4 | 1,32 | *p=*0.98 n.s. |
| Treatment*Season | 25.86 | 7,32 | *p=*1.83e-11 *** |
| **Musanze post-hoc pairwise comparisons** | | | |
| **ANOVA fixed effect** | **Contrast** | **Estimate** | ***P* value** |
| Treatment * Season:  Contrasts within 2025A | Untreated - Inoc (low) | -3.16 | *p=*0.002 ** |
|  | Untreated - Inoc (high) | -4.47 | *p=*2.46e-5 *** |
|  | Untreated - Organic + Inorganic + Inoc (low) | -17.53 | *p=*3.88e-18 *** |
|  | Untreated - Organic + Inorganic + Inoc (high) | -9.34 | *p=*1.86e-11 *** |
|  | Untreated - Organic + Inorganic | -12.16 | *p=*2.33e-14 *** |
|  | Untreated - Organic | -3.22 | *p=*0.001 ** |
|  | Untreated - Inorganic | -2.56 | *p=*0.008 ** |
|  | Inoc (low) - Organic + Inorganic + Inoc (low) | -14.37 | *p=*3.60e-16 *** |
|  | Inoc (low) - Organic + Inorganic + Inoc (high) | -6.18 | *p=*1.19e-7 *** |
|  | Inoc (low) - Organic + Inorganic | -9.0 | *p=*4.13e-11 *** |
|  | Inoc (high) - Organic + Inorganic + Inoc (low) | -13.06 | *p=*3.90e-15 *** |
|  | Inoc (high) - Organic + Inorganic + Inoc (high) | -4.87 | *p=*7.11e-6 *** |
|  | Inoc (high) - Organic + Inorganic | -7.69 | *p=*1.32e-9 *** |
|  | Inoc (high) - Inorganic | 1.91 | *p=*0.046 * |
|  | Organic + Inorganic + Inoc (low) - Organic + Inorganic + Inoc (high) | 8.19 | *p=*3.33e-10 *** |
|  | Organic + Inorganic + Inoc (low) - Organic + Inorganic | 5.37 | *p=*1.46e-6 *** |
|  | Organic + Inorganic + Inoc (low) - Organic | 14.31 | *p=*3.60e-16 *** |
|  | Organic + Inorganic + Inoc (low) - Inorganic | 14.97 | *p=*1.98e-16 *** |
|  | Organic + Inorganic + Inoc (high) - Organic + Inorganic | -2.82 | *p=*0.004 ** |
|  | Organic + Inorganic + Inoc (high) - Organic | 6.12 | *p=*1.33e-7 *** |
|  | Organic + Inorganic + Inoc (high) - Inorganic | 6.78 | *p=*1.93e-8 *** |
|  | Organic + Inorganic - Organic | 8.95 | *p=*4.34e-11 *** |
|  | Organic + Inorganic - Inorganic | 9.60 | *p=*1.05e-11 *** |
| Treatment * Season:  Contrasts within 2025B | Untreated - Inoc (low) | -9.50 | *p=*4.89e-11 *** |
|  | Untreated - Inoc (high) | -5.45 | *p=*2.56e-6 *** |
|  | Untreated - Organic + Inorganic + Inoc (low) | -10.81 | *p=*3.37e-12 *** |
|  | Untreated - Organic + Inorganic + Inoc (high) | -7.1 | *p=*1.87e-8 *** |
|  | Untreated - Organic + Inorganic | -8.54 | *p=*4.54e-10 *** |
|  | Untreated - Organic | -6.17 | *p=*2.87e-7 *** |
|  | Untreated - Inorganic | -7.9 | *p=*2.13e-9 *** |
|  | Inoc (low) - Inoc (high) | 4.05 | *p=*1.79e-4 *** |
|  | Inoc (low) - Organic + Inorganic + Inoc (high) | 2.40 | *p=*0.018 * |
|  | Inoc (low) - Organic | 3.33 | *p=*0.002 ** |
|  | Inoc (high) - Organic + Inorganic + Inoc (low) | -5.37 | *p=*2.91e-6 *** |
|  | Inoc (high) - Organic + Inorganic | -3.09 | *p=*0.003 ** |
|  | Inoc (high) - Inorganic | -2.45 | *p=*0.017 * |
|  | Organic + Inorganic + Inoc (low) - Organic + Inorganic + Inoc (high) | 3.71 | *p=*4.85e-4 *** |
|  | Organic + Inorganic + Inoc (low) - Organic + Inorganic | 2.28 | *p=*0.023 * |
|  | Organic + Inorganic + Inoc (low) - Organic | 4.64 | *p=*2.81e-5 *** |
|  | Organic + Inorganic + Inoc (low) - Inorganic | 2.91 | *p=*0.005 ** |
|  | Organic + Inorganic - Organic | 2.37 | *p=*0.019 * |
| **Nyamagabe ANOVA results** | | | |
| **Fixed effect** | **F statistic** | **Degrees of freedom** | ***P* value** |
| Treatment | 27.54 | 7,32 | *p=*7.94e-12 *** |
| Season | 61.98 | 1,32 | *p=*5.55e-9 *** |
| Treatment*Season | 2.91 | 7,32 | *p=*0.018 * |
| **Nyamagabe post-hoc pairwise comparisons** | | | |
| **ANOVA fixed effect** | **Contrast** | **Estimate** | ***P* value** |
| Treatment * Season:  Contrasts within 2025A | Untreated - Inoc (low) | -4.76 | *p=*3.76e-4 *** |
|  | Untreated - Inoc (high) | -3.16 | *p=*0.011 * |
|  | Untreated - Organic + Inorganic + Inoc (low) | -10.49 | *p=*1.25e-9 *** |
|  | Untreated - Organic + Inorganic + Inoc (high) | -7.10 | *p=*1.41e-6 *** |
|  | Untreated - Organic + Inorganic | -7.58 | *p=*7.98e-7 *** |
|  | Untreated - Organic | -3.69 | *p=*0.004 ** |
|  | Untreated - Inorganic | -4.57 | *p=*5.62e-4 *** |
|  | Inoc (low) - Organic + Inorganic + Inoc (low) | -5.73 | *p=*3.23e-5 *** |
|  | Inoc (low) - Organic + Inorganic + Inoc (high) | -2.34 | *p=*0.050 * |
|  | Inoc (low) - Organic + Inorganic | -2.82 | *p=*0.020 * |
|  | Inoc (high) - Organic + Inorganic + Inoc (low) | -7.33 | *p=*1.02e-6 *** |
|  | Inoc (high) - Organic + Inorganic + Inoc (high) | -3.95 | *p=*0.002 * |
|  | Inoc (high) - Organic + Inorganic | -4.43 | *p=*7.31e-4 *** |
|  | Organic + Inorganic + Inoc (low) - Organic + Inorganic + Inoc (high) | 3.39 | *p=*0.007 ** |
|  | Organic + Inorganic + Inoc (low) - Organic + Inorganic | 2.91 | *p=*0.017 * |
|  | Organic + Inorganic + Inoc (low) - Organic | 6.80 | *p=*2.50e-6 *** |
|  | Organic + Inorganic + Inoc (low) - Inorganic | 5.92 | *p=*2.22e-5 *** |
|  | Organic + Inorganic + Inoc (high) - Organic | 3.42 | *p=*0.007 ** |
|  | Organic + Inorganic + Inoc (high) - Inorganic | 2.54 | *p=*0.035 * |
|  | Organic + Inorganic - Organic | 3.90 | *p=*0.002 ** |
|  | Organic + Inorganic - Inorganic | 3.02 | *p=*0.014 * |
| Treatment * Season:  Contrasts within 2025B | Untreated - Organic + Inorganic + Inoc (low) | -6.21 | *p=*2.05e-5 *** |
|  | Untreated - Organic + Inorganic + Inoc (high) | -5.59 | *p=*4.67e-5 *** |
|  | Untreated - Organic + Inorganic | -4.41 | *p=*7.00e-4 *** |
|  | Untreated - Organic | -2.95 | *p=*0.018 * |
|  | Inoc (low) - Organic + Inorganic + Inoc (low) | -6.53 | *p=*2.05e-5 *** |
|  | Inoc (low) - Organic + Inorganic + Inoc (high) | -5.91 | *p=*2.76e-5 *** |
|  | Inoc (low) - Organic + Inorganic | -4.73 | *p=*3.68e-4 *** |
|  | Inoc (low) - Organic | -3.27 | *p=*0.010 * |
|  | Inoc (high) - Organic + Inorganic + Inoc (low) | -6.24 | *p=*2.05e-5 *** |
|  | Inoc (high) - Organic + Inorganic + Inoc (high) | -5.62 | *p=*4.67e-5 *** |
|  | Inoc (high) - Organic + Inorganic | -4.44 | *p=*7.00e-4 *** |
|  | Inoc (high) - Organic | -2.98 | *p=*0.017 * |
|  | Organic + Inorganic + Inoc (low) - Organic | 3.26 | *p=*0.010 * |
|  | Organic + Inorganic + Inoc (low) - Inorganic | 5.93 | *p=*2.76e-5 *** |
|  | Organic + Inorganic + Inoc (high) - Organic | 2.64 | *p=*0.031 * |
|  | Organic + Inorganic + Inoc (high) - Inorganic | 5.31 | *p=*8.77e-5 *** |
|  | Organic + Inorganic - Inorganic | 4.12 | *p=*0.001 ** |
|  | Organic - Inorganic | 2.67 | *p=*0.031 * |

***Supplementary Table 10.*** *Results of analyses of variance (ANOVAs) from linear models (~ Treatment * Season), and post-hoc pairwise comparisons run on total potato tuber yield. Data were subset by Site prior to running models. P-values from post-hoc analyses are corrected for multiple comparisons; only marginally and statistically significant contrasts are shown. *p<0.05; **p<0.01; ***p<0.001; ·p<0.1; n.s. non-significant.*

| **Musanze ANOVA results** | | | |
| --- | --- | --- | --- |
| **Fixed effect** | **F statistic** | **Degrees of freedom** | ***P* value** |
| Treatment | 184.65 | 7,32 | *p=*5.07e-24 *** |
| Season | 828.38 | 1,32 | *p=*1.91e-24 *** |
| Treatment*Season | 37.90 | 7,32 | *p=*9.92e-14 *** |
| **Musanze post-hoc pairwise comparisons** | | | |
| **ANOVA fixed effect** | **Contrast** | **Estimate** | ***P* value** |
| Treatment * Season:  Contrasts within 2025A | Untreated - Inoc (low) | -13.41 | *p=*7.96e-14 *** |
|  | Untreated - Inoc (high) | -12.41 | *p=*5.05e-13 *** |
|  | Untreated - Organic + Inorganic + Inoc (low) | -29.7 | *p=*7.90e-23 *** |
|  | Untreated - Organic + Inorganic + Inoc (high) | -28.29 | *p=*1.77e-22 *** |
|  | Untreated - Organic + Inorganic | -26.7 | *p=*6.91e-22 *** |
|  | Untreated - Organic | -12.15 | *p=*8.21e-13 *** |
|  | Untreated - Inorganic | -16.74 | *p=*3.62e-16 *** |
|  | Inoc (low) - Organic + Inorganic + Inoc (low) | -16.29 | *p=*6.67e-16 *** |
|  | Inoc (low) - Organic + Inorganic + Inoc (high) | -14.88 | *p=*5.97e-15 *** |
|  | Inoc (low) - Organic + Inorganic | -13.29 | *p=*9.29e-14 *** |
|  | Inoc (low) - Inorganic | -3.33 | *p=*0.004 ** |
|  | Inoc (high) - Organic + Inorganic + Inoc (low) | -17.29 | *p=*1.71e-16 *** |
|  | Inoc (high) - Organic + Inorganic + Inoc (high) | -15.88 | *p=*1.08e-15 *** |
|  | Inoc (high) - Organic + Inorganic | -14.29 | *p=*1.52e-14 *** |
|  | Inoc (high) - Inorganic | -4.33 | *p=*3.14e-4 *** |
|  | Organic + Inorganic + Inoc (low) - Organic + Inorganic | 3 | *p=*0.009 ** |
|  | Organic + Inorganic + Inoc (low) - Organic | 17.55 | *p=*1.40e-16 *** |
|  | Organic + Inorganic + Inoc (low) - Inorganic | 12.96 | *p=*1.70e-13 *** |
|  | Organic + Inorganic + Inoc (high) - Organic | 16.13 | *p=*7.72e-16 *** |
|  | Organic + Inorganic + Inoc (high) - Inorganic | 11.55 | *p=*2.89e-12 *** |
|  | Organic + Inorganic - Organic | 14.55 | *p=*1.02e-14 *** |
|  | Organic + Inorganic - Inorganic | 9.96 | *p=*1.06e-10 *** |
|  | Organic - Inorganic | -4.58 | *p=*1.64e-4 *** |
| Treatment * Season:  Contrasts within 2025B | Untreated - Inoc (low) | -9.70 | *p=*9.43e-10 *** |
|  | Untreated - Inoc (high) | -7.52 | *p=*1.67e-7 *** |
|  | Untreated - Organic + Inorganic + Inoc (low) | -13.65 | *p=*6.37e-13 *** |
|  | Untreated - Organic + Inorganic + Inoc (high) | -12.06 | *p=*8.53e-12 *** |
|  | Untreated - Organic + Inorganic | -10.73 | *p=*1.11e-10 *** |
|  | Untreated - Organic | -6.56 | *p=*1.71e-6 *** |
|  | Untreated - Inorganic | -9.19 | *p=*2.65e-9 *** |
|  | Inoc (low) - Organic + Inorganic + Inoc (low) | -3.95 | *p=*0.001 ** |
|  | Inoc (low) - Organic + Inorganic + Inoc (high) | -2.36 | *p=*0.044 * |
|  | Inoc (low) - Organic | 3.14 | *p=*0.009 ** |
|  | Inoc (high) - Organic + Inorganic + Inoc (low) | -6.13 | *p=*4.98e-6 *** |
|  | Inoc (high) - Organic + Inorganic + Inoc (high) | -4.54 | *p=*3.36e-4 *** |
|  | Inoc (high) - Organic + Inorganic | -3.21 | *p=*0.008 ** |
|  | Organic + Inorganic + Inoc (low) - Organic + Inorganic | 2.92 | *p=*0.014 * |
|  | Organic + Inorganic + Inoc (low) - Organic | 7.09 | *p=*4.58e-7 *** |
|  | Organic + Inorganic + Inoc (low) - Inorganic | 4.46 | *p=*3.86e-4 *** |
|  | Organic + Inorganic + Inoc (high) - Organic | 5.5 | *p=*2.59e-5 *** |
|  | Organic + Inorganic + Inoc (high) - Inorganic | 2.87 | *p=*0.015 * |
|  | Organic + Inorganic - Organic | 4.17 | *p=*7.82e-4 *** |
|  | Organic - Inorganic | -2.63 | *p=*0.025 * |
| **Nyamagabe ANOVA results** | | | |
| **Fixed effect** | **F statistic** | **Degrees of freedom** | ***P* value** |
| Treatment | 58.26 | 7,32 | *p=*2.06e-16 *** |
| Season | 128.35 | 1,32 | *p=*9.83e-13 *** |
| Treatment*Season | 3.52 | 7,32 | *p=*0.007 ** |
| **Nyamagabe post-hoc pairwise comparisons** | | | |
| **ANOVA fixed effect** | **Contrast** | **Estimate** | ***P* value** |
| Treatment * Season:  Contrasts within 2025A | Untreated - Inoc (low) | -6.17 | *p=*1.99e-5 *** |
|  | Untreated - Inoc (high) | -5.08 | *p=*2.54e-4 *** |
|  | Untreated - Organic + Inorganic + Inoc (low) | -15.26 | *p=*6.88e-13 *** |
|  | Untreated - Organic + Inorganic + Inoc (high) | -12.54 | *p=*5.91e-11 *** |
|  | Untreated - Organic + Inorganic | -11.11 | *p=*7.69e-10 *** |
|  | Untreated - Organic | -4.68 | *p=*5.81e-4 *** |
|  | Untreated - Inorganic | -8.04 | *p=*3.64e-7 *** |
|  | Inoc (low) - Organic + Inorganic + Inoc (low) | -9.09 | *p=*3.51e-8 *** |
|  | Inoc (low) - Organic + Inorganic + Inoc (high) | -6.37 | *p=*1.30e-5 *** |
|  | Inoc (low) - Organic + Inorganic | -4.94 | *p=*3.32e-4 *** |
|  | Inoc (high) - Organic + Inorganic + Inoc (low) | -10.18 | *p=*3.49e-9 *** |
|  | Inoc (high) - Organic + Inorganic + Inoc (high) | -7.46 | *p=*1.17e-6 *** |
|  | Inoc (high) - Organic + Inorganic | -6.03 | *p=*2.59e-5 *** |
|  | Inoc (high) - Inorganic | -2.97 | *p=*0.021* |
|  | Organic + Inorganic + Inoc (low) - Organic + Inorganic + Inoc (high) | 2.72 | *p=*0.032 * |
|  | Organic + Inorganic + Inoc (low) - Organic + Inorganic | 4.15 | *p=*0.002 ** |
|  | Organic + Inorganic + Inoc (low) - Organic | 10.58 | *p=*1.82e-9 *** |
|  | Organic + Inorganic + Inoc (low) - Inorganic | 7.217 | *p=*1.92e-6 *** |
|  | Organic + Inorganic + Inoc (high) - Organic | 7.85 | *p=*5.05e-7 *** |
|  | Organic + Inorganic + Inoc (high) - Inorganic | 4.49 | *p=*8.64e-4 *** |
|  | Organic + Inorganic - Organic | 6.42 | *p=*1.24e-5 *** |
|  | Organic + Inorganic - Inorganic | 3.06 | *p=*0.018 * |
|  | Organic - Inorganic | -3.36 | *p=*0.010 * |
| Treatment * Season:  Contrasts within 2025B | Untreated - Organic + Inorganic + Inoc (low) | -11.54 | *p=*9.12e-10 *** |
|  | Untreated - Organic + Inorganic + Inoc (high) | -9.66 | *p=*1.15e-8 *** |
|  | Untreated - Organic + Inorganic | -6.02 | *p=*3.75e-5 *** |
|  | Untreated - Organic | -3.89 | *p=*0.004 ** |
|  | Inoc (low) - Organic + Inorganic + Inoc (low) | -10.17 | *p=*6.19e-9 *** |
|  | Inoc (low) - Organic + Inorganic + Inoc (high) | -8.28 | *p=*2.40e-7 *** |
|  | Inoc (low) - Organic + Inorganic | -4.64 | *p=*8.43e-4 *** |
|  | Inoc (high) - Organic + Inorganic + Inoc (low) | -10.03 | *p=*6.19e-9 *** |
|  | Inoc (high) - Organic + Inorganic + Inoc (high) | -8.14 | *p=*2.52e-7 *** |
|  | Inoc (high) - Organic + Inorganic | -4.5 | *p=*0.001 ** |
|  | Organic + Inorganic + Inoc (low) - Organic + Inorganic | 5.53 | *p=*1.05e-4 *** |
|  | Organic + Inorganic + Inoc (low) - Organic | 7.66 | *p=*7.24e-7 *** |
|  | Organic + Inorganic + Inoc (low) - Inorganic | 10.10 | *p=*6.19e-9 *** |
|  | Organic + Inorganic + Inoc (high) - Organic + Inorganic | 3.64 | *p=*0.007 ** |
|  | Organic + Inorganic + Inoc (high) - Organic | 5.77 | *p=*6.29e-5 *** |
|  | Organic + Inorganic + Inoc (high) - Inorganic | 8.22 | *p=*2.40e-7 *** |
|  | Organic + Inorganic - Inorganic | 4.58 | *p=*9.11e-4 *** |

***Supplementary Table 11.*** *Results of analyses of variance (ANOVAs) from linear models (~ Treatment * Season), and post-hoc pairwise comparisons run on potato plant biomass. Data were subset by Site prior to running models. P-values from post-hoc analyses are corrected for multiple comparisons; only marginally and statistically significant contrasts are shown. *p<0.05; **p<0.01; ***p<0.001; ·p<0.1; n.s. non-significant.*

| **Musanze ANOVA results** | | | |
| --- | --- | --- | --- |
| **Fixed effect** | **F statistic** | **Degrees of freedom** | ***P* value** |
| Treatment | 15.53 | 7,32 | *p=*1.12e-8 *** |
| Season | 1402.13 | 1,32 | *p=*5.34e-8 *** |
| Treatment*Season | 12.43 | 7,32 | *p=*1.44e-7 *** |
| **Musanze post-hoc pairwise comparisons** | | | |
| **ANOVA fixed effect** | **Contrast** | **Estimate** | ***P* value** |
| Treatment * Season:  Contrasts within 2025B | Untreated - Inoc (low) | -6.61 | *p=*7.53e-7 *** |
|  | Untreated - Inoc (high) | -5.79 | *p=*5.23e-6 *** |
|  | Untreated - Organic + Inorganic + Inoc (low) | -12.01 | *p=*3.79e-12 *** |
|  | Untreated - Organic + Inorganic + Inoc (high) | -9.93 | *p=*2.06e-10 *** |
|  | Untreated - Organic + Inorganic | -9.84 | *p=*2.06e-10 *** |
|  | Untreated - Organic | -5.70 | *p=*5.94e-6 *** |
|  | Untreated - Inorganic | -6.66 | *p=*7.53e-7 *** |
|  | Inoc (low) - Organic + Inorganic + Inoc (low) | -5.40 | *p=*1.33e-5 *** |
|  | Inoc (low) - Organic + Inorganic + Inoc (high) | -3.32 | *p=*0.003 ** |
|  | Inoc (low) - Organic + Inorganic | -3.23 | *p=*0.004 ** |
|  | Inoc (high) - Organic + Inorganic + Inoc (low) | -6.22 | *p=*1.67e-6 *** |
|  | Inoc (high) - Organic + Inorganic + Inoc (high) | -4.14 | *p=*3.90e-4 *** |
|  | Inoc (high) - Organic + Inorganic | -4.05 | *p=*4.63e-4 *** |
|  | Organic + Inorganic + Inoc (low) - Organic + Inorganic | 2.17 | *p=*0.048 * |
|  | Organic + Inorganic + Inoc (low) - Organic | 6.30 | *p=*1.53e-6 *** |
|  | Organic + Inorganic + Inoc (low) - Inorganic | 5.34 | *p=*1.41e-5 *** |
|  | Organic + Inorganic + Inoc (high) - Organic | 4.22 | *p=*3.51e-4 *** |
|  | Organic + Inorganic + Inoc (high) - Inorganic | 3.26 | *p=*0.004 ** |
|  | Organic + Inorganic - Organic | 4.13 | *p=*3.90e-4 *** |
|  | Organic + Inorganic - Inorganic | 3.17 | *p=*0.004 ** |
| **Nyamagabe ANOVA results** | | | |
| **Fixed effect** | **F statistic** | **Degrees of freedom** | ***P* value** |
| Treatment | 20.02 | 7,32 | *p=*5.02e-10 *** |
| Site | 683.27 | 1,32 | *p=*3.68e-23 *** |
| Treatment*Site | 14.0 | 7,32 | *p=*3.76e-8 *** |
| **Nyamagabe post-hoc pairwise comparisons** | | | |
| **ANOVA fixed effect** | **Contrast** | **Estimate** | ***P* value** |
| Treatment * Season:  Contrasts within 2025B | Untreated - Inoc (low) | -2.67 | *p=*8.07e-5 *** |
|  | Untreated - Inoc (high) | -2.28 | *p=*5.09e-4 *** |
|  | Untreated - Organic + Inorganic + Inoc (low) | -7.32 | *p=*6.55e-13 *** |
|  | Untreated - Organic + Inorganic + Inoc (high) | -5.92 | *p=*8.39e-11 *** |
|  | Untreated - Organic + Inorganic | -4.38 | *p=*2.66e-8 *** |
|  | Untreated - Organic | -2.8 | *p=*4.36e-5 *** |
|  | Untreated - Inorganic | -2.51 | *p=*1.69e-4 *** |
|  | Inoc (low) - Organic + Inorganic + Inoc (low) | -4.65 | *p=*1.00e-8 *** |
|  | Inoc (low) - Organic + Inorganic + Inoc (high) | -3.25 | *p=*5.49e-6 *** |
|  | Inoc (low) - Organic + Inorganic | -1.71 | *p=*0.007 ** |
|  | Inoc (high) - Organic + Inorganic + Inoc (low) | -5.04 | *p=*2.73e-9 *** |
|  | Inoc (high) - Organic + Inorganic + Inoc (high) | -3.64 | *p=*9.38e-7 *** |
|  | Inoc (high) - Organic + Inorganic | -2.10 | *p=*0.001 ** |
|  | Organic + Inorganic + Inoc (low) - Organic + Inorganic + Inoc (high) | 1.4 | *p=*0.023 * |
|  | Organic + Inorganic + Inoc (low) - Organic + Inorganic | 2.94 | *p=*2.29e-5 *** |
|  | Organic + Inorganic + Inoc (low) - Organic | 4.52 | *p=*1.58e-8 *** |
|  | Organic + Inorganic + Inoc (low) - Inorganic | 4.81 | *p=*5.97e-9 *** |
|  | Organic + Inorganic + Inoc (high) - Organic + Inorganic | 1.54 | *p=*0.013 * |
|  | Organic + Inorganic + Inoc (high) - Organic | 3.12 | *p=*9.96e-6 *** |
|  | Organic + Inorganic + Inoc (high) - Inorganic | 3.41 | *p=*2.72e-6 *** |
|  | Organic + Inorganic - Organic | 1.58 | *p=*0.012 * |
|  | Organic + Inorganic - Inorganic | 1.87 | *p=*0.003 ** |

***Supplementary Table 12.*** *Results of analyses of variance (ANOVAs) from linear models (~ Treatment * Site), and post-hoc pairwise comparisons run on maize grain yield. Data were subset by Season prior to running models. P-values from post-hoc analyses are corrected for multiple comparisons; only marginally and statistically significant contrasts are shown. *p<0.05; **p<0.01; ***p<0.001; ·p<0.1; n.s. non-significant.*

| **Season 2025A ANOVA results** | | | |
| --- | --- | --- | --- |
| **Fixed effect** | **F statistic** | **Degrees of freedom** | ***P* value** |
| Treatment | 198.57 | 7,32 | *p=*1.63e-24 *** |
| Site | 0.56 | 1,32 | *p=*0.46 n.s. |
| Treatment*Site | 3.86 | 7,32 | *p=*0.004 ** |
| **Season 2025 post-hoc pairwise comparisons** | | | |
| **ANOVA fixed effect** | **Contrast** | **Estimate** | ***P* value** |
| Treatment * Site:  Contrasts within Kayonza | Untreated - Inoc (low) | -1.72 | *p=*8.99e-5 *** |
|  | Untreated - Inoc (high) | -2.43 | *p=*5.25e-7 *** |
|  | Untreated - Organic + Inorganic + Inoc (low) | -6.84 | *p=*1.39e-17 *** |
|  | Untreated - Organic + Inorganic + Inoc (high) | -8.58 | *p=*6.72e-20 *** |
|  | Untreated - Organic + Inorganic | -4.76 | *p=*1.66e-13 *** |
|  | Untreated - Organic | -1.56 | *p=*2.77e-4 *** |
|  | Untreated - Inorganic | -3.29 | *p=*1.15e-9 *** |
|  | Inoc (low) - Organic + Inorganic + Inoc (low) | -5.12 | *p=*2.64e-14 *** |
|  | Inoc (low) - Organic + Inorganic + Inoc (high) | -6.85 | *p=*1.39e-17 *** |
|  | Inoc (low) - Organic + Inorganic | -3.04 | *p=*5.97e-9 *** |
|  | Inoc (low) - Inorganic | -1.57 | *p=*2.75e-4 *** |
|  | Inoc (high) - Organic + Inorganic + Inoc (low) | -4.42 | *p=*1.11e-12 *** |
|  | Inoc (high) - Organic + Inorganic + Inoc (high) | -6.15 | *p=*2.46e-16 *** |
|  | Inoc (high) - Organic + Inorganic | -2.34 | *p=*9.82e-7 *** |
|  | Inoc (high) - Organic | 0.86 | *p=*0.031 * |
|  | Inoc (high) - Inorganic | -0.87 | *p=*0.031 * |
|  | Organic + Inorganic + Inoc (low) - Organic + Inorganic + Inoc (high) | -1.73 | *p=*8.97e-5 *** |
|  | Organic + Inorganic + Inoc (low) - Organic + Inorganic | 2.08 | *p=*6.69e-6 *** |
|  | Organic + Inorganic + Inoc (low) - Organic | 5.28 | *p=*1.29e-14 *** |
|  | Organic + Inorganic + Inoc (low) - Inorganic | 3.55 | *p=*2.17e-10 *** |
|  | Organic + Inorganic + Inoc (high) - Organic + Inorganic | 3.81 | *p=*4.20e-11 *** |
|  | Organic + Inorganic + Inoc (high) - Organic | 7.01 | *p=*1.35e-17 *** |
|  | Organic + Inorganic + Inoc (high) - Inorganic | 5.28 | *p=*1.29e-14 *** |
|  | Organic + Inorganic - Organic | 3.2 | *p=*2.06e-9 *** |
|  | Organic + Inorganic - Inorganic | 1.47 | *p=*5.33e-4 *** |
|  | Organic - Inorganic | -1.73 | *p=*8.97e-5 *** |
| Treatment * Site:  Contrasts within Ngoma | Untreated - Inoc (low) | -2.69 | *p=*8.25e-8 *** |
|  | Untreated - Inoc (high) | -3.41 | *p=*7.47e-10 *** |
|  | Untreated - Organic + Inorganic + Inoc (low) | -6.05 | *p=*9.98e-16 *** |
|  | Untreated - Organic + Inorganic + Inoc (high) | -8.16 | *p=*3.05e-19 *** |
|  | Untreated - Organic + Inorganic | -4.84 | *p=*1.91e-13 *** |
|  | Untreated - Organic | -2.89 | *p=*2.04e-8 *** |
|  | Untreated - Inorganic | -3.53 | *p=*3.65e-10 *** |
|  | Inoc (low) - Inoc (high) | -0.72 | *p=*0.076 · |
|  | Inoc (low) - Organic + Inorganic + Inoc (low) | -3.36 | *p=*9.69e-10 *** |
|  | Inoc (low) - Organic + Inorganic + Inoc (high) | -5.47 | *p=*1.15e-14 *** |
|  | Inoc (low) - Organic + Inorganic | -2.15 | *p=*4.02e-6 *** |
|  | Inoc (low) - Inorganic | -0.84 | *p=*0.039 * |
|  | Inoc (high) - Organic + Inorganic + Inoc (low) | -2.64 | *p=*1.15e-7 *** |
|  | Inoc (high) - Organic + Inorganic + Inoc (high) | -4.75 | *p=*2.74e-13 *** |
|  | Inoc (high) - Organic + Inorganic | -1.43 | *p=*8.39e-4 *** |
|  | Organic + Inorganic + Inoc (low) - Organic + Inorganic + Inoc (high) | -2.11 | *p=*5.31e-6 *** |
|  | Organic + Inorganic + Inoc (low) - Organic + Inorganic | 1.20 | *p=*0.004 ** |
|  | Organic + Inorganic + Inoc (low) - Organic | 3.16 | *p=*3.25e-9 *** |
|  | Organic + Inorganic + Inoc (low) - Inorganic | 2.51 | *p=*2.72e-7 *** |
|  | Organic + Inorganic + Inoc (high) - Organic + Inorganic | 3.31 | *p=*1.19e-9 *** |
|  | Organic + Inorganic + Inoc (high) - Organic | 5.27 | *p=*2.42e-14 *** |
|  | Organic + Inorganic + Inoc (high) - Inorganic | 4.62 | *p=*4.74e-13 *** |
|  | Organic + Inorganic - Organic | 1.95 | *p=*1.69e-5 *** |
|  | Organic + Inorganic - Inorganic | 1.31 | *p=*0.002 ** |
| Treatment * Site:  Contrasts between Kayonza - Ngoma | Organic + Inorganic + Inoc (low) | 1.19 | *p=*0.003 ** |
|  | Organic + Inorganic + Inoc (high) | 0.82 | *p=*0.038 * |
|  | Organic | -0.93 | *p=*0.019 * |
| **Season 2025B ANOVA results** | | | |
| **Fixed effect** | **F statistic** | **Degrees of freedom** | ***P* value** |
| Treatment | 43.40 | 7,32 | *p=*1.46e-14 *** |
| Site | 29.40 | 1,32 | *p=*5.82e-6 *** |
| Treatment*Site | 1.77 | 7,32 | *p=*0.13 n.s. |
| **Season 2025B post-hoc pairwise comparisons** | | | |
| **ANOVA fixed effect** | **Contrast** | **Estimate** | ***P* value** |
| Treatment | Untreated - Inoc (high) | -1.92 | *p=*4.74e-6 *** |
|  | Untreated - Organic + Inorganic + Inoc (low) | -3.69 | *p=*1.24e-11 *** |
|  | Untreated - Organic + Inorganic + Inoc (high) | -4.58 | *p=*1.18e-13 *** |
|  | Untreated - Organic + Inorganic | -2.81 | *p=*4.76e-9 *** |
|  | Untreated - Organic | -1.23 | *p=*0.001 ** |
|  | Untreated - Inorganic | -1.84 | *p=*7.98e-6 *** |
|  | Inoc (low) - Inoc (high) | -1.25 | *p=*0.001 ** |
|  | Inoc (low) - Organic + Inorganic + Inoc (low) | -3.03 | *p=*9.74e-10 *** |
|  | Inoc (low) - Organic + Inorganic + Inoc (high) | -3.92 | *p=*3.99e-12 *** |
|  | Inoc (low) - Organic + Inorganic | -2.14 | *p=*7.35e-7 *** |
|  | Inoc (low) - Organic | -0.57 | *p=*0.099 · |
|  | Inoc (low) - Inorganic | -1.18 | *p=*0.002 ** |
|  | Inoc (high) - Organic + Inorganic + Inoc (low) | -1.78 | *p=*1.22e-5 *** |
|  | Inoc (high) - Organic + Inorganic + Inoc (high) | -2.67 | *p=*1.11e-8 *** |
|  | Inoc (high) - Organic + Inorganic | -0.89 | *p=*0.014 * |
|  | Organic + Inorganic + Inoc (low) - Organic + Inorganic + Inoc (high) | -0.89 | *p=*0.014 * |
|  | Organic + Inorganic + Inoc (low) - Organic + Inorganic | 0.89 | *p=*0.014 * |
|  | Organic + Inorganic + Inoc (low) - Organic | 2.46 | *p=*5.27e-8 *** |
|  | Organic + Inorganic + Inoc (low) - Inorganic | 1.85 | *p=*7.64e-6 *** |
|  | Organic + Inorganic + Inoc (high) - Organic + Inorganic | 1.78 | *p=*1.22e-5 *** |
|  | Organic + Inorganic + Inoc (high) - Organic | 3.35 | *p=*1.07e-10 *** |
|  | Organic + Inorganic + Inoc (high) - Inorganic | 2.74 | *p=*6.73e-9 *** |
|  | Organic + Inorganic - Organic | 1.58 | *p=*6.71e-5 *** |
|  | Organic + Inorganic - Inorganic | 0.97 | *p=*0.009 ** |

***Supplementary Table 13.*** *Results of analyses of variance (ANOVAs) from linear models (~ Treatment * Site), and post-hoc pairwise comparisons run on maize plant biomass. Data were subset by Season prior to running models. P-values from post-hoc analyses are corrected for multiple comparisons; only marginally and statistically significant contrasts are shown. *p<0.05; **p<0.01; ***p<0.001; ·p<0.1; n.s. non-significant.*

| **Season 2025A ANOVA results** | | | |
| --- | --- | --- | --- |
| **Fixed effect** | ***F* statistic** | **Degrees of freedom** | ***P* value** |
| Treatment | 46.19 | 7,32 | *p=*5.98e-15 *** |
| Site | 182.97 | 1,32 | *p=*8.77e-15 *** |
| Treatment*Site | 1.77 | 7,32 | *p=*0.13 n.s. |
| **Season 2025A post-hoc pairwise comparisons** | | | |
| **ANOVA fixed effect** | **Contrast** | **Estimate** | ***P* value** |
| Treatment | Untreated - Inoc (low) | -0.53 | *p=*9.55e-6 *** |
|  | Untreated - Inoc (high) | -0.81 | *p=*5.92e-9 *** |
|  | Untreated - Organic + Inorganic + Inoc (low) | -1.23 | *p=*7.03e-13 *** |
|  | Untreated - Organic + Inorganic + Inoc (high) | -1.51 | *p=*5.12e-15 *** |
|  | Untreated - Organic + Inorganic | -1.07 | *p=*1.89e-11 *** |
|  | Untreated - Organic | -0.57 | *p=*3.53e-6 *** |
|  | Untreated - Inorganic | -0.86 | *p=*1.86e-9 *** |
|  | Inoc (low) - Inoc (high) | -0.28 | *p=*0.009 ** |
|  | Inoc (low) - Organic + Inorganic + Inoc (low) | -0.7 | *p=*1.18e-7 *** |
|  | Inoc (low) - Organic + Inorganic + Inoc (high) | -0.97 | *p=*1.40e-10 *** |
|  | Inoc (low) - Organic + Inorganic | -0.54 | *p=*9.31e-6 *** |
|  | Inoc (low) - Inorganic | -0.33 | *p=*0.003 ** |
|  | Inoc (high) - Organic + Inorganic + Inoc (low) | -0.42 | *p=*2.33e-4 *** |
|  | Inoc (high) - Organic + Inorganic + Inoc (high) | -0.695 | *p=*1.18e-7 *** |
|  | Inoc (high) - Organic + Inorganic | -0.26 | *p=*0.015 * |
|  | Inoc (high) - Organic | 0.24 | *p=*0.022 * |
|  | Organic + Inorganic + Inoc (low) - Organic + Inorganic + Inoc (high) | -0.28 | *p=*0.009 ** |
|  | Organic + Inorganic + Inoc (low) - Organic | 0.66 | *p=*3.08e-7 *** |
|  | Organic + Inorganic + Inoc (low) - Inorganic | 0.37 | *p=*9e-4 *** |
|  | Organic + Inorganic + Inoc (high) - Organic + Inorganic | 0.44 | *p=*1.37e-4 *** |
|  | Organic + Inorganic + Inoc (high) - Organic | 0.94 | *p=*2.98e-10 *** |
|  | Organic + Inorganic + Inoc (high) - Inorganic | 0.65 | *p=*3.95e-7 *** |
|  | Organic + Inorganic - Organic | 0.5 | *p=*2.55e-5 *** |
|  | Organic + Inorganic - Inorganic | 0.21 | *p=*0.044 * |
|  | Organic - Inorganic | -0.29 | *p=*0.008 ** |
| **Season 2025B ANOVA results** | | | |
| **Fixed effect** | **F statistic** | **Degrees of freedom** | ***P* value** |
| Treatment | 13.7701899 | 7,32 | *p=*4.53e-8 *** |
| Site | 1.95095537 | 1,32 | *p=*0.17 n.s. |
| Treatment*Site | 0.68942467 | 7,32 | *p=*0.68 n.s. |
| **Season 2025B post-hoc pairwise comparisons** | | | |
| **ANOVA fixed effect** | **Contrast** | **Estimate** | ***P* value** |
| Treatment | Untreated - Inoc (high) | -1.95 | *p=*1.2e-4 *** |
|  | Untreated - Organic + Inorganic + Inoc (low) | -2.5 | *p=*5.19e-6 *** |
|  | Untreated - Organic + Inorganic + Inoc (high) | -3.44 | *p=*2.37e-8 *** |
|  | Untreated - Organic + Inorganic | -2.15 | *p=*4.79e-5 *** |
|  | Untreated - Organic | -1.48 | *p=*0.002 ** |
|  | Untreated - Inorganic | -1.95 | *p=*1.2e-4 *** |
|  | Inoc (low) - Inoc (high) | -1.18 | *p=*0.011 * |
|  | Inoc (low) - Organic + Inorganic + Inoc (low) | -1.73 | *p=*4.93e-4 *** |
|  | Inoc (low) - Organic + Inorganic + Inoc (high) | -2.67 | *p=*2.25e-6 *** |
|  | Inoc (low) - Organic + Inorganic | -1.39 | *p=*0.004 ** |
|  | Inoc (low) - Inorganic | -1.19 | *p=*0.011 * |
|  | Inoc (high) - Organic + Inorganic + Inoc (high) | -1.49 | *p=*0.002 ** |
|  | Organic + Inorganic + Inoc (low) - Organic + Inorganic + Inoc (high) | -0.94 | *p=*0.042 * |
|  | Organic + Inorganic + Inoc (low) - Organic | 1.02 | *p=*0.029 * |
|  | Organic + Inorganic + Inoc (high) - Organic + Inorganic | 1.29 | *p=*0.007 ** |
|  | Organic + Inorganic + Inoc (high) - Organic | 1.96 | *p=*1.2e-4 *** |
|  | Organic + Inorganic + Inoc (high) - Inorganic | 1.49 | *p=*0.002 ** |

***Supplementary Table 14.*** *Results of analyses of variance (ANOVAs) from linear models (~ Treatment * Season), and post-hoc pairwise comparisons run on maize grain yield. Data were subset by Site prior to running models. P-values from post-hoc analyses are corrected for multiple comparisons; only marginally and statistically significant contrasts are shown. *p<0.05; **p<0.01; ***p<0.001; ·p<0.1; n.s. non-significant.*

| **Kayonza ANOVA results** | | | |
| --- | --- | --- | --- |
| **Fixed effect** | **F statistic** | **Degrees of freedom** | ***P* value** |
| Treatment | 111.45 | 7,32 | *p=*1.22e-20 *** |
| Season | 168.05 | 1,32 | *p=*2.79e-14 *** |
| Treatment*Season | 17.55 | 7,32 | *p=*2.56e-9 *** |
| **Kayonza post-hoc pairwise comparisons** | | | |
| **ANOVA fixed effect** | **Contrast** | **Estimate** | ***P* value** |
| Treatment * Season:  Contrasts within 2025A | Untreated - Inoc (low) | -1.72 | *p=*1.69e-4 *** |
|  | Untreated - Inoc (high) | -2.43 | *p=*1.28e-6 *** |
|  | Untreated - Organic + Inorganic + Inoc (low) | -6.84 | *p=*5.83e-17 *** |
|  | Untreated - Organic + Inorganic + Inoc (high) | -8.58 | *p=*2.96e-19 *** |
|  | Untreated - Organic + Inorganic | -4.76 | *p=*6.15e-13 *** |
|  | Untreated - Organic | -1.56 | *p=*4.85e-4 *** |
|  | Untreated - Inorganic | -3.29 | *p=*3.49e-9 *** |
|  | Inoc (low) - Organic + Inorganic + Inoc (low) | -5.12 | *p=*1.01e-13 *** |
|  | Inoc (low) - Organic + Inorganic + Inoc (high) | -6.85 | *p=*5.83e-17 *** |
|  | Inoc (low) - Organic + Inorganic | -3.04 | *p=*1.71e-8 *** |
|  | Inoc (low) - Inorganic | -1.57 | *p=*4.84e-4 *** |
|  | Inoc (high) - Organic + Inorganic + Inoc (low) | -4.42 | *p=*3.96e-12 *** |
|  | Inoc (high) - Organic + Inorganic + Inoc (high) | -6.15 | *p=*9.99e-16 *** |
|  | Inoc (high) - Organic + Inorganic | -2.34 | *p=*2.32e-6 *** |
|  | Inoc (high) - Organic | 0.86 | *p=*0.039 * |
|  | Inoc (high) - Inorganic | -0.87 | *p=*0.039 * |
|  | Organic + Inorganic + Inoc (low) - Organic + Inorganic + Inoc (high) | -1.73 | *p=*1.69e-4 *** |
|  | Organic + Inorganic + Inoc (low) - Organic + Inorganic | 2.08 | *p=*1.45e-5 *** |
|  | Organic + Inorganic + Inoc (low) - Organic | 5.28 | *p=*4.99e-14 *** |
|  | Organic + Inorganic + Inoc (low) - Inorganic | 3.55 | *p=*6.90e-10 *** |
|  | Organic + Inorganic + Inoc (high) - Organic + Inorganic | 3.81 | *p=*1.39e-10 *** |
|  | Organic + Inorganic + Inoc (high) - Organic | 7.01 | *p=*5.71e-17 *** |
|  | Organic + Inorganic + Inoc (high) - Inorganic | 5.28 | *p=*4.99e-14 *** |
|  | Organic + Inorganic - Organic | 3.2 | *p=*6.11e-9 *** |
|  | Organic + Inorganic - Inorganic | 1.47 | *p=*8.96e-4 *** |
|  | Organic - Inorganic | -1.73 | *p=*1.69e-4 *** |
| Treatment * Site:  Contrasts within 2025B | Untreated - Inoc (high) | -1.46 | *p=*0.002 ** |
|  | Untreated - Organic + Inorganic + Inoc (low) | -3.02 | *p=*9.67e-8 *** |
|  | Untreated - Organic + Inorganic + Inoc (high) | -3.71 | *p=*2.91e-9 *** |
|  | Untreated - Organic + Inorganic | -2.31 | *p=*7.95e-6 *** |
|  | Untreated - Organic | -0.90 | *p=*0.041 * |
|  | Untreated - Inorganic | -1.89 | *p=*1.08e-4 *** |
|  | Inoc (low) - Inoc (high) | -1.20 | *p=*0.008 ** |
|  | Inoc (low) - Organic + Inorganic + Inoc (low) | -2.77 | *p=*3.54e-7 *** |
|  | Inoc (low) - Organic + Inorganic + Inoc (high) | -3.46 | *p=*7.72e-9 *** |
|  | Inoc (low) - Organic + Inorganic | -2.05 | *p=*3.51e-5 *** |
|  | Inoc (low) - Inorganic | -1.63 | *p=*5.77e-4 *** |
|  | Inoc (high) - Organic + Inorganic + Inoc (low) | -1.56 | *p=*8.66e-4 *** |
|  | Inoc (high) - Organic + Inorganic + Inoc (high) | -2.25 | *p=*1.05e-5 *** |
|  | Organic + Inorganic + Inoc (low) - Organic | 2.12 | *p=*2.44e-5 *** |
|  | Organic + Inorganic + Inoc (low) - Inorganic | 1.14 | *p=*0.011 * |
|  | Organic + Inorganic + Inoc (high) - Organic + Inorganic | 1.40 | *p=*0.002 ** |
|  | Organic + Inorganic + Inoc (high) - Organic | 2.81 | *p=*3.25e-7 *** |
|  | Organic + Inorganic + Inoc (high) - Inorganic | 1.82 | *p=*1.54e-4 *** |
|  | Organic + Inorganic - Organic | 1.41 | *p=*0.002 ** |
|  | Organic - Inorganic | -0.98 | *p=*0.027 * |
| **Ngoma ANOVA results** | | | |
| **Fixed effect** | **F statistic** | **Degrees of freedom** | ***P* value** |
| Treatment | 87.51 | 7,32 | *p=*4.82e-19 *** |
| Season | 26.38 | 1,32 | *p=*1.34e-5 *** |
| Treatment*Season | 2.83 | 7,32 | *p=*0.021 * |
| **Ngoma post-hoc pairwise comparisons** | | | |
| **ANOVA fixed effect** | **Contrast** | **Estimate** | ***P* value** |
| Treatment * Season:  Contrasts within 2025A | Untreated - Inoc (low) | -2.69 | *p=*2.38e-6 *** |
|  | Untreated - Inoc (high) | -3.41 | *p=*4.00e-8 *** |
|  | Untreated - Organic + Inorganic + Inoc (low) | -6.05 | *p=*1.54e-13 *** |
|  | Untreated - Organic + Inorganic + Inoc (high) | -8.16 | *p=*6.35e-17 *** |
|  | Untreated - Organic + Inorganic | -4.84 | *p=*2.12e-11 *** |
|  | Untreated - Organic | -2.89 | *p=*7.13e-7 *** |
|  | Untreated - Inorganic | -3.53 | *p=*2.12e-8 *** |
|  | Inoc (low) - Organic + Inorganic + Inoc (low) | -3.36 | *p=*4.99e-8 *** |
|  | Inoc (low) - Organic + Inorganic + Inoc (high) | -5.47 | *p=*1.54e-12 *** |
|  | Inoc (low) - Organic + Inorganic | -2.15 | *p=*6.24e-5 *** |
|  | Inoc (high) - Organic + Inorganic + Inoc (low) | -2.64 | *p=*3.13e-6 *** |
|  | Inoc (high) - Organic + Inorganic + Inoc (high) | -4.75 | *p=*2.94e-11 *** |
|  | Inoc (high) - Organic + Inorganic | -1.43 | *p=*0.005 ** |
|  | Organic + Inorganic + Inoc (low) - Organic + Inorganic + Inoc (high) | -2.11 | *p=*7.78e-5 *** |
|  | Organic + Inorganic + Inoc (low) - Organic + Inorganic | 1.20 | *p=*0.015 * |
|  | Organic + Inorganic + Inoc (low) - Organic | 3.16 | *p=*1.44e-7 *** |
|  | Organic + Inorganic + Inoc (low) - Inorganic | 2.51 | *p=*6.50e-6 *** |
|  | Organic + Inorganic + Inoc (high) - Organic + Inorganic | 3.31 | *p=*5.92e-8 *** |
|  | Organic + Inorganic + Inoc (high) - Organic | 5.27 | *p=*3.08e-12 *** |
|  | Organic + Inorganic + Inoc (high) - Inorganic | 4.62 | *p=*4.85e-11 *** |
|  | Organic + Inorganic - Organic | 1.95 | *p=*2.0e-4 *** |
|  | Organic + Inorganic - Inorganic | 1.31 | *p=*0.009 ** |
| Treatment * Season:  Contrasts within 2025B | Untreated - Inoc (low) | -1.07 | *p=*0.031 * |
|  | Untreated - Inoc (high) | -2.38 | *p=*2.29e-5 *** |
|  | Untreated - Organic + Inorganic + Inoc (low) | -4.37 | *p=*4.55e-10 *** |
|  | Untreated - Organic + Inorganic + Inoc (high) | -5.46 | *p=*4.86e-12 *** |
|  | Untreated - Organic + Inorganic | -3.30 | *p=*1.03e-7 *** |
|  | Untreated - Organic | -1.56 | *p=*0.003 ** |
|  | Untreated - Inorganic | -1.79 | *p=*6.93e-4 *** |
|  | Inoc (low) - Inoc (high) | -1.31 | *p=*0.010 * |
|  | Inoc (low) - Organic + Inorganic + Inoc (low) | -3.30 | *p=*1.03e-7 *** |
|  | Inoc (low) - Organic + Inorganic + Inoc (high) | -4.39 | *p=*4.55e-10 *** |
|  | Inoc (low) - Organic + Inorganic | -2.23 | *p=*5.36e-5 *** |
|  | Inoc (high) - Organic + Inorganic + Inoc (low) | -1.99 | *p=*2.14e-4 *** |
|  | Inoc (high) - Organic + Inorganic + Inoc (high) | -3.08 | *p=*3.48e-7 *** |
|  | Organic + Inorganic + Inoc (low) - Organic + Inorganic + Inoc (high) | -1.09 | *p=*0.030 * |
|  | Organic + Inorganic + Inoc (low) - Organic + Inorganic | 1.07 | *p=*0.031 * |
|  | Organic + Inorganic + Inoc (low) - Organic | 2.81 | *p=*1.74e-6 *** |
|  | Organic + Inorganic + Inoc (low) - Inorganic | 2.57 | *p=*7.05e-6 *** |
|  | Organic + Inorganic + Inoc (high) - Organic + Inorganic | 2.15 | *p=*8.08e-5 *** |
|  | Organic + Inorganic + Inoc (high) - Organic | 3.90 | *p=*4.89e-9 *** |
|  | Organic + Inorganic + Inoc (high) - Inorganic | 3.66 | *p=*1.56e-8 *** |
|  | Organic + Inorganic - Organic | 1.74 | *p=*8.90e-4 *** |
|  | Organic + Inorganic - Inorganic | 1.51 | *p=*0.003 ** |

***Supplementary Table 15.*** *Results of analyses of variance (ANOVAs) from linear models (~ Treatment * Season), and post-hoc pairwise comparisons run on maize plant biomass. Data were subset by Site prior to running models. P-values from post-hoc analyses are corrected for multiple comparisons; only marginally and statistically significant contrasts are shown. *p<0.05; **p<0.01; ***p<0.001; ·p<0.1; n.s. non-significant.*

| **Kayonza ANOVA results** | | | |
| --- | --- | --- | --- |
| **Fixed effect** | ***F* statistic** | **Degrees of freedom** | ***P* value** |
| Treatment | 9.97 | 7,32 | *p=*1.51e-6 *** |
| Season | 169.64 | 1,32 | *p=*2.46e-14 *** |
| Treatment*Season | 1.23 | 7,32 | *p=*0.31 n.s. |
| **Kayonza post-hoc pairwise comparisons** | | | |
| **ANOVA fixed effect** | **Contrast** | **Estimate** | ***P* value** |
| Treatment | Untreated - Inoc (low) | -1.05 | *p=*0.01 * |
|  | Untreated - Inoc (high) | -1.61 | *p=*3.11e-4 *** |
|  | Untreated - Organic + Inorganic + Inoc (low) | -2.17 | *p=*7.45e-6 *** |
|  | Untreated - Organic + Inorganic + Inoc (high) | -2.50 | *p=*9.44e-7 *** |
|  | Untreated - Organic + Inorganic | -1.84 | *p=*7.37e-5 *** |
|  | Untreated - Organic | -1.13 | *p=*0.009 ** |
|  | Untreated - Inorganic | -1.67 | *p=*2.29e-4 *** |
|  | Inoc (low) - Organic + Inorganic + Inoc (low) | -1.11 | *p=*0.009 *** |
|  | Inoc (low) - Organic + Inorganic + Inoc (high) | -1.45 | *p=*9.81e-4 *** |
|  | Inoc (high) - Organic + Inorganic + Inoc (high) | -0.89 | *p=*0.035 * |
|  | Organic + Inorganic + Inoc (low) - Organic | 1.04 | *p=*0.014 * |
|  | Organic + Inorganic + Inoc (high) - Organic | 1.37 | *p=*0.002 ** |
|  | Organic + Inorganic + Inoc (high) - Inorganic | 0.83 | *p=*0.049 * |
| **Ngoma ANOVA results** | | | |
| **Fixed effect** | **F statistic** | **Degrees of freedom** | ***P* value** |
| Treatment | 23.26 | 7,32 | *p=*7.36e-11 *** |
| Season | 555.76 | 1,32 | *p=*8.56e-22 *** |
| Treatment*Season | 5.34 | 7,32 | *p=*4.04e-4 *** |
| **Ngoma post-hoc pairwise comparisons** | | | |
| **ANOVA fixed effect** | **Contrast** | **Estimate** | ***P* value** |
| Treatment * Season:  Contrasts within 2025A | Untreated - Organic + Inorganic + Inoc (low) | -1.11 | *p=*0.019 * |
|  | Untreated - Organic + Inorganic + Inoc (high) | -1.39 | *p=*0.003 ** |
|  | Inoc (low) - Organic + Inorganic + Inoc (high) | -1.02 | *p=*0.028 * |
|  | Organic + Inorganic + Inoc (high) - Organic | 0.91 | *p=*0.049 * |
| Treatment * Season:  Contrasts within 2025B | Untreated - Inoc (high) | -1.74 | *p=*1.25e-5 *** |
|  | Untreated - Organic + Inorganic + Inoc (low) | -2.01 | *p=*2.55e-6 *** |
|  | Untreated - Organic + Inorganic + Inoc (high) | -3.49 | *p=*5.09e-11 *** |
|  | Untreated - Organic + Inorganic | -1.89 | *p=*5.54e-6 *** |
|  | Untreated - Organic | -1.37 | *p=*2.43e-4 *** |
|  | Untreated - Inorganic | -1.69 | *p=*1.76e-5 *** |
|  | Inoc (low) - Inoc (high) | -1.63 | *p=*3.10e-5 *** |
|  | Inoc (low) - Organic + Inorganic + Inoc (low) | -1.90 | *p=*5.54e-6 *** |
|  | Inoc (low) - Organic + Inorganic + Inoc (high) | -3.38 | *p=*6.12e-11 *** |
|  | Inoc (low) - Organic + Inorganic | -1.77 | *p=*1.24e-5 *** |
|  | Inoc (low) - Organic | -1.25 | *p=*6.62e-4 *** |
|  | Inoc (low) - Inorganic | -1.58 | *p=*4.13e-5 *** |
|  | Inoc (high) - Organic + Inorganic + Inoc (high) | -1.75 | *p=*1.25e-5 *** |
|  | Organic + Inorganic + Inoc (low) - Organic + Inorganic + Inoc (high) | -1.48 | *p=*9.30e-5 *** |
|  | Organic + Inorganic + Inoc (high) - Organic + Inorganic | 1.61 | *p=*3.28e-5 *** |
|  | Organic + Inorganic + Inoc (high) - Organic | 2.13 | *p=*1.23e-6 *** |
|  | Organic + Inorganic + Inoc (high) - Inorganic | 1.8 | *p=*1.04e-5 *** |

***Supplementary Table 16.*** *Analysis of variance (ANOVA) from linear model (~ Treatment * Site), run on maize potato nutrient data (i.e., iron concentration). *p<0.05; **p<0.01; ***p<0.001; ·p<0.1; n.s. non-significant.*

| **ANOVA results** | | | |
| --- | --- | --- | --- |
| **Fixed effect** | ***F* statistic** | **Degrees of freedom** | ***P* value** |
| Treatment | 0.71 | 7,32 | *p=*0.66 n.s. |
| Site | 20.4 | 1,32 | *p=*8.04e-5 *** |
| Treatment*Site | 0.49 | 7,32 | *p=*0.83 n.s. |

***Supplementary Table 17.*** *Analyses of variance (ANOVAs) from linear models (~ Treatment * Site) and post-hoc pairwise comparisons run on maize nutrient data. P-values from post-hoc analyses are corrected for multiple comparisons; only marginally and statistically significant contrasts are shown. *p<0.05; **p<0.01; ***p<0.001; ·p<0.1; n.s. non-significant.*

| **Crude protein ANOVA results** | | | |
| --- | --- | --- | --- |
| **Fixed effect** | ***F* statistic** | **Degrees of freedom** | ***P* value** |
| Treatment | 3.35 | 7,32 | *p=*0.009 ** |
| Site | 0.52 | 1,32 | *p=*0.48 n.s. |
| Treatment*Site | 0.31 | 7,32 | *p=*0.94 n.s. |
| **Crude protein post-hoc pairwise comparisons** | | | |
| **ANOVA fixed effect** | **Contrast** | **Estimate** | ***P* value** |
| Treatment | Organic + Inorganic - Organic | 1.58 | *p=*0.032 * |
| **Potassium (K) ANOVA results** | | | |
| **Fixed effect** | ***F* statistic** | **Degrees of freedom** | ***P* value** |
| Treatment | 0.19 | 7,32 | *p=*0.99 n.s. |
| Site | 3.43 | 1,32 | *p=*0.073 · |
| Treatment*Site | 0.63 | 7,32 | *p=*0.73 n.s. |
| **Phosphorus (P) ANOVA results** | | | |
| **Fixed effect** | ***F* statistic** | **Degrees of freedom** | ***P* value** |
| Treatment | 0.73 | 7,32 | *p=*0.65 n.s. |
| Site | 7.78 | 1,32 | *p=*0.009 ** |
| Treatment*Site | 1.62 | 7,32 | *p=*0.17 n.s. |
| **Calcium (Ca) ANOVA results** | | | |
| **Fixed effect** | ***F* statistic** | **Degrees of freedom** | ***P* value** |
| Treatment | 0.65 | 7,32 | *p=*0.71 n.s. |
| Site | 12.3 | 1,32 | *p=*0.001 *** |
| Treatment*Site | 1.07 | 7,32 | *p=*0.41 n.s. |
| **Organic carbon (OC) ANOVA results** | | | |
| **Fixed effect** | ***F* statistic** | **Degrees of freedom** | ***P* value** |
| Treatment | 1.86 | 7,32 | *p=*0.11 n.s. |
| Site | 12.65 | 1,32 | *p=*0.001 ** |
| Treatment*Site | 0.64 | 7,32 | *p=*0.72 n.s. |
